# TBL38 is an atypical homogalacturonan acetylesterase with a peculiar cell wall microdomain localization in Arabidopsis seed mucilage secretory cells

**DOI:** 10.1101/2023.10.18.562901

**Authors:** Bastien G. Dauphin, David Ropartz, Philippe Ranocha, Maxime Rouffle, Camille Carton, Aurélie Le Ru, Yves Martinez, Isabelle Fourquaux, Simon Ollivier, Jessica Mac-Bear, Pauline Trezel, Audrey Geairon, Elisabeth Jamet, Christophe Dunand, Jérôme Pelloux, Marie-Christine Ralet, Vincent Burlat

## Abstract

Plant cell walls are made of complex polysaccharidic/proteinaceous network whose biosynthesis and dynamics implicate several cell compartments and impact plant development. The synthesis and remodeling of homogalacturonan pectins is associated with multiple developmental processes ranging from growth to response to biotic/abiotic stress. It encompasses Golgi-localized methylation and acetylation and subsequent demethylation and deacetylation in the cell wall. In the last decade, our comprehension of plant polysaccharides acetylation has increased significantly thanks to the study of the TRICHOME BIREFRINGENCE-LIKE (TBL) protein family. TBLs are mostly described as Golgi-localized acetyltransferases specifically targeting diverse hemicelluloses or pectins. Various *tbl* mutants showed altered wall mechanical properties and dynamics. Here, we study *TBL38* that is co-expressed with *PECTIN METHYLESTERASE INHIBITOR6* (*PMEI6*) and *PEROXIDASE 36* (*PRX36*) during the development of Arabidopsis seed mucilage secretory cells (MSCs). We demonstrate the atypical TBL38 cell wall localization restricted to the PMEI6/PRX36 MSC cell wall microdomain. A *tbl38* mutant displays an intriguing homogalacturonan immunological phenotype in this cell wall microdomain and in a MSC surface-enriched abrasion powder. This fraction was further characterized by mass spectrometry oligosaccharide profiling revealing an increased homogalacturonan acetylation phenotype. Finally, a recombinant TBL38 is shown to display pectin acetylesterase activity *in vitro*. These results indicate that TBL38 is an atypical cell wall-localized TBL that displays a homogalacturonan acetylesterase activity rather than a Golgi-localized acetyltransferase activity as observed in previously studied TBLs. TBL38 function during seed development is discussed.

## Introduction

Plant cells specifically design, synthesize and constantly remodel their cell wall (CW) to achieve proper cell shape and function along development to face environmental constraints. Such dynamics lead to specific composition and properties of CW polymers. Accumulating evidence of chemical group modifications on particular polysaccharides involved in key developmental processes can be found in the literature. The importance of pectin methylesterification dynamics has been largely studied, particularly for homogalacturonans (HGs). In the Golgi apparatus, HGs are methylated on the COOH of galacturonic acids (GalA) through methyl-transferase activity (Driouich et al., 2012) and later undergo additional remodeling once in the CW. Methylated HG can be demethylesterified by pectin methyl esterases (PMEs) whose activity can be regulated by PME inhibitors (PMEIs). The joint activity of both enzyme categories is the source of many developmental roles (Wolf et al., 2009). Additionally, *O*-acetylation of CW polymers such as pectins and hemicelluloses in the Golgi lumen has been described in multiple studies highlighting three groups of proteins involved in this process (Gille and Pauly, 2012). In *Arabidopsis thaliana*, (i) ALTERED XYLOGLUCAN 9 (AXY9) is encoded by a unique gene likely indirectly responsible for the non-specific acetylation of xyloglucan and xylan hemicelluloses (Schultink et al., 2015) leading to smaller plants when mutated. (ii) The REDUCED WALL *O*-ACETYLATION (RWA) family has four members in *A. thaliana* which also participate in the *O*-acetylation machinery of xylans, mannans, xyloglucans and pectins (Manabe et al., 2011; Manabe et al., 2013). (iii) The TRICHOME BIREFRINGENCE-LIKE (TBL) family comprises 46 members distributed in four phylogenetic clades in *A. thaliana* that target specific CW polysaccharides (**Figure 1; See Supplemental Table 1**). Originally named based on the observed changes in the birefringence properties of a mutant of *TRICHOME BIREFRINGENCE (TBR)* (Potikha and Delmer, 1995), 21 TBLs have been characterized for their *O*-acetyltransferase activity specific for the xyloglucan backbone and side chains, xylans, mannans, rhamnogalacturonan I (RG-I) or HGs (**Figure 1; Supplemental Table 1**, reviewed in (Gille and Pauly, 2012; Pauly and Ramírez, 2018). These activities are highly clade-specific since nine TBLs belonging to a similar clade displayed xylan-acetyltransferase activity (**Figure 1, Supplemental Table 1**).

**Figure 1:**
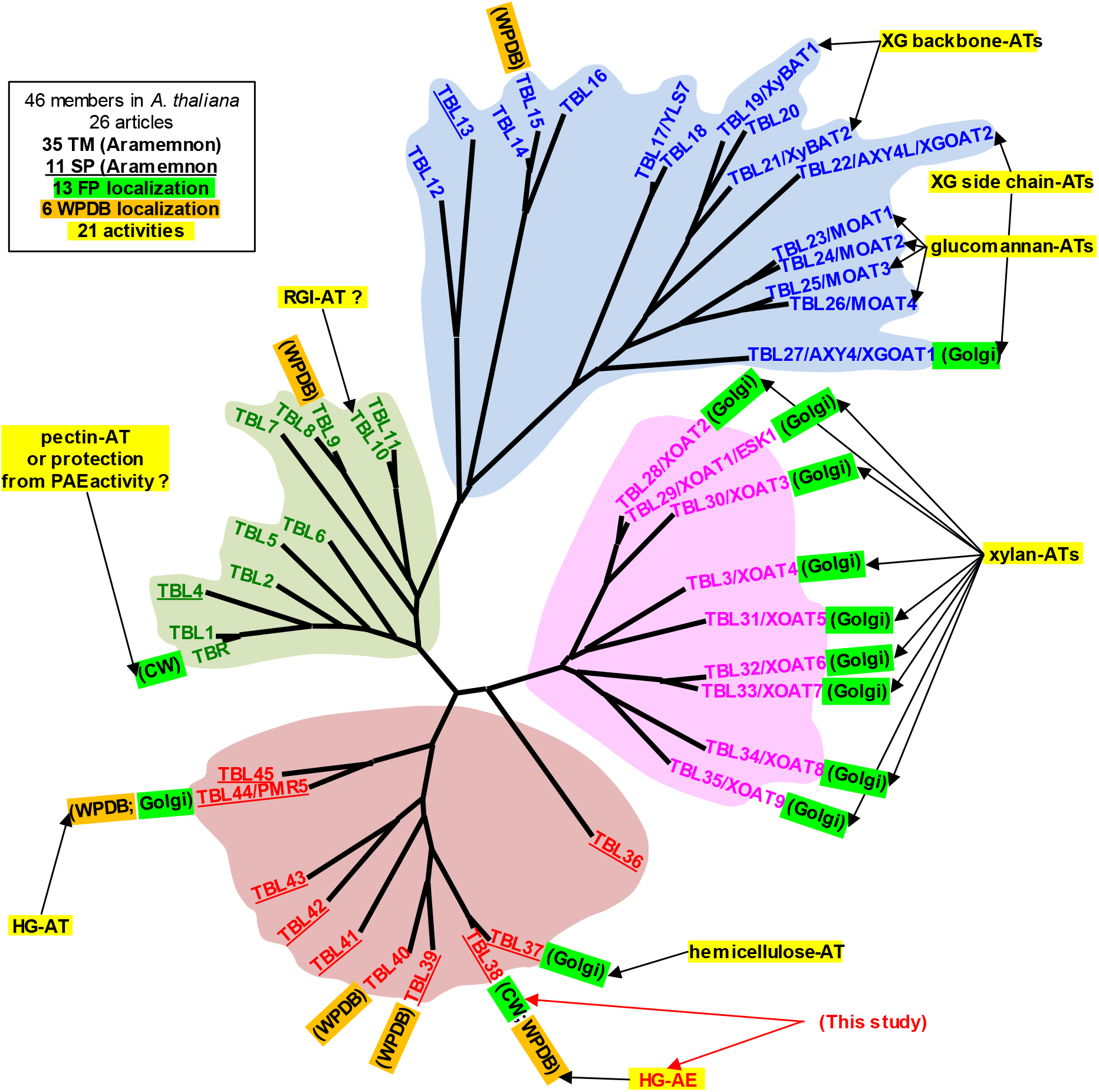
The TBL family mostly comprises Golgi-localized acetyltransferases specifically acting on various CW polymers. The 46 members show high sequence homology and two conserved GDSL and DXXH motifs associated with esterase activities. Previous work showed that the esterase activity isnot targeted on CW polymers but rather allows the formation of an acyl-enzyme intermediate, and described multiple TBLs as participating in *O*-acetylation of a wide range of CW polymers in the Golgi with the notable exception of TBR which was positioned to the CW. Twenty one recombinant enzymatic activities (yellow), 13 fluorescent protein (FP) localization (green) and 6 CW proteome occurrence https://www.polebio.lrsv.ups-tlse.fr/WallProtDB/ (orange) were reported. http://aramemnon.uni-koeln.de/ predicts that the N-terminus hydrophobic domains either acts as a transmembrane (TM) domain (35 non-underlined proteins) or as a cleaved signal peptide (11 underlined proteins). So far, each TBL or TBL phylogenetic cluster seems to specifically act on a CW polysaccharide which assumes various roles in development. The phylogenetic tree was adapted from (Bischoff et al., 2010). AE, acetylesterase; AT, acetyltransferase; PAE, pectin acetyl esterase; HG, homogalacturonan; RGI, rhamnogalacturonan I; XG, xyloglucan. See **Supplemental Table 1** for detailed information and references. This study reports on TBL38 peculiar localization and activity.

Mutations on particular *TBLs* led to (i) defect in CW structure and overall plant development as observed in *tbr, tbl3, tbl29* and *tbl37* (Bischoff et al., 2010; Lefebvre et al., 2011; Sun et al., 2020), and (ii) increased resistance to drought stress (Stranne et al., 2018) or pathogen infection (Chiniquy et al., 2019) in *tbl10* and *tbl44/pmr5*, respectively (**Supplemental Table 1**). Like AXY9, TBLs have the conserved GDSL domain which is typically associated with esterase activity/lipase (Akoh et al., 2004; Lai et al., 2017), and a DXXH motif also found in a rhamnogalacturonan acetylesterase of *Aspergillus aculeatus* (Molgaard and Larsen, 2004). The TBL acetylesterase activity allows the formation of an acyl-enzyme intermediate necessary for the *O*-acetyltransferase activity targeting various CW polymers (Gille and Pauly, 2012; Lunin et al., 2020) (**Figure 1, Supplemental Table 1**). Out of the 23 characterized TBLs, 12 have been localized to the Golgi alongside with AXY9 and RWAs, supporting their role as *O*-acetyltransferases (**Figure 1, Supplemental Table 1**). Surprisingly, TBR was localized to the CW (Sinclair et al., 2017). Yet, although its precise activity is still unknown, an overall decrease in pectin acetylation was observed in *tbr*, suggesting an acetyltransferase activity that is surprising considering that the existence of a CW-localized acetyl donor is unclear (Sinclair et al., 2017). *In silico* prediction of the targeting function of the N-terminal hydrophobic sequence sorted the 46 TBLs in two groups: 35 with a transmembrane anchoring domain, and 11 with a cleavable signal peptide. Six TBLs have been identified in CW-enriched fractions without clear relationship with the *in silico* targeting predictions (**Supplemental Table 1**). Overall, experimental evidence depicts TBLs as acetylesterases forming an acyl-enzyme intermediate to further perform an *O*-acetyltransferase activity targeted on various CW polymers during their synthesis in the Golgi lumen.

Initially, we screened genes involved in *A. thaliana* seed mucilage release with particular interest on candidates that could modify the esterification status of HGs. The mucilage is a polysaccharidic hydrogel synthesized by the seed mucilage secretory cells (MSCs) in the outer most cell layer of the seed coat of the so-called myxospermous species (Francoz et al., 2015; Viudes et al., 2021; Western, 2012). During *A. thaliana* seed maturation, the HG methylesterification status is modified through the PECTIN METHYLESTERASE INHIBITOR6 6 (PMEI6) activity on an unknown PME in a CW microdomain at the top of the radial primary wall of MSCs, leading to a HG partially methylesterified pattern (Francoz et al., 2019b; Saez-Aguayo et al., 2013). This pattern acts as an anchoring platform for PEROXIDASE36 (PRX36) which then induces a local weakening and thinning of the CW microdomain (Francoz et al., 2019b; Kunieda et al., 2013). After seed inhibition, the swelling mucilage breaks the pre-weakened CW microdomain and is extruded around the seed. We identified *TBL38* (*At1g29050*) for its high co-expression level with *PRX36* and *PMEI6* during *A. thaliana* seed coat development. Based on the literature cited above and considering its co-expression with *PMEI6*, we first hypothesized TBL38 to be a Golgi localized protein acting as an acetyltransferase onto CW polymers possibly including HGs. Hereafter, we examined TBL38 localization, activity and putative role in MSCs and uncovered that TBL38 is an atypical CW-localized HG acetylesterase rather than a Golgi-localized acetyltransferase. Although TBLs are not thought to influence the methylesterification status of CW polymers, we investigated the putative indirect role of TBL38-dependent HG *O*-deacetylation on the HG methylation status and on PRX36 anchoring, thus adding another layer of complexity on the fine tuning of HG remodeling during plant development.

## Results

### *TBL38* is co-expressed with *PRX36* and *PMEI6* in mucilage secretory cells at intermediate developmental stages of seed development

Using the tissue-specific seed developmental kinetics GSE12404 dataset (Belmonte et al., 2013), we previously built a co-expression network centered on *PRX36* in which *PMEI6* came as the second hit (Francoz et al., 2019b) (**Supplemental Table 2**). We noticed the high co-expression of *TBL38* which was ranked 13 in this network with a strong specific seed coat expression at the linear cotyledon stage (**Figure 2A-B; Supplemental Table 2**). We filtered *TBL* expression data which was available for 34 out of the 46 *TBLs*, although only 23 were above the detection limit, and confirmed that *TBL38* was likely to be the only *TBL* highly co-expressed with *PRX36* and *PMEI6* (**Supplemental Table 2**). The seed coat transcriptomic data encompasses five cell layers: ii1 (inner integument), ii1’, ii2, oi1 (outer integument), and oi2, commonly addressed here as MSCs (Debeaujon et al., 2003; Francoz et al., 2015).

**Figure 2:**
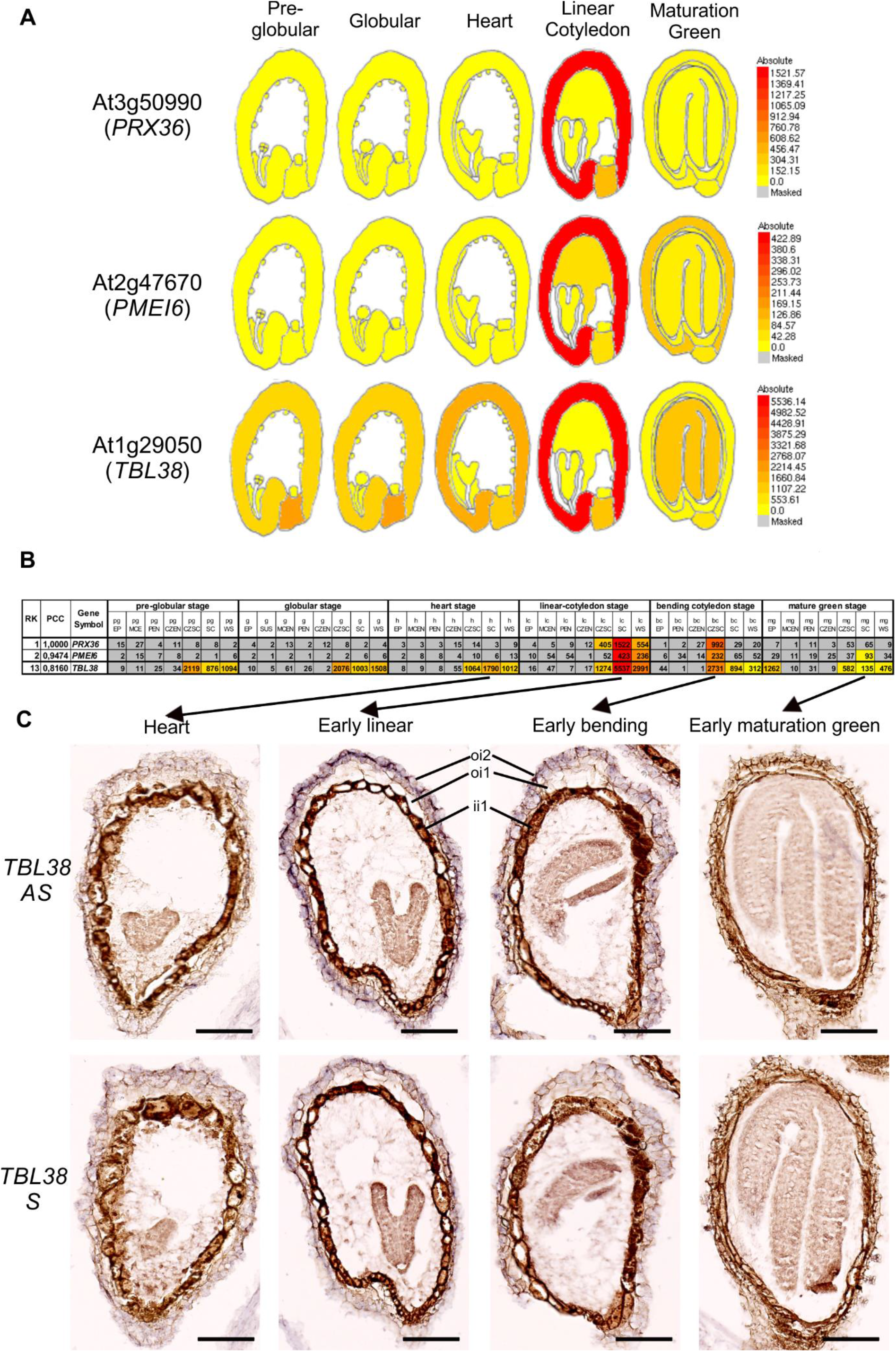
*TBL38* is co-expressed with *PRX36 and PMEI6* in the seed coat and transcript localization of *TBL38* is tied to the mucilage secretory cells (outer integument 2). **(A)** eFP browser pictogram displaying the similarity in the spatiotemporal expression of *PRX36*, *PMEI6* and *TBL38* in *A. thaliana* seed coat. Absolute tissue-specific expression level is represented in color variation from yellow (lowest) to red (highest). Adapted from http://bar.utoronto.ca/efp/cgi-bin/efpWeb.cgi. **(B)** *PRX36* co-expression network using the same dataset highlighting the similar profiles of *PRX36*, *PMEI6* and *TBL38* (see **Supplemental Table 2** for details). **(C)** *In situ* hybridization of dig-labeled RNA probes targeting *TBL38* on paraffin sections of developing wild-type seeds. Anti-sense probes showed highest purple signal in the outermost layer corresponding to MSCs between the early linear and early bending developmental stages. These signals should be subtracted from the overall background obtained on serial sections of the same seeds with the sense probe control and natural pigmentation. Bars: 200 µm.

Therefore, we determined MSC-specific *TBL38* expression using *in situ* hybridization on tissue array paraffin sections encompassing the whole developmental kinetics of wild-type seeds. *TBL38* anti-sense probes displayed a clear purple/violet hybridization signal in MSCs, particularly between the early linear and early bending cotyledon developmental stages (**Figure 2C**). No such signal could be observed at earlier and later stages with the anti-sense probe and at all developmental stages with the sense probe used as negative control to account for the overall non-specific background and natural pigmentation. We could associate the specific *TBL38* anti-sense probe signal to MSCs excluding the chalazal tissues, in agreement with -and refining-the transcriptomic data (**Figure 2A-C**).

### TBL38-TagRFP is transiently localized to the PRX36-PMEI6 outer periclinal/radial cell wall microdomain

We generated a *p35S::TBL38-TagRFP* construct to transiently transform *N. benthamiana* leaves. The TBL38-TagRFP fluorescence did not colocalize with a Golgi-YFP marker, but rather displayed a cellular delineation (**Figure 3A**). Under plasmolysis conditions, we could clearly attribute the TBL38-TagRFP signal to the CW separating from the receding plasma membrane YFP marker signals (**Figure 3A**). We concluded that TBL38 is a CW-localized protein similarly to TBR and contrarily to the 12 other Golgi-localized TBLs (Sinclair et al., 2017) (**Figure 1, Supplemental Table 1**). We then moved on to assess protein localization in *A. thaliana* seed MSCs. As seed development advances, MSCs become mostly made of different CWs, namely the primary CW, the mucilage and the columella (Francoz et al., 2015). Within MSCs, increasing evidence have positioned CW proteins in only one or two types of CW (Francoz et al., 2019b; McGee et al., 2019; Sola et al., 2019) or in sub-layer (microdomain) of one CW (Dauphin et al., 2022; Francoz et al., 2019b).

**Figure 3:**
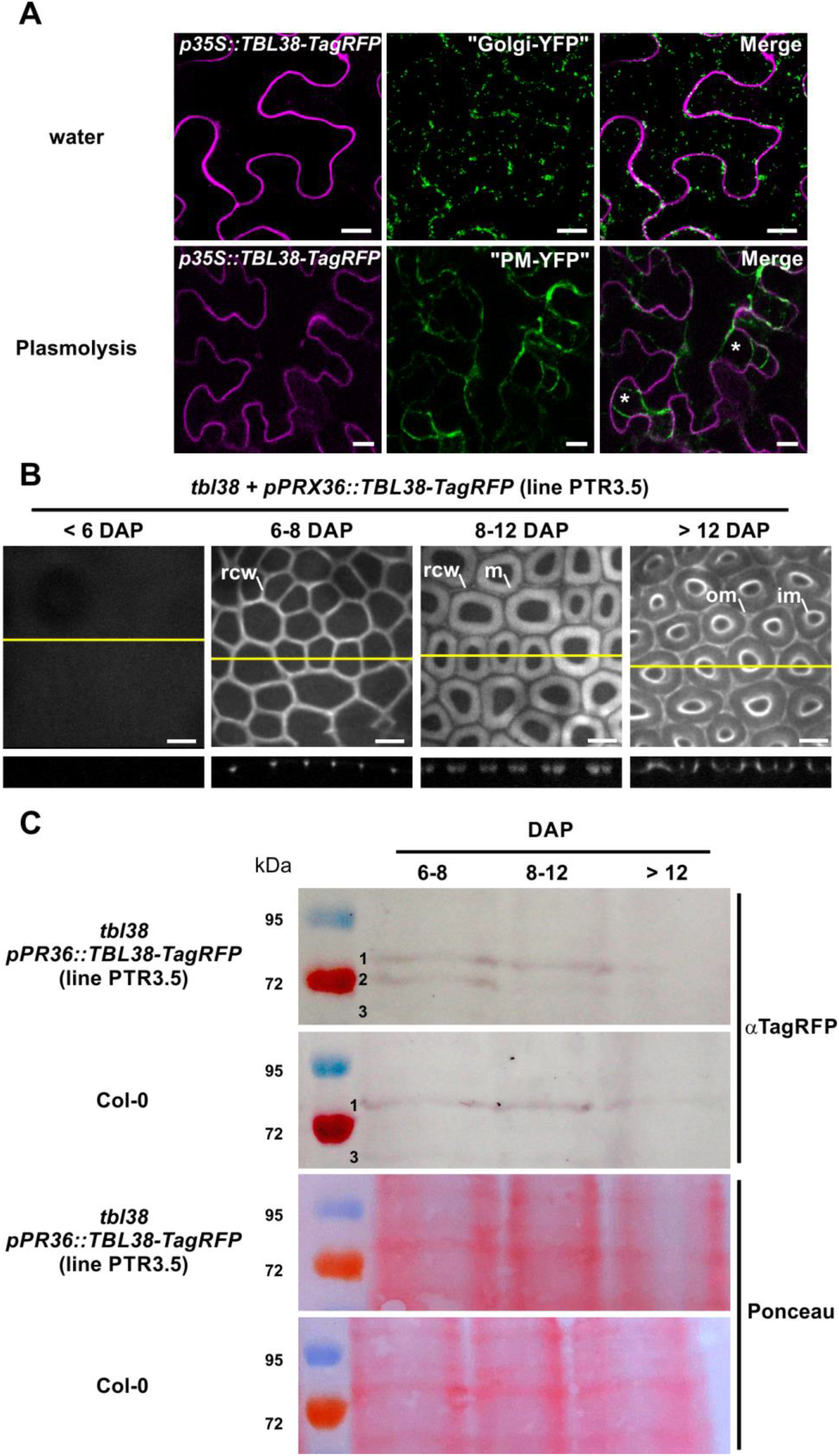
TBL38 is a cell wall protein transiently localized to the radial primary CW microdomain of MSCs. **(A)** Confocal microscopy visualization of both TBL38-TagRFP and a Golgi-YFP marker transiently overexpressed in *N. benthamiana* leaves indicates that TBL38 is not a Golgi-localized protein. Under plasmolysis, the TBL38-TagRFP signal and the plasma membrane-YFP marker signal are separated, indicating the localization of TBL38-TagRFP to the CW. First row images are extracted from a maximum projection of 17 stacks. Second row images are a representative view of a single confocal plan. Bars: 10 µm. **(B)** Confocal spinning disk observations of TBL38-TagRFP in *A. thaliana* developing seeds. At the onset of *pPRX36* activity (Francoz et al., 2019b), TBL38-TagRFP fluorescence was positioned to the radial primary CW of MSCs. Orthogonal projection refined this localization to the CW microdomain consisting in the top of the radial primary CW similarly to PRX36-TagRFP (Francoz et al., 2019b). However, contrary to the previously observed stable localization of PRX36-TagRFP along seed development, TBL38-TagRFP fluorescence was delocalized to the future mucilage pocket at 8-12 DAP and eventually to the outer and inner margins of the mucilage-filled pocket at > 12 DAP. Sum intensity projections were done using similarly sized projection of 30 Z stacks with no singular edition. rcw, radial CW microdomain; m, mucilage pocket; im and om, inner and outer margins of mucilage pocket, respectively. Bars: 25 µm. (see **Supplemental Figure 1** for full kinetics performed on three individual transformed lines). **(C)** αTag-RFP western blot performed using total protein extracts from developing seeds harvested at stages corresponding to the three sequential patterns of fluorescence shows that the band #2 corresponding to the TBL38-TagRFP fusion protein is only seen at the early stage when TBL38-TagRFP is localized to the CW microdomain. Non-specific bands #1 and #3 are also present in Col-0 samples (see **Supplemental Figure 2** for full membrane visualization).

To localize TBL38 in *A. thaliana* MSCs, we generated a *pPRX36:*TBL38-TagRFP construct which was stably transformed in *tbl38* plants. We localized TBL38-TagRFP to the radial primary CW of MSCs at the onset of *pPRX36* activity at early developmental stages (**Figure 3B; Supplemental Figure 1**). Orthogonal projection allowed to refine TBL38-positonning at the top of the radial primary CW, corresponding to the CW microdomain encompassing PRX36 (Francoz et al., 2019b). Intriguingly, while PRX36-TagRFP was stably accumulated at this location using the same promoter (Francoz et al., 2019b), TBL38-TagRFP appeared to be naturally delocalized to the future mucilage pocket and eventually pushed toward the inner and outer margins of the pocket when filled with mucilage after 12 DAP **(Figure 3B; Supplemental Figure 1)**.

We accurately sampled developing seeds at stages corresponding to the three above mentioned fluorescence patterns and performed a αTagRFP western blot (**Figure 3C; Supplemental Figure 2**). Excluding the non-specific bands (#1 and 3) also observed with similarly staged Col-0 seeds, a single specific band (#2) corresponding to the apparent molecular mass of TBL38-TagRFP fusion protein (70 kDa) was only observed at 6-8 DAP when TBL38-TagRFP was localized to the CW microdomain. The absence of such specific band when the fluorescence was later localized to the mucilage pocket suggests that this additional fluorescence pattern rather corresponds to the delocalized TagRFP possibly cleaved from TBL38. This hypothesis is strengthened by the appearance of a 28 kDa band that may correspond to cleaved TagRFP at 8-12 DAP that unfortunately migrated with an additional non-specific band of similar molecular mass also observed in Col-0 after 12 DAP (**Supplemental Figure 2**).

Altogether, expression of TBL38-TagRFP in both *N. benthamiana* and *A. thaliana* showed an unexpected CW localization making TBL38 the second TBL localized to the CW with TBR. Because its localization to the outer radial CW microdomain was strikingly similar to that of PRX36, we investigated TBL38 involvement in the above mentioned PRX36-PMEI6 molecular module and particularly its influence on HG modification.

### The mucilage release and the anchoring of PRX36 are not significantly modified in *tbl38*

Mutation in either *PRX36* or *PMEI6* results in impaired mucilage release because the primary CW microdomain improperly brakes up upon seed imbibition (Francoz et al., 2019b; Kunieda et al., 2013; Saez-Aguayo et al., 2013). Because *TBL38* and *PRX36* gene products have similar spatio-temporal expression and positioning, we assessed whether mucilage release was affected in mutated seeds. We isolated a homozygous KO line for *tbl38* (**Supplemental Figure 3**) and used it for comparison with Col-0 wild type seeds using ruthenium red staining of adherent mucilage. Visually, we did not see any differences between WT and *tbl38* seeds (**Supplemental Figure 4A**), contrarily to impaired mucilage previously observed in *prx36* or *pmei6* (Francoz et al., 2019b; Kunieda et al., 2013; Saez-Aguayo et al., 2013). Quantification of adherent mucilage area and circularity on > 1,000 seeds confirmed the similarity of the patterns of Col-0 and *tbl38* (**Supplemental Figure 4B**). We did not see any obvious change in mucilage staining intensity as sometimes observed in mutated background responsible for mucilage modification (Turbant et al., 2016).

Since PRX36 anchoring to a specific HG platform and subsequent CW weakening are required for proper mucilage release, we investigated PRX36 positioning in *tbl38.* For this, we used the αPRX36 antibody on paraffin sections of developing seeds from Col-0 or *tbl38*. Anti-PRX36 similarly labeled the top of the radial primary CW in Col-0 and *tbl38* at all developmental stages (**Supplemental Figure 4C**). *tbl38* did not show the loss of signal previously observed in *prx36* or *pmei6* (Francoz et al., 2019b). To further validate this result, we generated a *pPRX36::PRX36-TagRFP* construct, which we expressed in *prx36* and *tbl38* backgrounds. Observation of the PRX36-TagRFP signal was made on developing seeds aged from 6-to-12 DAP, a period which encompasses the onset of the *PRX36* promoter activity up to the last stage of the clearly visible PRX36-TagRFP fluorescence signal (Francoz et al., 2019b). PRX36-TagRFP showed very similar signal localization at the top of the radial primary CW in both *tbl38* and *prx36* (**Supplemental Figure 4D**). Again, no delocalization of the signal was observed in *tbl38* contrary to the PRX36-TagRFP delocalization in the mucilage pocket previously observed in *pmei6* (Francoz et al., 2019b).

So far, the particular PMEI6-specific HG methylation pattern that exists at the top of the radial primary CW linked proper mucilage release with PRX36 anchoring (Francoz et al., 2019b). Our results did not show such a relationship between TBL38, mucilage release and PRX36 positioning, suggesting that no changes in the HG methylation pattern occur in *tbl38*. However, TBL38 could be involved in acetylation of CW polymers similarly to other described TBLs. Because changes in HG acetylation could lead to different HG methylesterification status, possibly through indirect modulation of PME activity or hindering the dimerization of pectin chains (Kohn and Furda, 1968; Ralet et al., 2003), we first studied the HG methylation pattern in *tbl38*.

### The partially methylated homogalacturonan JIM7 epitope corresponding to the PRX36-PMEI6 CW microdomain is maintained in *tbl38*, while the supposed similar LM20 epitope is surprisingly lost

The specific HG platform enabling PRX36 anchoring, is generated through the inhibition of an unknown PME by PMEI6 leading to a partially methylesterified pattern in HGs at the top of the radial primary CW. This molecular pattern could be labeled by both JIM7 and LM20, two monoclonal antibodies routinely used to characterize partially methylesterified HGs, and these labelings were lost in *pmei6* (Francoz et al., 2019b). Thus, we incubated JIM7 and LM20 on paraffin sections of developing seeds from *tbl38* and Col-0.

As a control, JIM7 signals were correctly positioned at the top of the radial primary CW in Col-0 developmental kinetics, particularly as MSC development advanced to later stages (**Figure 4A**). Very similar JIM7 labeling was observed in *tbl38* (**Figure 4A**) contrary to the loss of signal previously observed in *pmei6* (Francoz et al., 2019b). A similar pattern was observed with LM20 along the developmental kinetics of the Col-0 control genotype (**Figure 4B**). However, the LM20 labeling was intriguingly lost in *tbl38* (**Figure 4B**). This was related to the *tbl38* mutation since LM20 labeling was restored along the developmental kinetics of *tbl38* plants complemented with a *pPRX36::TBL38-TagRFP* construct (**Figure 4B**). While both antibodies are generally used in parallel to similarly label partially methylesterified HG, their partially characterized epitopes may be somehow different (Clausen et al., 2003; Verhertbruggen et al., 2009). Overall, we concluded that *tbl38* mutation resulted in lack of the LM20 signal, but not of that of JIM7. Yet, it was still unclear if this could be due to putative changes in the acetylation of HGs which may prevent LM20 binding through steric hindrance, or to changes in HG methylation as indirect effect of acetylation. Regardless, we sought to investigate TBL38 biochemical function in MSCs.

**Figure 4:**
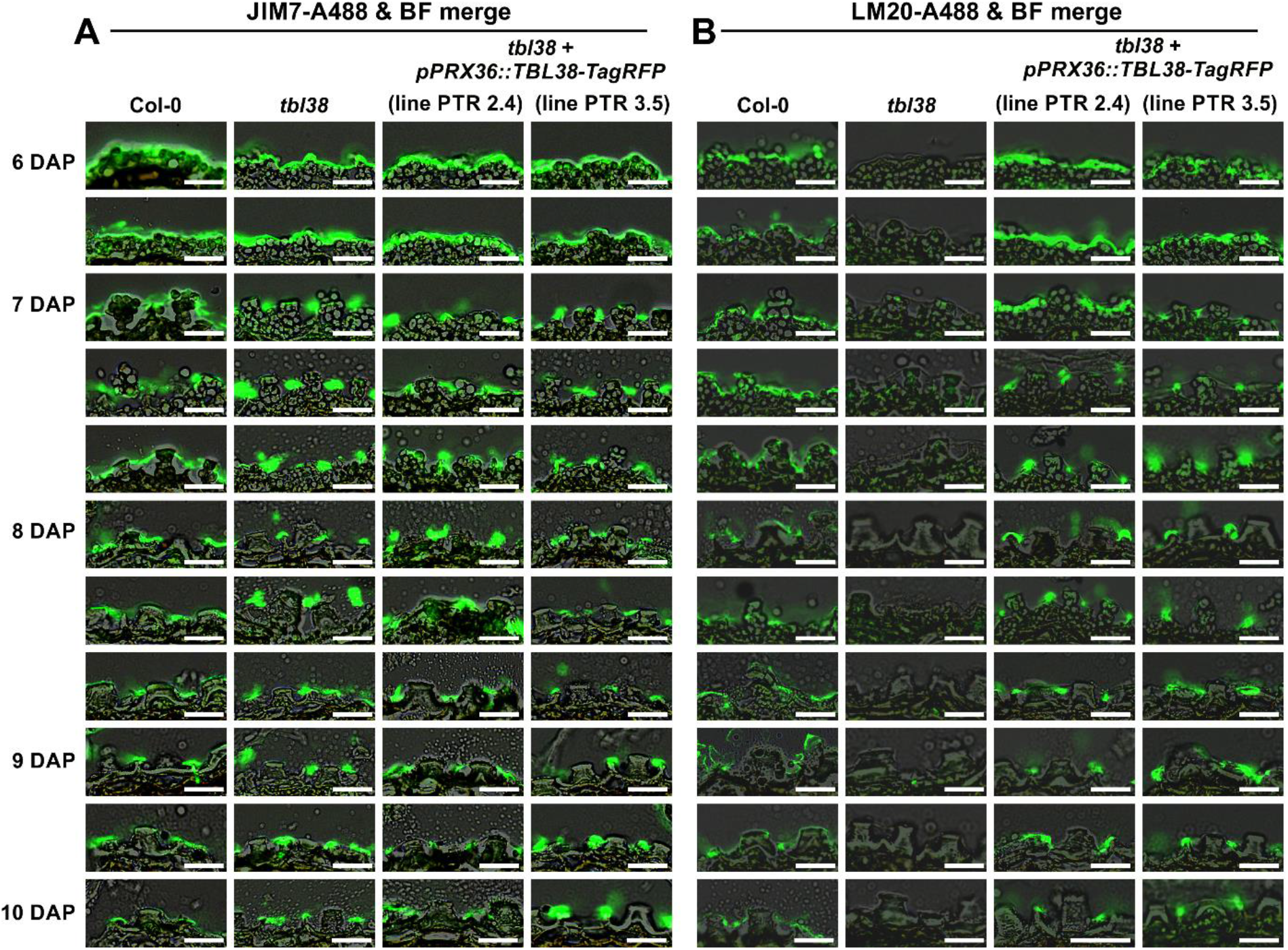
LM20 but not JIM7 labeling is lost in *tbl38*. **(A)** In Col-0, JIM7 labeling of partially methylesterified HG is visible as early as 6 DAP at the MSC surface and is restricted to the top of the radial primary CW microdomain of more mature MSCs (7-10 DAP). A very similar labeling pattern occurs in *tbl38* as well as in two complemented lines (*tbl38* transformed with *pPRX36::TBL38-TagRFP*). **(B)** LM20 was previously thought to bind to similar epitopes as JIM7 in MSCs since both epitopes were lost in *pmei6* (Francoz et al., 2019b). However, contrary to JIM7, the LM20 labeling at the top of the radial primary CW microdomain was lost in *tbl38* and restored in the two tested complemented lines. 40 x scans of fluorescence and brightfield channels were merged. The same parameters were used to analyze each image which are a representative of n > 20 MSCs at each developmental stage. Additional stages positioned between the staged ranks are shown. Bars: 25µm.

### TBL38 displays an atypical acetylesterase activity towards CW polymers

We took advantage of MSC surface accessibility to generate a MSC surface-enriched CW abrasion powder from dry seeds using home-made abrasion columns (**see Methods**). The resulting homogeneous powder (about 1 mg per 50 mg seeds) was collected through a nylon mesh and subsequently analyzed (**Supplemental Figure 5A-F**). We confirmed that the abrasion process was restricted to the MSC surface as MSC CW autofluorescence was strongly reduced but could still be barely observed in abraded seeds (**Supplemental Figure 5G-H**) which released residual amount of adherent mucilage when imbibed (**Supplemental Figure 5I-J**). Although the overall MSC surface morphology was affected, we could still see the base of broken columella and radial CW (**Supplemental Figure 5M-N**). We successfully recovered a surface-abraded powder containing the top of the radial primary wall, considering the loss of LM20 signal in Col-0 abraded seeds (**Supplemental Figure 5K-L**). We validated the method since JIM7 and LM20 dot blot signal intensities in *pmei6* were both about 5 % as compared to Col-0 (**Figure 5A, B**), in agreement with the previously observed loss of JIM7 and LM20 immunofluorescence signals in development kinetics of *pmei6* MSCs (Francoz et al., 2019b). Again, in agreement with the present immunofluorescence study on development kinetics (**Figure 4**), the JIM7 signal intensity from dry seed surface abraded fractions was very similar in all genotypes except *pmei6*, since *tbl38* and three complemented lines all showed 91-95 % of Col-0 intensity (**Figure 5A**). However, the LM20 signal intensity in *tbl38* was only 4 % of that in Col-0 control while the three complemented lines showed a strong -though partial-signal restoration with 66-79 % of Col-0 intensity (**Figure 5B**).

**Figure 5:**
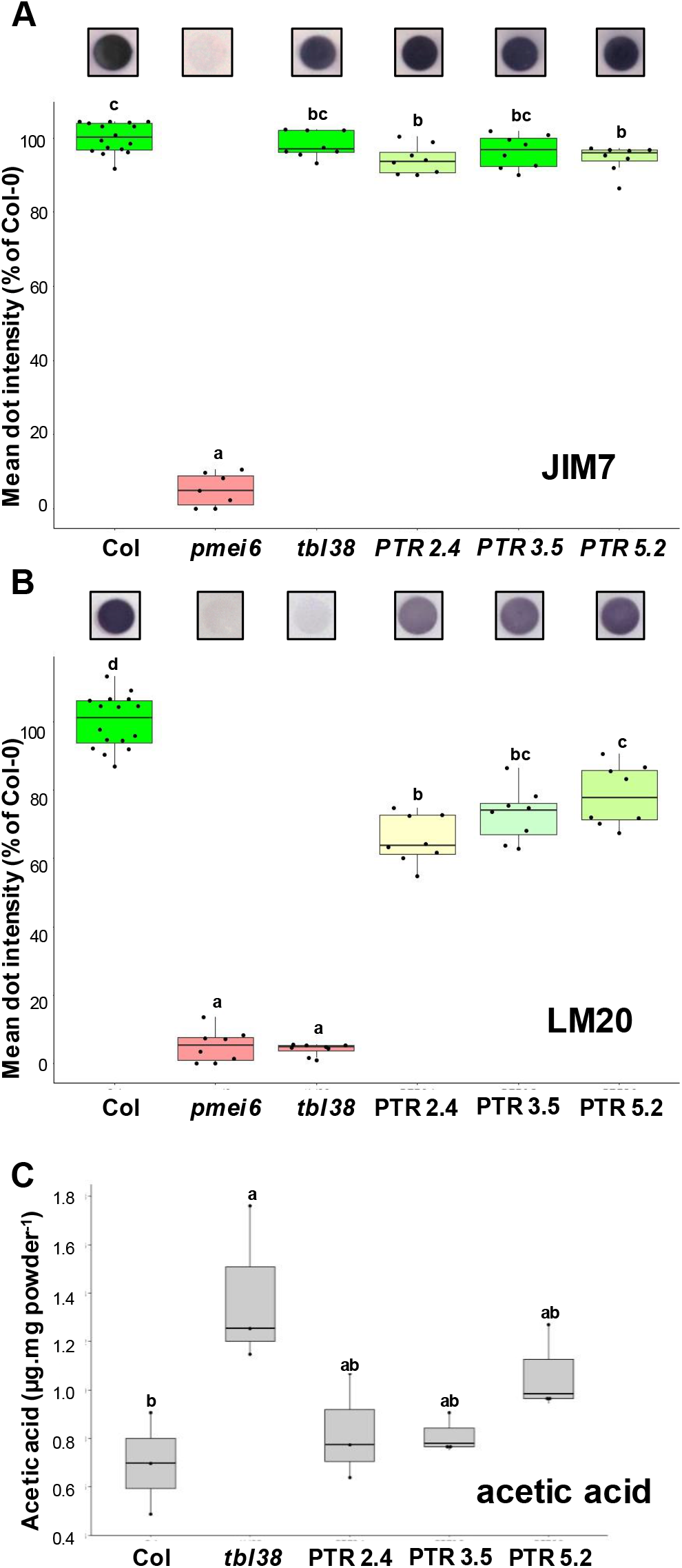
Dry seed surface enriched fraction shows a decreased LM20 epitope signal intensity and an increased acetic acid content in *tbl38* compared to Col-0, and the wild type phenotype is restored in complemented lines. JIM7. **(A)** and LM20 **(B)** immuno dot blot of dry seed surface CW-enriched fraction from Col-0, *pmei6*, *tbl38* and three *tbl38* complemented lines expressing *pPRX36*:TBL38-TagRFP (PTR2.4, 3.5 and 5.2). On top of the panels are shown screen shots of individual dots from the same nitrocellulose membrane. On the bottom of the panel are shown box plot representations of the data analysis. **(C)** Box plot representation of the acetic acid content released following acid hydrolysis of dry seed surface CW enriched fraction from Col-0, *tbl38* and three *tbl38* complemented lines expressing *pPRX36::TBL38-TagRFP* (PTR2.4, 3.5 and 5.2). Three biological replicates each with at least two technical replicates were analyzed. Letters above the bars show the statistical distribution of the results using ANOVA and TUKEY HSD tests.

So far, all the described TBLs were either directly characterized as acetyltransferases or associated with lower acetylation content of CW polymers in mutated backgrounds (**Supplemental Table 1**). Therefore, we quantified the acetylation of CW polymers using the surface-enriched CW abrasion powders. Surprisingly, the mean amount of released acetic acid in *tbl38* was 199 % of that in Col-0 and the level dropped back to 117-153 % of that of Col-0 in the three complemented lines (**Figure 5C**). This suggested that TBL38 activity was akin to an acetylesterase rather than the previously described acetyltransferase of other characterized TBLs. Because the increased acetylation level in *tbl38* could be achieved through acetylation of multiple polymers, we sought to identify TBL38 substrates in these fractions.

### TBL38 is an homogalacturonan acetylesterase

Surface-enriched CW abrasion powders were enzymatically hydrolyzed with a pectin lyase (PL) or a polygalacturonase (PG), displaying specificity toward highly and lowly methylesterified HGs, respectively. The resulting oligogalacturonates (OGAs) from Col-0, *tbl38* as well as *pmei6* and the complemented line PTR3.5 used as additional controls, have then been analyzed by mass spectrometry, which allowed to determine the degree of polymerization (DP), degree of methylation (DM) and degree of acetylation (DA). Technical repeats showed the low standard deviation of the intensities of identified species validating the approach despite the low amount of biological material **(Supplemental Table 3A)**. We first used a PL having a high affinity for HG with a high DM. The intensity of the identified OGA species was summed for the identified OGA species of same DA and DP regardless the DM, or same DM and DP regardless the DA (**Supplemental Table 3B**) and for each genotype, the results were plotted as DA vs DP (**Figure 6A-D**), or DM vs DP (**Figure 6E-H**). We observed a clear qualitative HG acetylation phenotype in *tbl38* as compared to Col-0 and *pmei6* (**Figure 6A-C**) and a qualitative complementation of PTR3.5 (**Figure 6D**). This was in agreement and refined the high acetylation phenotype observed in *tbl38* with the enzymatic assay (**Figure 5C**) since new OGA species of higher DA appeared in *tbl38* as compared to the three other genotypes. This strongly suggests that TBL38 could be an HG acetylesterase. Conversely, a quantitative HG methylation phenotype was observed for *pmei6* as compared to Col-0, *tbl38* and PTR3.5 (**Figure 6E-H**).

**Figure 6:**
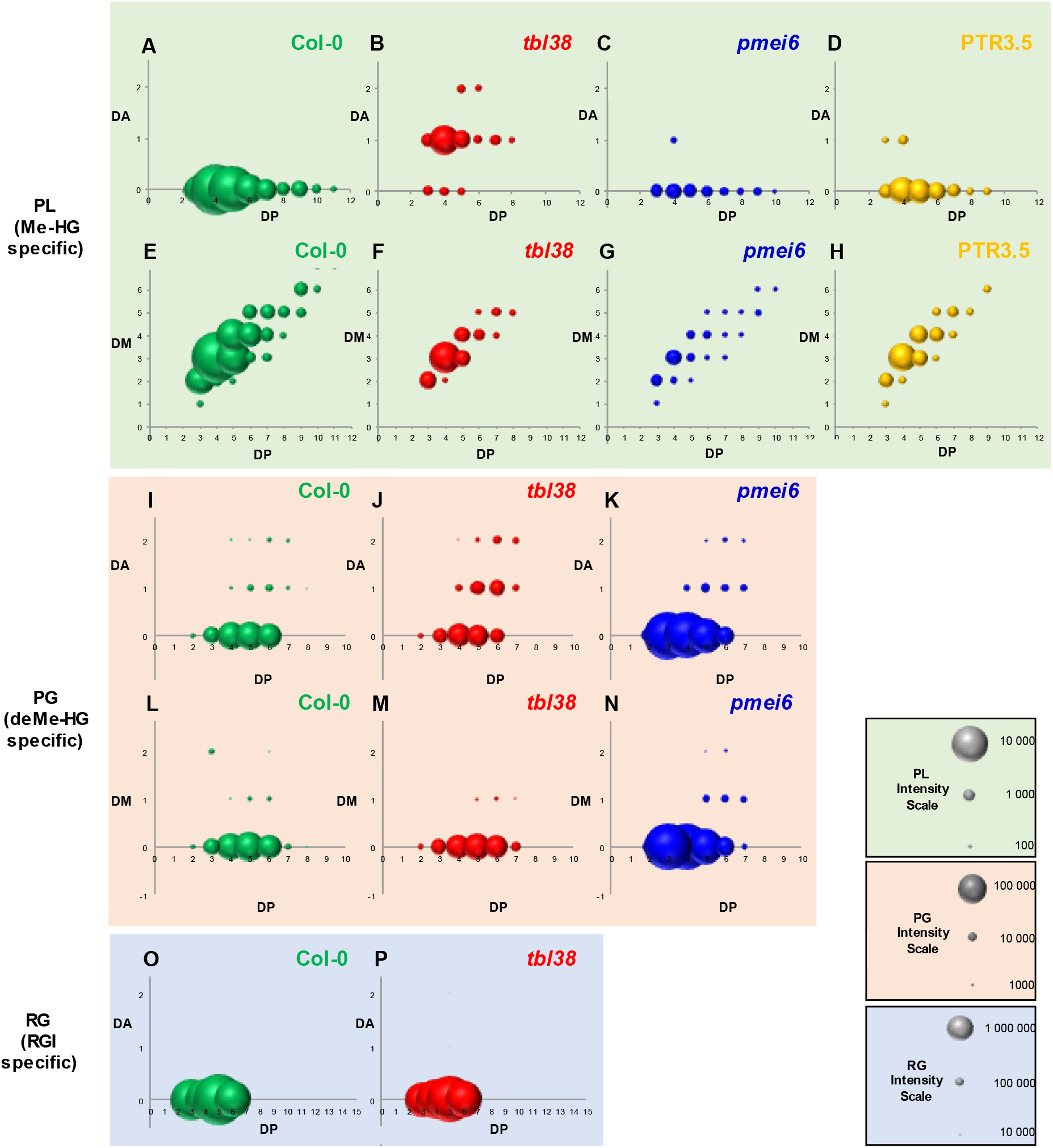
Enzymatic profiling reveals the HG acetylation qualitative phenotype of *tbl38* and the quantitative HG methylation phenotype of *pmei6*. MSC surface abrasion powders of Col-0, *tbl38*, *pmei6* and *tbl38* transformed with *pPRX36::TBL38-TagRFP* (line PTR3.5) were digested with pectin lyase (PL) specific for highly methylesterified HG **(A-H; Supplemental Table 3)**, polygalacturonase (PG) specific for low-methylesterified HG **(I-N; Supplemental Table 4)** or rhamnogalacturonan hydrolase (RG) specific for RGI backbone **(O-P; Supplemental Table 5)**. The identified HG species **(A-N)** or RGI species **(O-P)** were sorted according to their degree of acetylation (DA), degree of methylation (DM) and degree of polymerization (DP). The mean intensities were plotted either as DA vs DP or DM vs DP, regardless their DM and DA, respectively. See **Supplemental Table 3B**, **4A** and **5B** for details on the data treatment. The individual scales are provided on the Figure. Note the specific HG acetylation phenotype of *tbl38* with PL **(B)**, the qualitative complementation of PTR3.5 with PL **(D)**, the absence of RGI acetylation phenotype for *tbl38* (P) and the quantitative HG methylation phenotype of *pmei6* (G, N). These results argue for an HG acetylesterase role of TBL38 and are in consistent with the PMEI activity of PMEI6 (Saez-Aguayo et al., 2013).

The same analysis and data processing were performed with PG which has an affinity for lowly methylated HG (**Figure 6I-N; Supplemental Table 4A-D**). The DA vs DP profiles of Col-0, *tbl38* and *pmei6* were qualitatively similar though a slight increase of species of DA > 0 occurred for *tbl38* (**Figure 6I-K; Supplemental Table 4A-D**. The DM vs DP plots showed a quantitative phenotype of *pmei6* as compared to Col-0 and *tbl38* in agreement with the PMEI activity of PMEI6 (Saez-Aguayo et al., 2013) (**Figure 6L-N**). We also noticed an about ten-fold higher intensity of OGA species of low DM identified with PG (**Figure 6I-N; Supplemental Table 4A**) *vs* those of high DM species identified with PL (**Figure 6A-H; Supplemental Table 3B**). This could reflect the nature of the major HGs in MSCs (low DM and low DA) and/or be related to a possible lower activity of PL towards HGs with relatively high DA (Zeuner et al., 2020). To strengthen the HG high acetylation phenotype of *tbl38*, we additionally compared Col-0 and *tbl38* rhamnogalacturonate acetylation profiles to see whether TBL38 could also target the abundant RGI present in seed mucilage (**Figure 6O-P; Supplemental Table 5A**). No increased acetylation phenotype occurred in *tbl38* for RGI since the very strong intensity of the identified RGI species that were non-acetylated had similar profiles in both Col-0 and *tbl38.* This result demonstrates the specificity of the acetylation phenotype of *tbl38* towards HGs rather than RGI.

Finally, we produced recombinant TBL38 in *Pichia pastoris* (**Figure 7A**) and tested its acetyl esterase activity *in vitro* towards three acetylated substrates. Albeit low, we detected acetyl esterase activity using acetylated pectins from sugar beet and to a lesser extent with the generic acetylated substrate triacetin, while no significant activity could be detected using acetylated xylans (**Figure 7B**).

**Figure 7:**
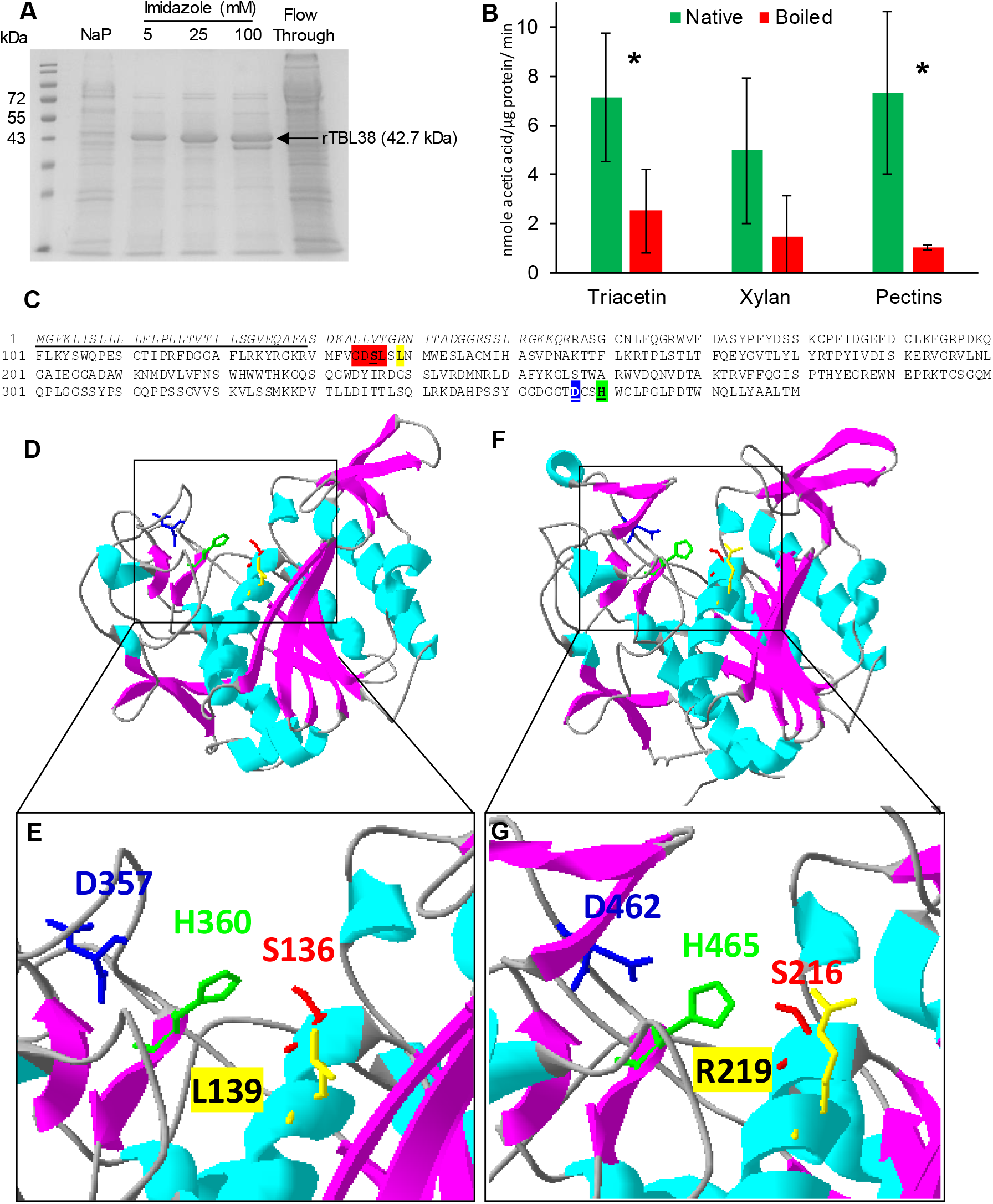
Pectin acetyl esterase activity of recombinant TBL38 and molecular modeling of TBL38 displaying its putative catalytic triad and substrate binding pocket. **(A)** SDS-PAGE showing the production of recombinant TBL38 (rTBL38) in *Pichia pastoris.* (B) *In vitro* enzymatic activity on acetylated substrates. The activity of the purified rTBL38 (100 mM imidazole fraction) was assayed using triacetin, acetylated xylans and sugar beet pectins as substrates and the ACET enzymatic acetic acid assay kit. Boiled samples were used as negative control, mean values ± SD, *, *p*-value < 0.05. **(C)** Amino acid sequence of TBL38 (At1g29050). The predicted signal peptide (https://services.healthtech.dtu.dk/services/SignalP-6.0/) is underlined. The amino acids in italics correspond to the 15% of the protein not covered by the molecular modeling performed in (D). GDSL and DxxS tetrads that are highly conserved among the 46 TBR/TBLs including TBL38 (Bischoff et al., 2010) are highlighted in red and blue, respectively. The amino acids participating to the putative catalytic triad S136-H360-D357 appear in bold underlined, in red, green and blue, respectively. L139 in TBL38 underlined in yellow replaces R219 in TBL29/ESK1/XOAT1 that was demonstrated to be necessary for acetyltransferase activity (Lunin et al., 2020). **(D)** The TBL38 structural model proposed in this study is a Phyre2 prediction (Kelley et al., 2015) with the highest ranking score (Confidence: 100.0%; Coverage: 85%). The template was the crystallographic structure (X-Ray diffraction, 1.85 Å) of *A. thaliana* TBL29 (At3g55990; PDB 6CCI; Phyred2 c6cciA) (Lunin et al., 2020). **(E)** Zoom on the putative catalytic triad S136-H360-D357 originally identified in Ser proteases (Dodson and Wlodawer, 1998), and necessary for the acyl-enzyme formation and acetyltransferase activity of TBL29 (Lunin et al., 2020). **(F and G)** structure and zoom on the catalytic triad of TBL29 for comparison (Lunin et al., 2020). Note that R219 necessary for acetyltransferase activity in TBL29 (Lunin et al., 2020) is replaced by L139 in TBL38.

## Discussion

### TBL38 is atypically positioned in the PRX36/PMEI6 cell wall microdomain of MSCs with a transient pattern

Originally, we searched for genes involved in the PMEI6-JIM7/LM20 HG epitopes-PRX36 molecular module (Francoz et al., 2019b) with a particular interest in HG remodeling enzymes. Using spatio-temporal co-expression with *PRX36*, we selected *TBL38* for its putative role in HG acetylation. Previously described TBLs are mostly anchored to the Golgi membrane (**Supplemental Table 1**) with the exception of TBR (Sinclair et al., 2017). Although the authors did not refute the possibility of some TBR-GFP signal in the Golgi, they concluded that TBR was a CW protein. The Golgi experimental localization matches with the N-terminal transmembrane domain prediction for 10 TBLs (TBL03, 27, 28, 29, 30, 31, 32, 33, 34, 35) while the Golgi-localized TBL37 has a predicted cleavable signal peptide (akin to a secreted CW protein), and conversely, TBR that has been localized to the CW has a predicted N-terminal transmembrane domain (**Supplemental Table 1**). Six TBLs including TBL38 have been identified in CW proteomes but these proteins did not match perfectly to the targeting prediction: there is a good match with cleavable signal peptide prediction for TBL9, 38, 39 and 45, but a bad match with transmembrane domain prediction for TBL15 and TBL40 (**Supplemental Table 1**). Here, we clearly localized TBL38-TagRFP in the CW of tobacco leaves or *A. thaliana* MSCs (**Figure 3**) in agreement with TBL38 presence in a CW proteome and its predicted cleavable signal peptide (**Supplemental Table 1**). In MSCs, we showed that this targeting is restricted to the CW microdomain harboring the PMEI6-JIM7/LM20 HG epitopes-PRX36 molecular module with an intriguing transient localization pattern (6-8 DAP). The localization to the CW microdomain occurs as soon as the fusion protein is produced just after the onset of *PRX36* promoter activity. We chose this promoter considering the strong co-expression of *PRX36* and *TBL38* and because it worked well with stable localization in various CW microdomains in previous studies (Francoz et al., 2019b; Kunieda et al., 2013; McGee et al., 2019; Sola et al., 2019). The physiological relevance of the transient localization pattern of TBL38-TagRFP is attested by the fact that the PTR lines displayed functional complementation in our different assays discussed hereafter. Then, the early delocalization of the fusion protein could be related to the loss of the putative anchoring motif (acetylated HG) as soon as TBL38 performed its acetylesterase activity, leading to degradation of the protein.

### TBL38 is a HG acetylesterase

So far, all described TBLs have been associated with a Golgi-localized acetyltransferase activity, either on hemicelluloses or pectins (**Figure 1; Supplemental Table 1**). Therefore, the question arose on whether the CW-localization of TBL38 could lead to similar activity. Yet, both acetic acid quantification and mass spectrometry analysis, as well as *in vitro* enzymatic assays on rTBL38 indicate that TBL38 is an acetylesterase targeting acetylated HGs as opposed to other TBLs (**Figures 5-7**). Sequence analysis of the TBL family highlighted the conservation of two characteristic domains: the GDS(L) and DXXH motifs commonly found in esterases (Akoh et al., 2004; Molgaard and Larsen, 2004). Crystallography analysis of TBL29/ESK1/XOAT1 pointed to the spatial reunion of these two domains forming the conserved Ser-His-Asp catalytic site, Ser coming from GDS(L) and Asp and His coming from DXXH (Lunin et al., 2020). TBL29 displayed an xylan acetyl transferase activity involving the formation of an acyl-enzyme intermediate on Ser129 (Lunin et al., 2020). Therefore, the acetyltransferase activity of TBL29 necessitated a generic acetylesterase activity on both the acetyl donor and the acyl-enzyme intermediate. Based on the crystallography structure of TBL29, we built an *in silico* model of TBL38 to determine the position of the above mentioned amino acids. The predicted catalytic site was correctly positioned in a putative polysaccharide binding groove (**Figure 7D, E**). Interestingly, in TBL29, the nearby R219 (**Figure 7F-G**) was shown to stabilize the catalysis (Lunin et al., 2020) as R219A site mutation led to loss of transfer of acetyl groups for the acetyl transferase activity without loss of the acetylesterase activity necessary for the acyl-enzyme formation (Lunin et al., 2020). TBL multiple alignment (Lunin et al., 2020) shows that 9 TBLs, including TBL38, do not have this particular Arg residue: TBL12 (E135), TBL30 (Y156), TBL37 (L143), TBL38 (L139) (**Figure 7C-E**), TBL39 (T125), TBL40 (L123), TBL41 (L112), TBL42 (N115), TBL43 (L122). All the other TBLs have an Arg except TBL44/PMR5 which possess another polar positive residue at this position (K145) and an acetyltransferase activity (Chiniquy et al., 2019). Among these 9 TBLs with no polar positive residue at this position, 4 correspond to the 6 TBLs identified in CW proteomes (**Supplemental Table 1**). It is therefore tempting to postulate that the absence of this residue could fit with the absence of a (Golgi-localized) acetyltransferase activity and the presence of a (CW-localized) acetylesterase activity as for TBL38. However, this cannot be the sole determinant since TBL37 and TBL30/XOAT3 both display a Golgi-localized acetyltransferase activity (Sun et al., 2020; Zhong et al., 2017). The characterization of TBL12, TBL39-43 could help in deciphering the importance of this residue at this specific position. Since we could not identify in TBL38 a particular motif or conformation which could explain such opposite esterase vs transferase activity, we propose that the opposite activity of TBL38 is primarily related to its CW localization. Indeed, TBL-dependent polysaccharide acetylation is done in two consecutive steps: hydrolysis of the acetyl group from the donor as evidence by the formation of the Acyl-enzyme intermediate, followed by transfer of the *O*-acetyl moiety to its polysaccharide target. To our knowledge, neither acetyl-CoA nor another putative acetyl donor is present in the CW, which could prevent the CW-localized TBL38 to complete the successive steps of the transferase reaction, the enzyme could then rather function as a direct acetylesterase on acetylated HGs. It should be noted that an esterase activity targeted on CW polymers was not observed for TBR despite its CW localization. *tbr* was associated with lower acetylation level of HGs (the opposite phenotype of *tbl38*), possibly through TBR-mediated protection of HG from pectin acetylesterases (Sinclair et al., 2017). Although is it unclear how such a mechanism could play out, it is difficult to explain the differences between the measured acetylation levels. Characterization of additional TBLs with regards to subcellular localization and acetylesterase activity is required for a better understanding of TBL roles.

### HG fine tuning of esterification is affected in *tbl38*

Although the control of HG acetylation and methylation is achieved by different proteins, proof of the direct and indirect interplay between both chemical groups was given. Indeed, HG acetylation could inhibit PME activity by steric hindrance (Celus et al., 2018; Kohn and Furda, 1968). Additionally, methyl ester groups are preferentially added on non-acetylated galacturonic acid moieties, which may explain the overall low level of HG acetylation observed in our results with Col-0 and in most plants (Pérez et al., 2000; Ralet et al., 2005), (**Figures 6 and 7**). Although we did not quantify HG methylesterification, the total number of identified species was particularly low in *tbl38*, when using a PL that has high affinity for highly methylesterified HGs (**Figure 6**). Even if we cannot exclude the possibility that the acetyl groups in *tbl38* negatively impact the PL activity, this could be associated with reduced HG methylesterification in *tbl38* considering the lack of LM20 labeling in *tbl38*. It should be noted, however, that the combined use of PG/PL might not give the full representation of HG species, since randomly methylated HG of DM around 50% are mediocre substrates for both PL and PG (Ralet et al., 2012). Additionally, JIM7 antibody still labels *tbl38* seeds suggesting that the lack of LM20 binding might rather be due to steric hindrance caused by HG acetylation, rather than changes in the pattern of methylesterification. Therefore, LM20 could display specificity for partially methylesterified and non-acetylated HGs, while JIM7 would be specific for partially methylesterified HGs regardless of the acetylation status. Interestingly, nanofibrils of HGs could be observed with LM20 but not with JIM7 in epidermis anticlinal CW from pavement cells (Haas et al., 2020; Haas et al., 2021). Considering the proposed different specificities of the two antibodies, one interpretation would be that the reduced steric hindrance and/or altered HG gelling properties (Ralet et al., 2003) related to the absence of acetyl groups would be necessary for proper assembly of HGs in nanofibrils. Evaluation of PME activity in *tbl38* developing MSCs could give clues as to TBL38 indirect influence on HG methylesterification. Regardless of the DM, our results confirmed increased amounts of HG acetylation in *tbl38* seed surface extracts. Contrary to methylesterification, acetylation does not influence the charge density and therefore the cation binding capacity of HGs (Dronnet et al., 1996; Ralet et al., 2003). However, experimental evidences have consistently associated highly acetylated HGs with reduced affinity to cations binding such as Ca^2+^ (Celus et al., 2018; Ralet et al., 2003). Conceptually, this could be due to direct influence of acetylation on conformation, access or cations binding properties, but also indirect effect on the PME activity as stated above (Oosterveld et al., 2000; Renard and Jarvis, 1999).

### Increased HG acetylation does not influence PRX36 anchoring

Our results have identified TBL38 as a CW localized HG acetylesterase with increased HG acetylation level in *tbl38* and no obvious effect on PRX36 positioning, despite the colocalization of TBL38 and PRX36 in the same CW microdomain. Previous work has demonstrated that PRX36 binding to HG was dependent on both methyl and Ca^2+^ as the PRX36-TagRFP localization was severely impacted in *pmei6* or using EDTA, respectively (Francoz et al., 2019b). To better understand the absence of PRX36 delocalization in *tbl38*, we investigated the implication of acetylated HGs on PRX36 docking using an *in silico* model based on PRX53 crystallographic data (Oosterveld et al., 2000). The previous *in silico* docking study tested the five OGAs of DP6 used to define JIM7 specificity (Clausen et al., 2003; Francoz et al., 2019b). It turned out that the three OGAs (so-called Clausen 3, 4 and to a lesser extend 5) that were the best recognized by JIM7 could be used for *in silico* docking on a PRX36 valley that was validated by site-directed mutagenesis (Francoz et al., 2019b). Here, we first randomly screened *in silico* PRX36 anchoring on 124 OGAs representing all the combinations of methylesterification from DP2 to DP6 **(Supplemental Table 6**). The *in silico* models consisted in nine poses of decreasing affinity (increasing energy level). To rationalize the distribution of these poses on the protein surface, we systematically measured the root mean square deviation (RMSD) showing the distance between the best pose and the eight other poses. We integrated the sum of energy levels and sum of RMSD for each OGA as a proxy of affinity and distances (**Supplemental Table 6**; see Methods). The 124 OGAs could then be sorted in a rationale - though of course imperfect manner-for their ability to bind to the PRX36 valley. This strongly reinforced the previous docking experiments made with *a priori* with Clausen1-5 OGAs (Francoz et al., 2019b) since Clausen 4 and Clausen 3 ranked 1 and 2 **(Supplemental Figure 6**; **Supplemental Table 6**). Finally, we tested the impact on *in silico* docking of the addition of 2-*O* or 3-*O* acetyl on the non-methylated galacturonic acids of Clausen 3 and 4 (**Supplemental Figure 6**). The addition of acetyl on hit number 2 (Clausen 3) strongly impaired PRX36 docking capacity, which does not fit with the observed results (no obvious modification of PRX36 anchoring and no mucilage release phenotype in *tbl38*). However, the addition of acetyl on hit number 1 (Clausen 4) did not have a strong impact on the docking capacity (**Supplemental Figure 6**). In turn, this is consistent with the immunolocalization of αPRX36 and confocal imaging of PRX36-TagRFP in *tbl38* background (**Supplemental Figure 4**) which did not show any obvious mislocalization of PRX36, suggesting that the Clausen 4 model might have a higher affinity towards PRX36 *in muro*.

In light of our results, we conclude that TBL38 acts as an HG acetylesterase in the PMEI6-JIM7/LM20 HG epitopes-PRX36 CW microdomain with no effect on PRX36 anchoring observed in *tbl38* contrarily to *pmei6*. The uncommon acetylesterase activity could be related to the localization to the CW were acetyl-CoA is absent. We could hypothesize that TBL38 could have an acetyltransferase activity if artificially targeted in a different environment such as the Golgi considering the presence of the canonical amino acids required for acetyltransferase activity. It remains to be understood why evolution has selected acetyltransferase-capable protein to deacetylate HGs instead of traditionally described PAEs, though in *A. thaliana*, no PAE have yet been linked to HG deacetylation. We were not able to associate the increased HG acetylation observed in *tbl38* to any clear developmental phenotype. However, considering the localization of this process to a remote CW microdomain, additional tools will be required to search for subtle developmental phenotypes, similarly to the development of the new original abrasive method allowing the characterization of the biochemical phenotype. Beyond this case study, this work paves the way for further understanding of the role of fine tuning of complex CW polysaccharides.

## Methods

### Plant material and growth conditions

*A. thaliana* mutants were ordered from the NASC for *tbl38*: SAIL_34_C03 (https://arabidopsis.info/; **Supplemental Figure 3**), or were previously available for *prx36* (SAIL_194_G03 (Kunieda et al., 2013)) and *pmei6* (SM_3.19557 (Saez-Aguayo et al., 2013)). Homozygous lines for *tbl38* were identified by PCR (**Supplemental Figure 3; Supplemental Table 7**). The knock out status of *tbl38* was determined by RT-PCR (**Supplemental Figure 3; Supplemental Table 7**). *A. thaliana* culture was performed as previously described (Francoz et al., 2019b).

*Nicotiana benthamiana* were cultivated in a growth chamber under a 16 h day / 8 h night cycle at 23°C (Neon (86.90 µmol/m/s-1) / 22°C upon 70 % humidity. They were transplanted after 14 days and fertilized each week.

### Transcriptomic data mining

*PRX36*, *PMEI6* and *TBL38* seed-specific expression profiles were obtained using the seed data source of eFP browser (http://bar.utoronto.ca/efp/cgi-bin/efpWeb.cgi?dataSource=Seed). Tissue-specific seed development transcriptomic data (Belmonte et al., 2013) was used to build the *PRX36* co-expression network (Francoz et al., 2019b). The *TBL* family (Bischoff et al., 2010) was filtered out.

### Bioinformatic analysis of the TBL family

Previous *TBL38* phylogeny (Bischoff et al., 2010) was used. The occurrence of TBLs in cell wall proteomes was assessed using http://www.polebio.lrsv.ups-tlse.fr/WallProtDB/. The topological prediction of the N-terminal hydrophobic sequence as being a transmembrane domain or a signal peptide was determined using http://aramemnon.uni-koeln.de/.

### Ruthenium Red Mucilage Release Test, Image Analysis and Statistical Analysis

High-throughput adherent mucilage release semi-quantitative phenotyping used the ruthenium red (**Supplemental Table 8**) staining method previously described allowing to calculate adherent mucilage area and circularity (Francoz et al., 2019b). The images were analyzed using ImageJ 1.8 (https://imagej.nih.gov/ij/) without edition of native images using an updated ImageJ script that facilitates cleaning of the data and that allows their automatic storage (see **Supplemental Methods** for more details).

### TagRFP reporting constructs, plant transformation and selection

The primers used for vector construction are listed in **Supplemental Table 7**. Full length cDNA clones and custom-ordered DNA plasmids used as template DNA for further cloning are listed in **Supplemental Table 9**. Level 0 GoldenGate generated constructs are listed in **Supplemental Table 10**. Level 1 GoldenGate plasmids assembled in pL1V-R2 vector and finalized Level 2 GoldenGate generated constructs are detailed in **Supplemental Table 11**. Level 1 plasmid was pICH47811 “pL1V-R2”pL1V-R2 vector (Weber et al., 2011). Level 2 plasmid was EC15027 “pL2V-HYG” with pICH471744 “pL1M-ELE-2” as a linker-containing plasmid (gift from Dr Pierre-Marc Delaux, LRSV, Auzeville-Tolosane, France). Every construct was checked by restriction analysis and sequencing prior to its transfer into *Agrobacterium tumefaciens* GV3101::pMP90 strain (Koncz and Schell, 1986). The transformed bacteria were grown in the presence of 15 µg.mL^−1^ gentamycin, 50 µg.mL^−1^ rifampicin and 25 µg.mL^−1^ hygromycin. *A. thaliana* plants were transformed by floral-dipping (Clough and Bent, 1998) or spraying flower buds with an Agromix containing 0.05 % (v/v) Silwet L-77 (De Sangosse 2000235). *proPRX36::PRX36-TagRFP* construct was transformed in *prx36* and *tbl38* plants. The *proPRX36::TBL38-TagRFP* construct was transformed in *tbl38* plants. These last complemented lines were referred as PTR2.4, PTR3.5 and PTR5.2 in the manuscript. Seeds were selected on Murashige and Skoog (MS) medium containing 25 µg.mL^−1^ hygromycin. Three independent homozygous transformed plant lines were studied for each construct.

The *pro35S::TBL38-TagRFP* construct was transiently co-transformed with either G-yb CD3-966 sialyltransferase-YFP Golgi marker or pm-yb-CD3-1006 aquaporin-YFP plasma membrane marker (Nelson et al., 2007) in 30-day old *N. benthamiana* leaves. The final inoculum consisted in a mix of various *A. tumefaciens* lines: 10 % (v/v) subcellular YFP-marker line, 80% (v/v) fusion protein-Tag-RFP line of interest, and 10 % (v/v) P19 silencing inhibitor-containing vector line. Leaf infiltration was done on the abaxial face through the stomata with a 1 mL syringe and pieces of leaves were mounted and analyzed under confocal microscopy 48 h-post agro-infiltration.

### Arabidopsis silique fixation, paraffin tissue array embedding and microtomy

The protocol was as previously described (Francoz et al., 2019b), with minor improvement using biopsy foam enabling handling more samples (see **Supplemental Methods** for more details).

### *In situ* RNA hybridization

*In situ* RNA hybridization experiments were performed as described (Francoz et al., 2019a; Francoz et al., 2016). More details are provided in **Supplemental Methods**.

### Immunofluorescence on tissue-array sections

We used our previously described protocol (Francoz et al., 2019b) with anti-PRX36 primary antibodies and goat-anti rabbit-A488 secondary antibodies (**Supplemental Table 8**) hybridized on serial sections of Col-0 and *tbl38* paraffin-embedded seed developmental kinetics. Serial sections of Col-0, *tbl38*, PTR2.4 and PTR3.5 developmental kinetics were similarly labeled with JIM7 or LM20 primary antibodies followed by goat anti-rat A488 secondary antibodies (**Supplemental Table 8**). More details are provided in **Supplemental Methods**.

### Confocal spinning disk microscopy of seed development kinetics of *A. thaliana* stable lines

Developing siliques taken from plants stably expressing the different TagRFP constructs in various genetic backgrounds were dissected and the replums containing the seeds were mounted under a coverslip in distilled water. Images were taken with a PLAN APO 20x/0.75 dry objective using the confocal spinning disk microscope from Perkin Elmer driven by the Volocity 6.3.0 software and equipped with a Yokogama CSU-X1 scan head, two EmCCD Hamamatsu C9100-13 cameras (Hamamatsu, Hamamatsu City, Japan) and a 580 nm beam splitter to separate dual staining on the two cameras as previously shown (Francoz et al., 2019b). Details of image processing are provided in the **Supplemental Methods**.

### Confocal microscopy of leaves of *N. benthamiana* transient transformants

Transiently transformed *N. benthamiana* leaves were observed 48 h post agroinfiltration using an upright confocal laser scanning microscope (LEICA SP8) with a 40 x apochromatic water immersion lens. TagRFP and YFP fluorescences were imaged with the following settings: excitation: 561 nm / emission: 582–622 nm; excitation 514 nm / emission: 527-530 nm, respectively. Z stacks (maximum intensity) were performed using ImageJ.

### SDS-PAGE and anti-TagRFP western-blots from *tbl38* complemented lines (PTR)

The TagRFP fluorescence patterns of PTR lines were monitored using spinning disk confocal microscopy along developmental kinetics of seeds from staged floral stem. The seeds that displayed the early (6-8 DAP), medium (8-12 DAP) and late (> 12 DAP) fluorescence patterns were carefully extracted from the dissected siliques (3-4 siliques per fluorescence pattern) and pooled in separated 2 mL tubes with a metallic grind ball before freezing in liquid nitrogen. Total proteins were extracted and analyzed by western blot as previously described (Francoz et al., 2019b), using anti-TagRFP primary and anti-rabbit-AP secondary antibodies (**Supplemental Table 8**) with minor modifications as detailed in **Supplemental Methods**.

### Dry seed MSC surface wall abrasive fraction

Abrasion columns were a home-made design allowing for homogenous dry seed MSC surface wall abrasive enrichment using plastic columns and collector tubes from the GeneJET Plasmid Miniprep Kit (Thermo Scientific K0502). The preparation of the columns, the abrasion procedure and the characterization of the efficiency are described in detail in **Supplemental Methods**.

### Immuno dotblot and semi-quantitative analysis from dry seed MSC abrasive fractions

Powder resulting of MSC surface abrasion was chemically extracted in the collector tube with extraction buffer (80 μL of 5 mM sodium acetate pH 4.6 containing 1 μL.mL^−1^ of plant protease inhibitor cocktail (Sigma P9599) per mg of powder). The collector tube with cap and a metallic grind ball was shaken at 250 rpm during 1 h at 4°C to ensure proper extraction. After a 5 min centrifugation at 14,400 rpm, the supernatant was recovered, diluted (1 μL in 50 μL milliQ-H_2_O) and deposited into sample wells onto a nitrocellulose membrane disposed into a 96-well Bio-Dot microfiltration apparatus (BioRad) previously soaked with in TBS (0.02 M Tris-HCl pH 7.5, 0.15 M NaCl) for 10 min. A 1 min 30 s of vacuum insured homogeneous transfer among the wells. The membrane was briefly air-dried to ensure proper adsorption. The detailed protocol of incubation with JIM7 and LM20 primary antibodies followed by goat anti-rabbit IgG-AP secondary antibodies as well as the semi-quantitative analysis of the results are provided in **Supplemental methods**.

### Acetyl group deesterification and acetic acid semi-quantitative analysis from dry seed MSC abrasive fractions

The acetylation of cell wall polymers in MSC surface wall abrasive samples was analyzed directly in the collection tube following adaptation of a previously described method (Stranne et al., 2018) using the K-ACET enzymatic Acetic Acid Assay Kit (Megazyme). The detailed protocol is provided in **Supplemental Methods**.

### Structural oligosaccharide composition of dry seed MSC abrasive fractions by IP-RP-UHPC-MS analysis

MSC surface wall powders were resuspended in 1 mL distilled water for 1 h. An aliquot (450 µL) was mixed with 50 µL of 500 mM sodium acetate buffer pH 5.0 and polysaccharides were hydrolyzed for 24 h at 40°C under 500 rpm shaking with either a pectin lyase (PL) specific for methylesterified HGs (Ralet et al., 2012), a polygalacturonase (PG) (Safran et al., 2023) specific for non-methylesterified HGs or a rhamnogalacturonase (RG, Swiss-Prot Q00018, provided by Novozymes, Copenhagen, Denmark) specific for the RGI backbone (Ralet et al., 2010). Resulting OGA digests were filtered on 0.45 µm and analyzed by IP-RP-UHPC-MS. Acquisitions were performed on a Select Series Cyclic IMS (Waters, Wilmslow, UK) coupled with a UHPLC system (Acquity H-Class Plus, Waters, Manchester, UK). The chromatographic separations were performed on an hypersil GOLD^TM^ (100 mm × 1 mm, packed with 1.9 μm porosity particles; Thermo-Fisher Scientific, Bremen, Germany). A ternary gradient was used (A: Milli-Q water, B: 100% methanol, and C: 20 mM heptylammonium formate, pH 6), from 2 % to 25 % of solvent B for 10 min, then up to 73 % at 23.5 min and maintained at 73 % for 4 min. Percentage of solvent C was kept constant at 25%. The flow rate was 0.175 mL min^−1^, and the column was heated to 45°C. Spectra were recorded in positive electrospray ionization (ESI) ionization mode in the m/z range 150–2000, with the TOF operating in the V-mode. The source parameters were as follows: capillary voltage 2.8 kV; cone voltage 120 V; source temperature 100°C; desolvation temperature 350°C; desolvation gas 350 L/h; and nebulization gas 6 bar. Data were recorded with the Quartz software (Waters embedded software, release 5) and processed using Mass Lynx 4.2 (Waters) and MzMine software (Pluskal et al., 2010). MzMine was used to produce a database of annotated structures (m/z; retention time; intensity). The identified species (charge states and anomers were merged) were annotated according to their degree of polymerization (DP), degree of methylesterification (DM) and degree of acetylation (DA). The data was analyzed using Microsoft Excel.

### Production of recombinant TBL38 in *Pichia pastoris* and enzymatic activity assays

The coding sequence of TBL38 (Q8VY22) was codon-optimized for *Pichia pastoris* and synthesized without signal peptide in frame with His-tag in pPCIZ-αB by ProteoGenix (Schiltigheim, France). rTBL38 was produced in *P. pastoris* following the previously described protocol (Lemaire et al., 2020). Activity assays of rTBL38 was performed with the acetic acid assay kit (K-ACETRM, Megazyme) using the activity buffer at 40°C and with three different acetylated substrates: 100 mM Triacetine, 10 mg.ml^−1^ xylan (24 % acetylation), and 10 mg.mL^−1^ sugar beet pectins (31 % acetylation). The detailed protocol is provided in **Supplemental Methods**.

### *In silico* model and docking simulations

Homology models for PRX36 and TBL38 (UniProt accession number Q9SD46 and Q8VY22, respectively) were built with the Phyre2 server (Kelley et al., 2015). For PRX36, the template was the crystallographic structure (X-Ray diffraction, 1.45 Å) of *A. thaliana* PRX53 (At5g06720; Protein Data Bank no.1PA2) (Ostergaard et al., 2000). For TBL38, the template was the crystallographic structure (X-Ray diffraction, 1.85 Å) of *A. thaliana* TBL29/ESK1/XOAT1 (Lunin et al., 2020). For comparison with the TBL38 model, the TBL29 structure was drawn as well using the 6cci pdb file (Lunin et al., 2020) visualized and analyzed with Swiss-PdbViewer (http://www.expasy.org/spdbv/) (Guex and Peitsch, 1997).

For PRX36 docking experiments, the α-D-(1-4) polygalacturonic acid structural model (Braccini et al., 1999) was retrieved from the Glyco3D portal (Perez et al., 2015) http://glyco3d.cermav.cnrs.fr/mol.php?type=polysaccharide&molecule=2504). It was modified to initially build the five hexagalacturonates models used to establish JIM7 specificity (Clausen et al., 2003). This selection was extended to the 124 oligogalacturonates (OGAs) from DP2 to DP6 covering all the theoretical combination of methylation (64 OGAs of DP6 + 32 OGAs of DP5 + 16 OGAs of DP4 + 8 OGAs of DP3 +4 OGAs of DP2). AutoDock Tools (Morris et al., 2009) and AutoDock Vina (Trott and Olson, 2010) were used for simulating the binding of the 124 OGAs to PRX36 within a search box encompassing the whole target protein enabling to recover the nine best poses for each OGA. Details on data processing are provided in **Supplemental Methods**.

## Acknowledgments

We thank the Paul Sabatier Toulouse 3 University and the Centre National de la Recherche Scientifique (CNRS) for granting this work. This work was specifically supported by the French National Research Agency project ‘‘MicroWall’’ (ANR-18-CE20-0007). It was also supported by the French Laboratory of Excellence project ‘‘TULIP’’ (ANR-10-LABX-41; ANR-11-IDEX-0002-02). We are grateful to Dr. H. North (IJPB, Versailles, France) and I.-H. Nishimura (Kyoto University, Japan) for providing seeds and to Dr. P.-M. Delaux (LRSV, Auzeville-Tolosane, France) for the gift of GoldenGate plasmids and helpful discussions about Golden Gate cloning.

## Author contributions

Conceptualization, V.B.; Methodology, all authors; Investigation, B.G.D, D.R., M.R., P.R., C.C., S.O., J. M.-B., P.T., A.G. and V.B.; Writing – Original Draft Writing, B.G.D. and V.B.; Review & Editing, B.G.D., D.R., P.R., E.J., C.D., J.P., M.-C.R. and V.B. All authors read and approved the manuscript; Funding Acquisition, J.P., M.-C.R. and V.B.; Resources, J.P., M.-C.R. and V.B; Supervision, J.P., M.-C.R. and V.B.

