## Supplementary material for "TBL38 is an atypical homogalacturonan acetylesterase with a peculiar cell wall microdomain localization in Arabidopsis seed mucilage secretory cells": Sup Material

**Supplemental data:**

**Supplemental Figure 1: Spinning disk confocal microscopy observations of the developmental kinetics of three independent complemented lines of *tbl38* transformed with *pPRX36::TBL38-TagRFP*.**

**Supplemental Figure 2: αTagRFP Western blot analysis of *tbl38* transformed with *pPRX36::TBL38-TagRFP* (complemented line PTR3.5).**

**Supplemental Figure 3: *tbl38* knock out mutant characterization.**

**Supplemental Figure 4: *tbl38* seeds show no defect in mucilage release and accordingly no obvious consequence on PRX36 anchoring.**

**Supplemental Figure 5: Characterization of the abrasion process of dry seed surface to produce a mucilage secretory cell (MSC) surface-enriched fraction**

**Supplemental Figure 6: The acetylation of the best oligogalacturonate hit for PRX36 docking does not impair the docking prediction.**

**Supplemental Table 1: Phylogeny, subcellular-localization and activity of TBL enzymes in *A. thaliana*.**

**Supplemental Table 2: Selection of *TBL38* in *PRX36*/*PMEI6* co-expression network for further functional genomics and biochemical studies.**

(**Supplemental Tables 3-5: Excel files**)

**Supplemental Table 6: Random systematic screening of molecular docking of PRX36 on 124 oligogalacturonates (OGAs) of DP6-D2 covering all theoretical demethylesterification patterns highlights the specific interaction of PRX36 with the two hexagalacturonates constituting the epitope of JIM7.**

**Supplemental Table 7: PCR oligonucleotide primers.**

**Supplemental Table 8: Histochemical dyes, primary and secondary antibodies.**

**Supplemental Table 9: cDNA and custom-ordered plasmids used as template DNA for further cloning.**

**Supplemental Table 10: Level 0 Golden Gate generated constructs.**

**Supplemental Table 11: Level 1 in pL1V-R2 vector and finalized Level 2 Golden Gate generated constructs.**

**Supplemental Methods**

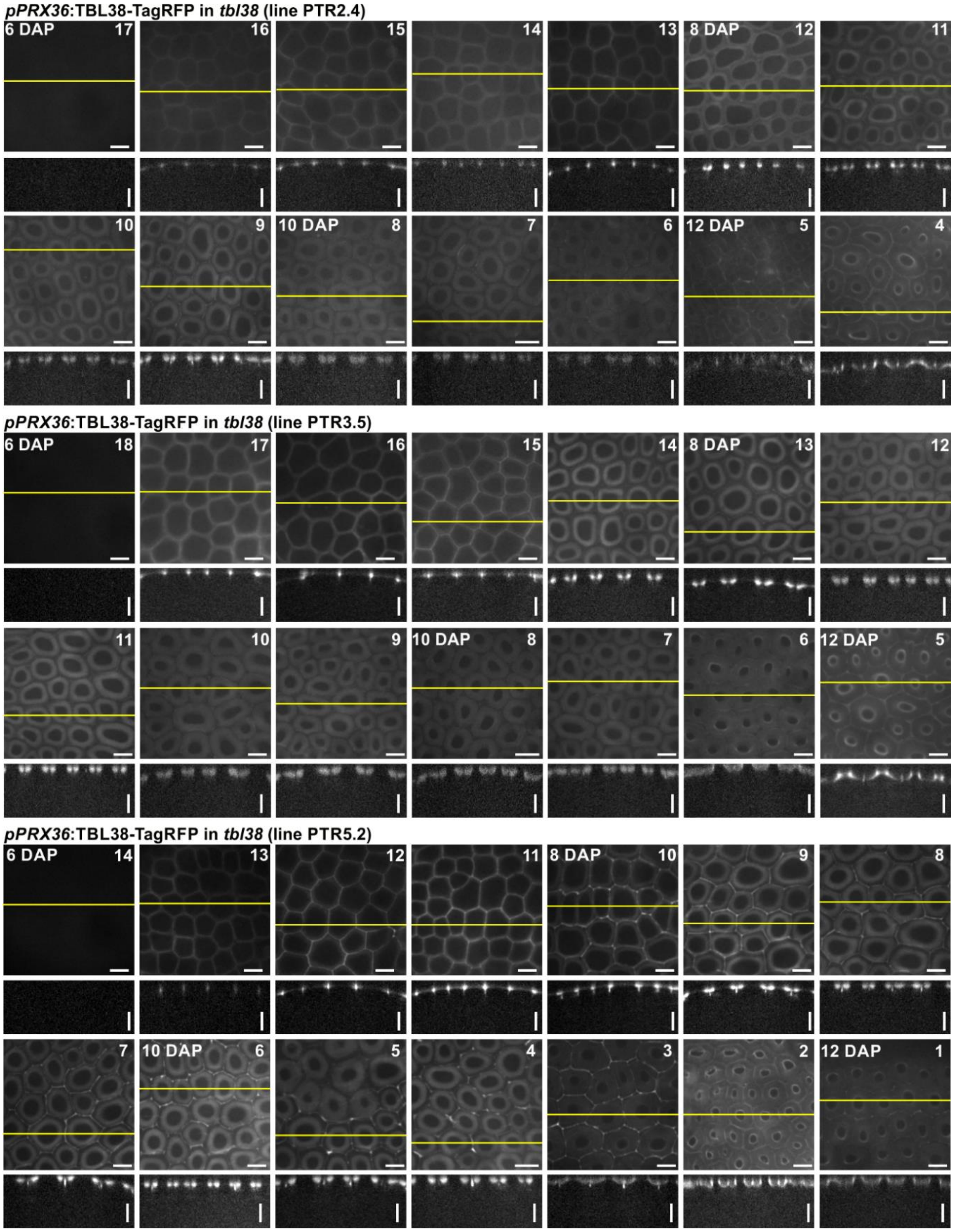


**Supplemental Figure 1: Spinning disk confocal microscopy observations of the developmental kinetics of three independent complemented lines of tbl38 transformed with *pPRX36::TBL38-TagRFP.*** For lines PTR2.4, 3.5 and 5.2, seeds from 14 consecutive silique ranks were observed. Each image is representative of > 10-20 seeds within each silique. Silique ranks on the floral stems are labelled in the top right corner of the images; the stage of selected siliques is labelled in day after pollination (DAP) on the top left corner of the images. The sum view of 70 confocal stacks are presented with a unique intensity setting (the absolute min and max intensity values were applied to all images to ensure fair comparison of the fluorescence intensities). The yellow lines correspond to the position of the orthogonal view shown below each image. Note that following the onset of *pPRX36* activity at 6 DAP, the localization of the fluorescence to the MSC CW microdomain at the top of the radial CW occurs during 3-4 ranks and is followed by a gradual delocalization of the fluorescence to the mucilage pocket, Horizontal and vertical bars: 25 µm.

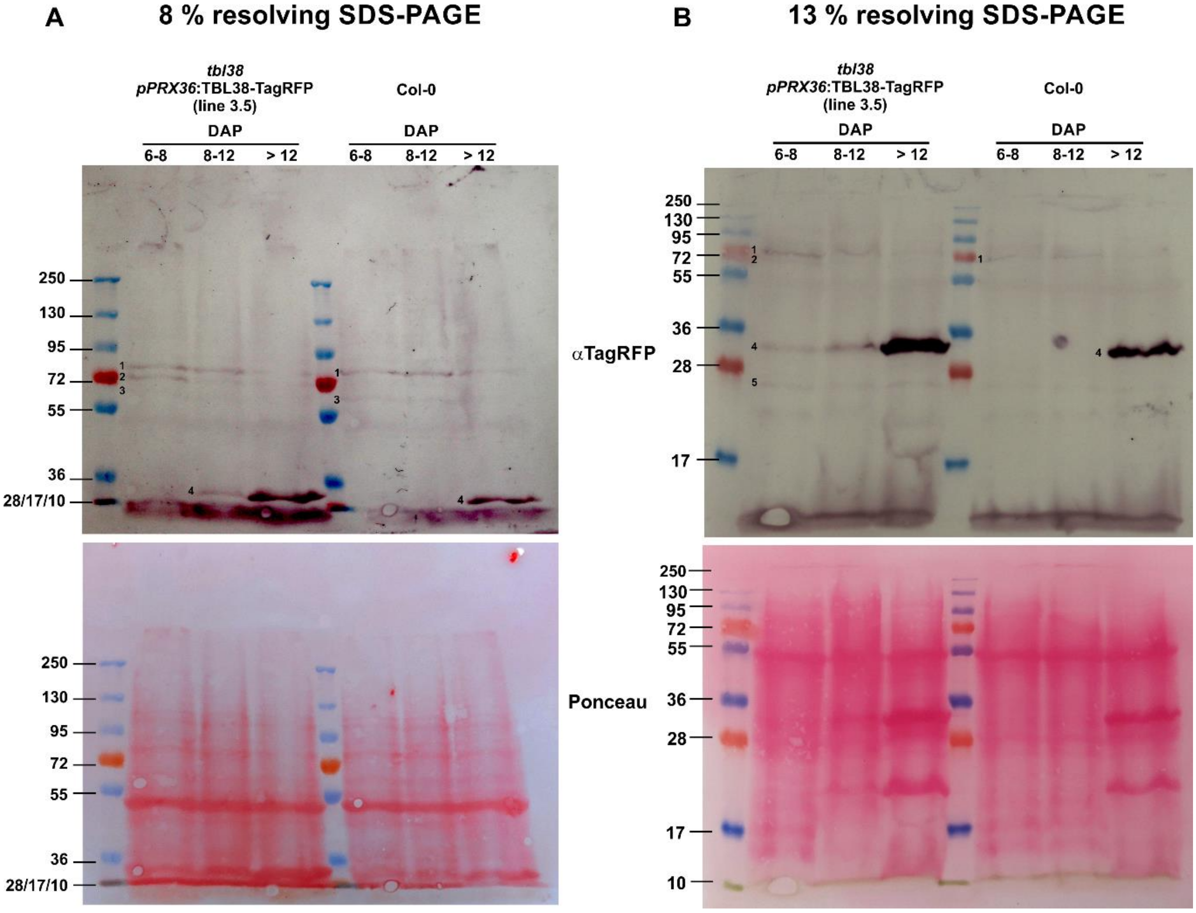


**Supplemental Figure 2: αTagRFP Western blot analysis of *tbl38* transformed with *pPRX36::TBL38-TagRFP* (complemented line PTR3.5).** Developmental kinetics of line PTR3.5 siliques were controlled for fluorescence patterns with Spinning disk confocal microscopy and seeds with the three fluorescence patterns were carefully sampled (seeds from 3-4 silique ranks per sample). Similar seeds from Col-0 used as a negative control were sampled according to the silique ranks. Protein extracts were run on 8 % **(A)** or 13 % **(B)** SDS-PAGE and αTagRFP Western blot were performed following Ponceau red staining of the membranes. Only the youngest samples (6-8 DAP) that displayed a fluorescence pattern in the CW microdomain at the top of radial MSC CW show the specific band # 2 corresponding to TBL38-TagRFP fusion proteins. This band is no longer seen in 8-12 DAP and > 12 DAP samples that displayed a delocalization of the fluorescence to the mucilage pocket (see **Figure 3** and **Supplemental Figure 2**). Bands # 1 and 3 are non-specific since these are also present in the Col-0 control samples. Band # 4 may correspond to a degradation product of TBL38-TagRFP as well as an additional non-specific band of similar molecular mass since it is also seen at late stages in Col-0 and could not be distinguished even on 13 % SDS PAGE.

A

B

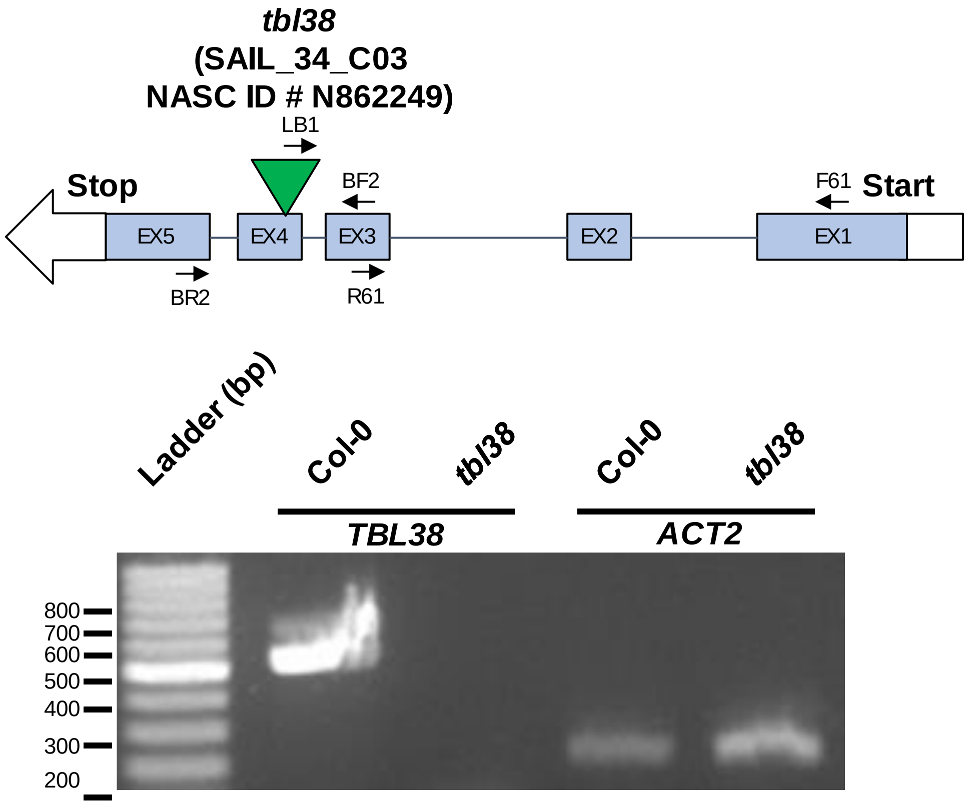


C

| Primer code | Primer sequence (5'-3') | use |
|---|---|---|
| BF2 | CTTGGCATTGGTGGACTCACAAAG | Genotyping |
| BR2 | CATCGAACTCAACACTTTCGACACTACTCC | Genotyping |
| LB1 (SAIL T-DNA) | GCCTTTTCAGAAATGGATAAATAGCCTTGCTTCC | Genotyping |
| F61 | AAAACCCAAAACCTTTGATATTTGTCTAA | RT-PCR |
| R61 | CACTAGAAGATTTACGAATGTCATTTACAGA | RT-PCR |

**Supplemental Figure 3: *tbl38* knock out mutant characterization. (A)** Genomic map of *TBL38* (*AT1G29050*) and T-DNA insertion position in *tbl38* mutant (SAIL_34_C03 ; NASC ID # N862249 in Col-3). The green triangle indicates that the insertion site of the T-DNA in the mutant line determined by sequencing is in exon 4 and not in intron 3 as reported in TAIR data (https://www.arabidopsis.org/. **(B)** RT-PCR amplification of *TBL38* and *ACTIN2* (*ACT2*) used as a loading control performed with 40 cycles to challenge the putative residual low expression levels in *tbl38* vs Col-0 wild type control. Similar result was found on cDNAs prepared from two independent sampling of 6 to 8 DAP-old siliques. **(C)** Primer sequences used for genotyping and RT-PCR (see position in panel **A**)

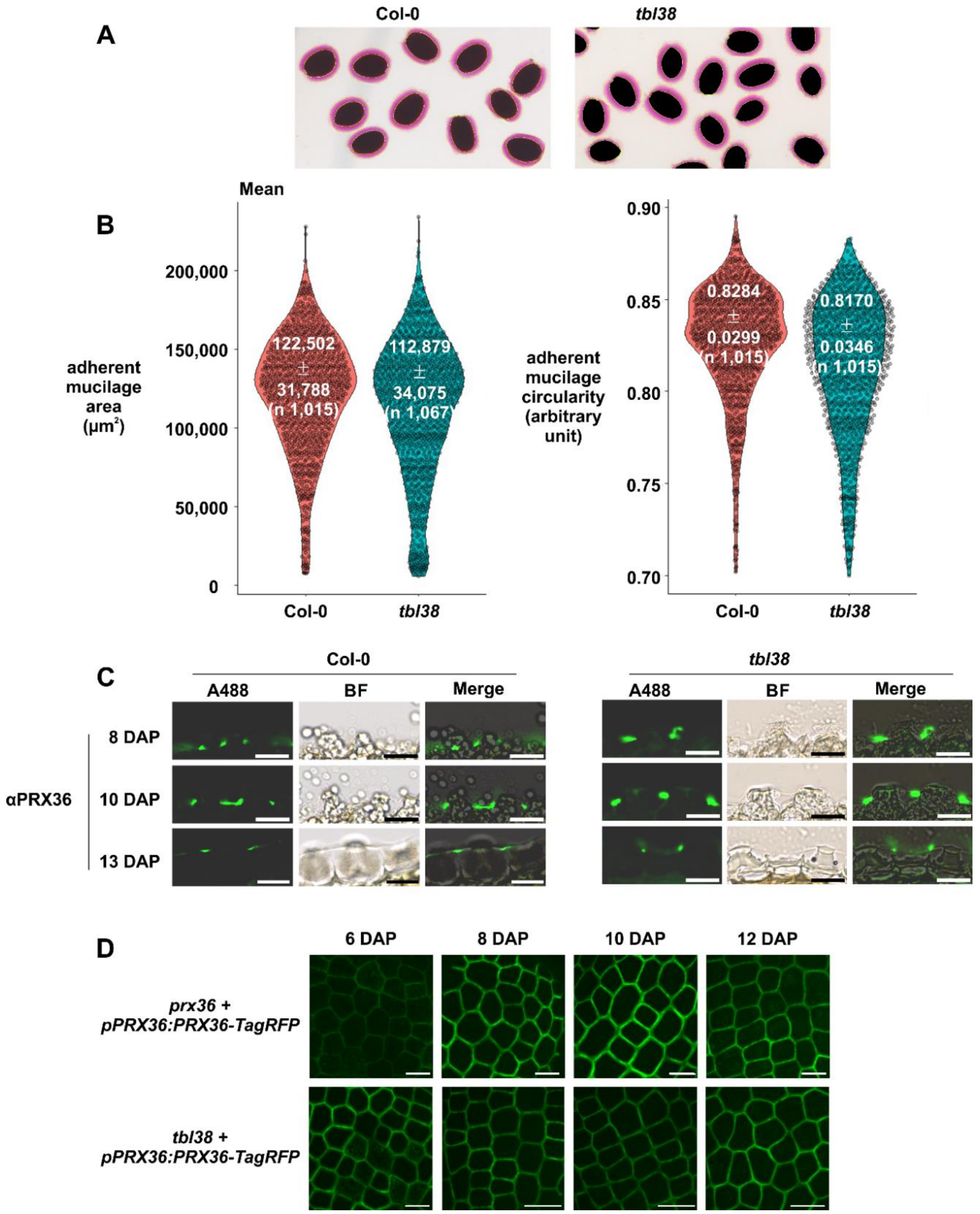


**Supplemental Figure 4: *tbl38* seeds show no defect in mucilage release and accordingly no obvious consequence on PRX36 anchoring. (A)** Example of screenshot from scan of Ruthenium red staining of released adherent mucilage shows similar patterns between Col-0 and *tbl38*. Note the yellow masks used in (B). *tbl38* does not show the phenotypes previously observed in *prx36* and *pmei6* (Francoz et al., 2019b; Kunieda et al., 2013; Saez-Aguayo et al., 2013). **(B)** Violin plot representation of measurement of adherent mucilage area and circularity using the yellow masks shown in (A) confirm the absence of any obvious phenotype. **(C)** αPRX36 labeling of paraffin sections positioned PRX36 at the top of radial primary cell wall from MSC in both Col-0 and *tbl38* contrary to the previously demonstrated loss of labeling in *prx36* and *pmei6* (Francoz et al., 2019b). Bars: 25 µm. **(D)** PRX36-TagRFP localization is not affected in *tbl38* developing seeds. Confocal spinning disk observations of stable *proPRX36::PRX36-TagRFP* in *A. thaliana* reveal no obvious mislocalization of the signal in *tbl38*. Similar protein accumulation at 6 DAP was observed in both genotypes. Images are the result of a maximum projection of similar stacks (Z>30) which were not edited. Bars: 50 µm.

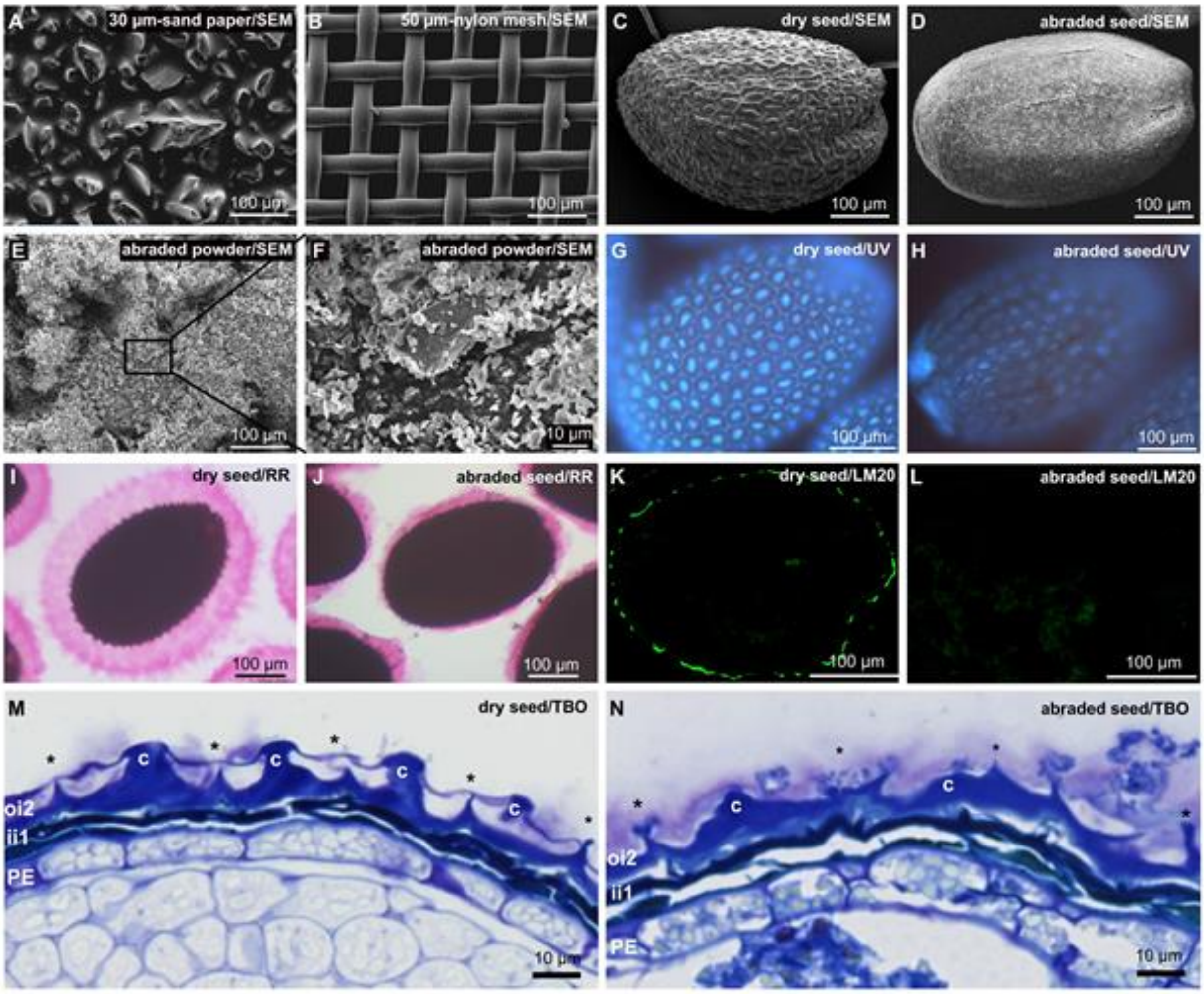


**Supplemental Figure 5: Characterization of the abrasion process of dry seed surface to produce a mucilage secretory cell (MSC) surface-enriched fraction.** Scanning electron microscopy (SEM) view of sand paper **(A)**, nylon mesh **(B)** included in the home-made abrasion column (see methods), dry seed before **(C)** and after **(D)** 5 x 1 min abrasion illustrating the homogeneous etching of the dry seed surface. SEM view of the recovered MSC surface-enriched powder **(E, F)** illustrating the small size and relatively homogeneous particles. Note that the same scale is used for **A** to **E** to better compare the sizes. UV imaging of dry seed before **(G)** and after **(H)** abrasion illustrating the loss of autofluorescence of MSC radial walls and the decrease of autofluorescence of columella. Ruthenium red (RR) staining of dry seed before **(I)** and after **(J)** abrasion showing the remaining thin layer of adherent mucilage. LM20 immunofluorescence of dry seed before **(K)** and after **(L)** abrasion showing that the LM20 cell wall microdomain is abraded. Toluidine blue O (TBO) staining of dry seed before **(M)** and after **(N)** abrasion further showing that the etching occurred only at the surface of the MSC. *, radial primary wall, c, columella, oi2, outer integument 2, ii1, inner integument 1, PE, peripheral endosperm.

(A)

| RK with ac | RK | Clausen et al 2013 | OGAs (code and structure) | DP | DM | DA | Σ EL/ Σ RMSD (RANKED) | Σ EL | Affinity EL = Energy level (kcal/mol) 9 poses of docking vs all surface of PRX36: 1 | 2 | 3 | 4 | 5 | 6 | 7 | 8 | 9 | Σ RMSD | Distance RMSD (Root-mean-square deviation) 9 poses of docking vs all surface of PRX36: 1 | 2 | 3 | 4 | 5 | 6 | 7 | 8 | 9 |
|---|---|---|---|---|---|---|---|---|---|---|---|---|---|---|---|---|---|---|---|---|---|---|---|---|---|---|---|
| 1 | | Clausen 4 *O*2-ac | 063b-ó8888ó | 6 | 50% | 50% | -1.2099 | -54.9 | -6.8 | -6.6 | -6.3 | -6.1 | -6.1 | -5.9 | -5.8 | -5.7 | -5.6 | 45.4 | 0.0 | 3.3 | 2.7 | 5.3 | 4.6 | 30.8 | 6.2 | 5.3 | 29.2 |
| 2 | 1 | Clausen 4 | 063-o8888o | 6 | 67% | 0% | -1.1306 | -51.3 | -6.7 | -6.2 | -5.7 | -5.6 | -5.6 | -5.4 | -5.4 | -5.4 | -5.3 | 45.4 | 0.0 | 2.1 | 2.6 | 2.9 | 2.2 | 4.3 | 2.0 | 27.4 | 1.9 |
| 3 | | Clausen 4 *O*3-ac | 063c-ò8888ò | 6 | 50% | 50% | -1.0667 | -48.4 | -5.7 | -5.4 | -5.4 | -5.4 | -5.3 | -5.3 | -5.3 | -5.3 | -5.3 | 45.4 | 0.0 | 2.2 | 16.4 | 16.5 | 4.5 | 4.5 | 19.2 | 3.2 | 16.4 |
| 4 | 2 | Clausen 3 | 047-oo888o | 6 | 50% | 0% | -1.0269 | -53.8 | -6.7 | -6.3 | -6.2 | -6.1 | -5.9 | -5.8 | -5.7 | -5.6 | -5.5 | 52.4 | 0.0 | 6.1 | 4.9 | 4.2 | 5.0 | 7.0 | 1.8 | 4.6 | 18.8 |
| 98 | | Clausen 3 *O*2-ac | 047b-óó888ó | 6 | 50% | 50% | -0.3459 | -62.8 | -7.1 | -7.1 | -7.1 | -7.0 | -7.0 | -7.0 | -6.9 | -6.9 | -6.7 | 181.6 | 20.4 | 0.0 | 16.9 | 17.1 | 9.8 | 30.8 | 35.1 | 17.1 | 34.4 |
| 122 | | Clausen 3 *O*3-ac | 047c-òò888ò | 6 | 50% | 50% | -0.2731 | -61.0 | -7.0 | -6.9 | -6.9 | -6.8 | -6.8 | -6.7 | -6.7 | -6.6 | -6.6 | 223.4 | 0.0 | 30.8 | 18.3 | 26.5 | 31.4 | 19.2 | 31.4 | 34.6 | 31.0 |

(B) Clausen 4 (C) Clausen 4 *O*2-ac (D) Clausen 4 *O*3-ac (E) Clausen 3 (F) Clausen 3 *O*2-ac (G) Clausen 3 *O*3-ac

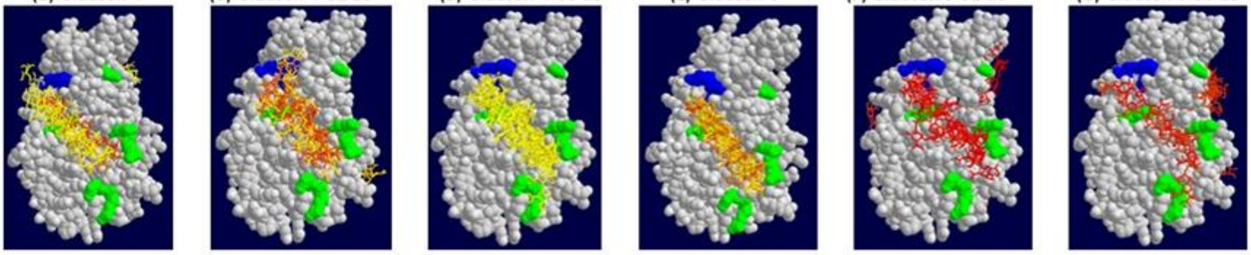

**Supplemental Figure 6: The acetylation of the best oligogalacturonate (OGA) hit for PRX36 docking does not impair the docking prediction. (A)** The two best hits obtained following random systematic screening of molecular docking of PRX36 on 124 OGAs (DP6-DP2) covering all theoretical demethylesterification patterns correspond to two OGAs of DP6 constituting the epitope of JIM7 (**Supplemental Table 6**). These two OGAs named Clausen 3 and Clausen 4 (Francoz et al., 2019b) were further acetylated *in silico* in their non-methylated galacturonic acids either in *O*-2 or *O*-3 positions. The molecular docking performed on the whole PRX36 surface was compared between the non-acetylated and acetylated OGAs using autodock vina showing the 9 best poses including their individual energy level (EL) and Root-mean-square deviation (RMSD) indicating the affinity of the interaction (the lower, the better) and the distance in Angstrom of the 9 poses for a given OGA (the lower, the shorter), respectively. (i) Data mining was performed as detailed in **Supplemental Table 6** and methods and the OGAs were sorted according to SEL/SRMSD used to integrate the affinity and the distance (the lowest, the highest affinity and shorter distance). Red-to-yellow and blue-to-yellow heatmaps were drawn for EL and RMSD, respectively. The EL or RMSD values appearing in bold underlined frames correspond to poses fitting to the PRX36 valley previously demonstrated (Francoz et al., 2019b) to accommodate the hexagalacturonates Clausen 3, 4 and 5 (047-oo888o, 063-o8888o and 027-8o8o8o) that are recognized by the JIM7 monoclonal antibody (Clausen et al., 2003). **Note that the docking specificity of the best hit (Clausen 4) is not significantly modified between the non-acetylated and the two acetylated versions whereas the docking of the second best hit (Clausen 3) is strongly impaired following acetylation. (B-G)** visualization of the docking models for the six combinations of hexagalacturonates. Key: RK, rank; DP, degree of polymerization; DM, degree of methylation; DA, degree of acetylation; OGA, oligogalacturonate; o, demethylesterified galacturonic acid; ó, *O*2-acetylated demethylesterified galacturonic acid;,ò, *O*3-acetylated demethylesterified galacturonic acid; 8, methylated galacturonic acid.The red and blue amino acids in **(B-G)** correspond to polar positive and hydrophobic amino acids from the OGA docking valley whose experimental mutation impaired PRX36 localization (Francoz et al., 2019b). The OGAs in **(B-G)** were color coded following the energy level heat map shown in **(A)**

**Supplemental Table 1: Phylogeny, subcellular-localization and activity of TBL enzymes in *A. thaliana*.**

| Phylogeny | | | Subcellular localization | | | | | | *In vitro* Assay | | | Reverse genetic | | | | |
|---|---|---|---|---|---|---|---|---|---|---|---|---|---|---|---|---|
| Name | Gene ID | Clade[a] | Aramemnon topology[b] | WP DB[c] | Subcell loc[d] | Fluo Tag[e] | Species[f] | Activity and function[g] | Recom-binant protein[h] | Substrate[i] | Product[j] | Biochemical phenotype[k] | Develop-mental phenotype[l] | Comple-mentation[m] | Misce-llaneous | Reference |
| TBR | At5g06700 | A | TM? | | CW | GFP | *A.th.* | Pectin-AT or protection of *O*-acetylated pectin from de-acetylation by pectin acetyl-esterases in the CW | | | | Reduced secondary wall cellulose (crystalline) content; Increased PME activity; Reduced pectin acetylation; Increased pectin methyl-esterification | Absence of secondary wall in trichomes; Reduced etiolated hypocotyl elongation | Full, with 35S promoter and native promoter | | (Bischoff et al., 2010; Potikha and Delmer, 1995; Sinclair et al., 2017) |
| TBL01 | At3g12060 | A | TM? | | | | | | | | | | | | | |
| TBL02 | At1g60790 | A | TM? | | | | | | | | | | | | | |
| TBL04 | At5g49340 | A | SP? | | | | | | | | | | | | | |
| TBL05 | At5g20590 | A | TM? | | | | | | | | | | | | | |
| TBL06 | At3g62390 | A | TM? | | | | | | | | | | | | | |
| TBL07 | At1g48880 | A | TM? | | | | | | | | | | | | | |
| TBL08 | At3g11570 | A | TM? | | | | | | | | | | | | | |
| TBL09 | At5g06230 | A | TM? | etiolated hypo-cotyls | | | | | | | | | | | | (Feiz et al., 2006) |

| | | | | | | | | | | | | | | | | |
|---|---|---|---|---|---|---|---|---|---|---|---|---|---|---|---|---|
| TBL10 | At3g06080 | A | TM? | | | | *A.th.* | Putative RGI-AT | | | | Reduced acetylation of RGI in rosette leaves | enhanced drought tolerance | Two independent KO mutants | | (Stranne et al., 2018) |
| TBL11 | At5g19160 | A | TM? | | | | | | | | | | | | | |
| TBL12 | At5g64470 | B | TM? | | | | | | | | | | | | | |
| TBL13 | At2g14530 | B | SP? | | | | | | | | | | | | | |
| TBL14 | At5g64020 | B | TM? | | | | | | | | | | | | | |
| TBL15 | At2g37720 | B | TM? | leaves | | | | | | | | | | | | (Durufle et al., 2019) |
| TBL16 | At5g20680 | B | TM? | | | | | | | | | | | | | |
| TBL17/YLS7 | At5g51640 | | TM? | | | | | | | | | | | | yellow-leaf-specific gene | (Yoshida et al., 2001) |
| | | | | | | | | | | | | | | | Causal candidate gene underlying the QTL4 for root length in drougtht GWAS | (El-Soda et al., 2015) |
| TBL18 | At4g25360 | B | TM? | | | | | | | | | | | | | |
| TBL19/XyBAT1 | At5g15900 | B | TM? | | | | | xyloglucan (XG) backbone AT (XyBAT) | human HEK293F cells (secreted form) | Cello-hexaose | acetyl groups onto 6-*O* of Glc residues in cellohexaose (XG backbone) | Heterologous expression of a rice XG backbone AT (XyBAT) in Arabidopsis led to a severe reduction in cell expansion and plant growth and a drastic alteration in xyloglucan xylosylation pattern with the non classical acetylation of XG backbone: XGG, XGGG, XXGG, XXGG, XXGGG and XXGGG (G, non acetylated Glc; G acetylated Glc; X, xylosylated Glc) | | | TBL19,20, 21 are orthologs from rice and tomato XyBAT acetylating XG backbone. TBL19 and 21 activity | (Zhong et al., 2020) |

| | | | | | | | | | | | | | | | | |
|---|---|---|---|---|---|---|---|---|---|---|---|---|---|---|---|---|
| TBL20 | At3g02440 | B | TM? | | | | | no | human HEK293F cells (secreted form) | Cello-hexaose | no activity | | | | is intriguing since acety-lation of XG backbone is reported only in grasses and Solanaceae. It might exist in specialized cells. *In planta*, the XG backbone might be first acetylated and then xylosylated | (Zhong et al., 2020) |
| TBL21/XyBAT2 | At5g15890 | B | TM? | | | | | xyloglucan (XG) backbone AT (XyBAT) | human HEK293F cells (secreted form) | Cello-hexaose | acetyl groups onto 6-*O* of Glc residues in cellohexaose (XG backbone) | | | | | (Zhong et al., 2020) |
| TBL22/AXY4L/XGOAT2 | At3g28150 | B | TM? | | | | | xyloglucan (XG) side chain AT and weak esterase activity | human HEK293F cells (secreted form) | Fucosylated Gal of XG side chain | acetyl groups onto 6-*O* (3-*O*, 4-*O* and 4,6-di-*O*) of fucosylated Gal residues on XG side chains. | Drastic reduction of acetylated XG in seeds | | | | (Gille et al., 2011; Zhong et al., 2018b) |
| TBL23/MOAT1 | At4g11090 | B | TM? | | | | | gluco-mannan AT | human HEK293F cells (secreted form) | Manno-hexaose | acetyl groups onto 2-*O* and 3-*O* of Man residues on Mannohexaose | Simultaneous RNAi inhibition of the expression of *MOAT1, 2, 3* and *4* leads to a drastic reduction in glucomannan acetylation in | None detected | | | (Zhong et al., 2018a) |
| TBL24/MOAT2 | At4g23790 | B | TM? | | | | | gluco-mannan AT | human HEK293F cells (secreted form) | Manno-hexaose | | | | | | (Zhong et al., 2018a) |

| | | | | | | | | | | | | | | | | |
|---|---|---|---|---|---|---|---|---|---|---|---|---|---|---|---|---|
| TBL25/MOAT3 | At1g01430 | B | TM? | | | | | gluco-mannan AT | human HEK293F cells (secreted form) | Manno-hexaose | side chains. | Arabidopsis floral stem | | | | (Zhong et al., 2018a) |
| TBL26/MOAT4 | At4g01080 | B | TM? | | | | | gluco-mannan AT | human HEK293F cells (secreted form) | Manno-hexaose | | | | | | (Zhong et al., 2018a) |
| TBL27/AXY4/XGOAT1 | At1g70230 | B | TM? | | Golgi | GFP | *N. bentha.* | xyloglucan (XG) side chain AT and weak esterase activity | human HEK293F cells (secreted form) | Fucosylated Gal of XG side chain | acetyl groups onto 6-*O* (3-*O*, 4-*O* and 4,6-di-*O*) of fucosylated Gal residues on XG side chains. | drastic reduction of acetylated XG in seedlings, roots, leaves | | | TBL27 acts down-stream of MYB103 to affect Al sensitivity through XG *O*-acety-lation | (Gille et al., 2011; Wu et al., 2022; Zhong et al., 2018b; Zhu et al., 2014) |
| TBL28/XOAT2 | At2g40150 | C | TM? | | Golgi | m-Cherry | *N. bentha.* | xylan AT | human HEK293F cells (secreted form) | Xylan (Xyl6) | **Xyl-2Ac**; Xyl-3Ac; Xyl-2,3Ac | | No obvious secondary wall phenotype | | | (Zhong et al., 2017) |

| | | | | | | | | | | | | | | | | |
|---|---|---|---|---|---|---|---|---|---|---|---|---|---|---|---|---|
| TBL29/XOAT1/ESK1 | At3g55990 | C | TM? | | Golgi | YFP | N/A | xylan AT necessary for correct xylan substitution enabling correct interaction with cellulose microfibrils | human HEK293F cells (secreted form) | Xylan (Xyl6) | **Xyl-2Ac; Xyl-3Ac; Xyl-2,3Ac** | The recombinant protein activity is confirmed in *esk1* stems | collapsed vessels | Various patterns of complementation with *ESK1*, *TBL28* or *TBL30* | Crystal structure allowing molecular modeling of other TBLs | (Grantham et al., 2017; Lefebvre et al., 2011; Lunin et al., 2020; Xiong et al., 2013; Yuan et al., 2013; Zhong et al., 2017) |
| TBL30/XOAT3 | At2g40160 | C | TM? | | Golgi | m-Cherry | *N. bentha.* | xylan AT | human HEK293F cells (secreted form) | Xylan (Xyl6) | Xyl-2Ac; **Xyl-3Ac** | | | | | (Zhong et al., 2017) |
| TBL03/XOAT4 | At5g01360 | C | TM? | | Golgi | YFP | *A.th.* proto. | xylan AT | human HEK293F cells (secreted form) | Xylan (Xyl6) | **Xyl-2Ac; Xyl-3Ac**; Xyl-2,3Ac | Decreased cellulose, Ara, Gal Rha, Fuc and uronic acid content; Increased PME activity | Reduced inflorescence stem elongation | | | (Bischoff et al., 2010; Yuan et al., 2016b; Zhong et al., 2017) |
| TBL31/XOAT5 | At1g73140 | C | TM? | | Golgi | YFP | *A.th.* proto. | xylan AT | human HEK293F cells (secreted form) | Xylan (Xyl6) | Xyl-2Ac; **Xyl-3Ac**; Xyl-2,3Ac | | | | | (Yuan et al., 2016b; Zhong et al., 2017) |

| | | | | | | | | | | | | | | | | |
|---|---|---|---|---|---|---|---|---|---|---|---|---|---|---|---|---|
| TBL32/XOAT6 | At3g11030 | C | TM? | | Golgi | YFP | *A.th.* proto. | xylan AT | human HEK293F cells (secreted form) | Xylan (GlcA)Xyl4 | **Xyl-3Ac-2GlcA** | | | | | (Yuan et al., 2016c; Zhong et al., 2017) |
| TBL33/XOAT7 | At2g40320 | C | TM? | | Golgi | YFP | *A.th.* proto. | xylan AT | human HEK293F cells (secreted form) | Xylan (GlcA)Xyl4 | **Xyl-3Ac-2GlcA** | | | | | (Yuan et al., 2016c; Zhong et al., 2017) |
| TBL34/XOAT8 | At2g38320 | C | TM? | | Golgi | YFP | *A.th.* proto. | xylan AT | human HEK293F cells (secreted form) | Xylan (Xyl6) | **Xyl-2Ac; Xyl-3Ac;** Xyl-2,3Ac | | | | | (Yuan et al., 2016a; Zhong et al., 2017) |
| TBL35/XOAT9 | At5g01620 | C | TM? | | Golgi | YFP | *A.th.* proto. | xylan AT | human HEK293F cells (secreted form) | Xylan (Xyl6) | **Xyl-2,3Ac** | | | | | (Yuan et al., 2016a; Zhong et al., 2017) |
| TBL36 | At3g54260 | D | SP? | | | | | | | | | | | | | |
| TBL37 | At2g34070 | D | SP? | | Golgi | GFP | *N. bentha.* | Hemicellulose-AT Stem and leaves | | | | Reduced poly-saccharide (hemicellulose) acetylation | Dwarf, collapsed vessels, reduced secondary CW thickening in fibers, reduced mesophyll cell size | Full, pTBL37::TBL37 | Directly activated by MYC2 Enhances herbivore resi-stance | (Sun et al., 2020) |

| | | | | | | | | | | | | | | | | |
|---|---|---|---|---|---|---|---|---|---|---|---|---|---|---|---|---|
| TBL38 | At1g29050 | D | SP? | leaves ; stems | CW | Tag RFP | *A. th.; N. bentha.* | Homo-galacturonan-AE Localized to a CW microdomain of seed mucilage secretory cells | *P. pastoris* | HG | | Increase in *O*-acetylation of homogalacturonan; Loss of LM20 but not JIM7 epitopes | | Almost full, pPRX36::TBL38-TagRFP | | (Durufle et al., 2019) This study |
| TBL39 | At2g42570 | D | SP? | roots | | | | | | | | | | | | (Nguyen-Kim et al., 2016) |
| TBL40 | At2g31110 | D | TM? | roots | | | | | | | | | | | | (Nguyen-Kim et al., 2016) |
| TBL41 | At3g14850 | D | SP? | | | | | | | | | | | | | |
| TBL42 | At1g78710 | D | SP? | | | | | | | | | | | | | |
| TBL43 | At2g30900 | D | SP? | | | | | | | | | | | | | |
| TBL44/PMR5 | At5g58600 | D | SP? | | Endo-membranes? | GFP | *A.th.* | Homogalacturonan-AT | *E. coli* | Oligo-Galacturonides | Acetylated oligo-galacturonides | Altered pectin, decreased acetate and decreased cellulose contents | Increased pathogen resistance | Full, pTBL44::TBL44-GFP | Alteration of pathogen resi-stance | (Chiniquy et al., 2019) |
| TBL45 | At2g30010 | D | SP? | inflorescence | | N/A | *A.th.* | | | | | | | | | (Xu et al., 2016) |

[a] Phylogeny according to Bischoff et al., (2010). The clade letters were added here for sake of clarity ; [b] Prediction of transmembrane anchoring (TM?) or signal peptide cleavage (SP?) according to http://aramemnon.uni-koeln.de/ ; [c] Occurrence in cell wall proteomes (http://www.polebio.lrsv.ups-tlse.fr/WallProtDB/) ; [d] Experimentally assessed subcellular localization (CW, cell wall) ; [e] Fluorescent tag used for subcellular localization ; [f] Species in which the localization was performed (*A. th., A. thaliana*; proto., protoplast; *N. bentha., N. benthamiana*); [g] Summary of enzymatic activity and function (AT, acetyl transferase; AE, acetyl esterase) ; [h] Heterologous system used for recombinant protein production; [i] Polysaccharide oligomer used as a substrate for *in vitro* activity ; [j] *In vitro* product ; [k] Biochemical phenotype observed in KO mutant, RNAi lines or overexpressors ; [l] Developmental phenotype observed in KO mutant, RNAi lines or overexpressors; [m] Construct used for complementation assay. "Full" indicates a full complementation. N/A, non available.

**Supplemental Table 2: Selection of *TRICHOME BIREFRINGENCE-LIKE38* (*TBL38)* in the *PRX36/PMEI6* co-expression network for further functional genomics and biochemical studies.** *PMEI6* is the first hit of the *PRX36* co-expression network using the tissue-specific seed developmental kinetics GSE12404 dataset (Belmonte et al., 2013). Mutants from both genes have similar mucilage phenotype (Koua et al., 2009; Saez-Aguayo et al., 2013) and a molecular link has been established to explain the similar phenotype: PMEI6 activity enables to produce a homogalacturonan specific methylesterification pattern necessary for PRX36 anchoring to the cell wall microdomain of mucilage secretory cells (Francoz et al., 2019b). The accurately positioned PRX36 further loosens the CW microdomain to enable correct mucilage release upon seed imbibition. In an effort to search new proteins involved in HG remodeling, *TBL38* labeled in green has been selected for functional genomic study. Relative expression values are colored from yellow to red. Expression below the 45-cutoff are represented in grey and genes that never reach the cutoff are masked (*TBL5; TBL26/MOAT4; TBL23/MOAT1; TBL18; TBL21/XyBAT2; TBL17/YLS7; TBL07; TBL19/XyBAT1;TBL03/XOAT4;TBL20; TBL44/PMR5; TBL27/AXY4/XGOAT1; TBL34/XOAT8*). No information was available for: *TBL02; TBL04; TBL08; TBL09; TBL11; TBL14; TBL24/MOAT2; TBL29/XOAT1/ESK1; TBL31/XOAT5; TBL33/XOAT7*. RK, rank; PCC, Pearson Correlation Coefficient; EP, embryo proper; SUS, suspensor; MCE, micropylar endosperm; PEN, peripheral endosperm; CZEN, chalazal endosperm; CZSC, chalazal seed coat; SC, seed coat; WS, whole seed.

| RK | PCC | ORF/AGI | pre-globular stage | | | | | | | globular stage | | | | | | | | heart stage | | | | | | | linear-cotyledon stage | | | | | | | bending cotyledon stage | | | | | | mature green stage | | | | | |
|---|---|---|---|---|---|---|---|---|---|---|---|---|---|---|---|---|---|---|---|---|---|---|---|---|---|---|---|---|---|---|---|---|---|---|---|---|---|---|---|---|---|---|---|
| | | | pg EP | pg MCE | pg PEN | pg CZEN | pg CZSC | pg SC | pg WS | g EP | g SUS | g MCEN | g PEN | g CZEN | g CZSC | g SC | g WS | h EP | h MCEN | h PEN | h CZEN | h CZSC | h SC | h WS | lc EP | lc MCEN | lc PEN | lc CZEN | lc CZSC | lc SC | lc WS | bc EP | bc PEN | bc CZEN | bc CZSC | bc SC | bc WS | mg EP | mg MCEN | mg PEN | mg CZEN | mg CZSC | mg SC |
| 1 | 1,0000 | *PRX36* | 15 | 27 | 4 | 11 | 8 | 8 | 2 | 4 | 2 | 13 | 2 | 12 | 8 | 2 | 4 | 3 | 3 | 3 | 15 | 14 | 3 | 9 | 4 | 5 | 9 | 12 | 405 | 1522 | 554 | 1 | 2 | 27 | 992 | 29 | 20 | 7 | 1 | 11 | 3 | 53 | 65 |
| 2 | 0,9474 | *PMEI6* | 2 | 15 | 7 | 8 | 2 | 1 | 6 | 2 | 1 | 2 | 1 | 2 | 2 | 6 | 6 | 2 | 8 | 8 | 4 | 10 | 6 | 13 | 10 | 54 | 54 | 1 | 52 | 423 | 236 | 6 | 34 | 14 | 232 | 65 | 52 | 29 | 11 | 19 | 25 | 37 | 93 |
| 13 | 0,8160 | *TBL38* | 9 | 11 | 25 | 34 | 2119 | 876 | 1094 | 10 | 5 | 61 | 26 | 2 | 2076 | 1003 | 1508 | 8 | 9 | 8 | 55 | 1064 | 1790 | 1012 | 16 | 47 | 7 | 17 | 1274 | 5537 | 2991 | 44 | 1 | 1 | 2731 | 894 | 312 | 1262 | 10 | 31 | 9 | 582 | 135 |
| 5 198 | 0,1004 | *TBL40* | 5 | 32 | 5 | 25 | 104 | 3 | 70 | 1 | 2 | 13 | 3 | 1 | 64 | 11 | 27 | 3 | 6 | 2 | 17 | 166 | 9 | 15 | 1 | 7 | 1 | 1 | 95 | 2 | 10 | 1 | 1 | 3 | 67 | 4 | 1 | 6 | 10 | 10 | 1 | 57 | 9 |
| 6 959 | 0,0498 | *TBL06* | 46 | 12 | 8 | 50 | 88 | 237 | 243 | 69 | 20 | 84 | 8 | 2 | 54 | 184 | 98 | 52 | 38 | 27 | 3 | 56 | 232 | 81 | 124 | 37 | 23 | 1 | 35 | 129 | 70 | 94 | 38 | 4 | 26 | 77 | 56 | 101 | 78 | 37 | 42 | 10 | 61 |
| 6 994 | 0,0490 | *TBL45* | 197 | 49 | 22 | 25 | 41 | 203 | 171 | 107 | 437 | 19 | 14 | 8 | 21 | 271 | 260 | 100 | 36 | 14 | 14 | 28 | 663 | 272 | 74 | 262 | 1 | 41 | 51 | 224 | 114 | 25 | 20 | 103 | 30 | 88 | 53 | 19 | 12 | 5 | 61 | 60 | 291 |
| 7 753 | 0,0306 | *TBL10* | 96 | 113 | 103 | 50 | 169 | 123 | 70 | 79 | 344 | 490 | 159 | 28 | 276 | 162 | 90 | 108 | 294 | 252 | 27 | 286 | 159 | 186 | 50 | 44 | 610 | 23 | 383 | 82 | 132 | 39 | 199 | 111 | 204 | 47 | 69 | 19 | 13 | 26 | 80 | 152 | 40 |
| 7 844 | 0,0285 | *TBL30/XOAT3* | 13 | 14 | 13 | 30 | 26 | 61 | 50 | 10 | 11 | 48 | 11 | 9 | 10 | 95 | 38 | 12 | 17 | 21 | 2 | 18 | 119 | 47 | 10 | 9 | 11 | 12 | 19 | 44 | 30 | 9 | 11 | 43 | 16 | 25 | 21 | 22 | 15 | 6 | 13 | 17 | 239 |
| 9 223 | -0,0026 | *TBL13* | 48 | 16 | 36 | 13 | 31 | 51 | 55 | 51 | 35 | 7 | 20 | 4 | 49 | 46 | 68 | 72 | 46 | 65 | 12 | 48 | 42 | 68 | 105 | 55 | 72 | 3 | 23 | 53 | 51 | 61 | 47 | 40 | 42 | 55 | 55 | 45 | 68 | 58 | 51 | 40 | 64 |
| 10 258 | -0,0224 | *TBL12* | 53 | 65 | 91 | 45 | 49 | 50 | 48 | 43 | 57 | 154 | 58 | 53 | 36 | 42 | 51 | 41 | 60 | 79 | 104 | 34 | 45 | 106 | 69 | 58 | 57 | 148 | 56 | 77 | 46 | 75 | 91 | 46 | 40 | 49 | 84 | 49 | 17 | 44 | 32 | 48 | 48 |
| 12 225 | -0,0541 | *TBL28/XOAT2* | 15 | 21 | 9 | 9 | 15 | 10 | 16 | 1 | 1 | 4 | 16 | 15 | 5 | 9 | 9 | 26 | 11 | 1 | 20 | 6 | 11 | 21 | 32 | 85 | 5 | 4 | 13 | 25 | 61 | 42 | 31 | 9 | 6 | 9 | 37 | 195 | 203 | 145 | 78 | 2 | 15 |
| 12 396 | -0,0565 | *TBL22/AXY4L/XGOAT2* | 10 | 2 | 9 | 28 | 77 | 58 | 42 | 6 | 120 | 51 | 17 | 13 | 25 | 39 | 24 | 88 | 911 | 160 | 17 | 35 | 39 | 162 | 22 | 693 | 697 | 93 | 33 | 151 | 142 | 74 | 430 | 269 | 93 | 24 | 168 | 480 | 90 | 140 | 590 | 53 | 61 |
| 13 137 | -0,0662 | *TBL35/XOAT9* | 156 | 8 | 40 | 9 | 67 | 108 | 86 | 99 | 142 | 27 | 22 | 3 | 25 | 48 | 70 | 85 | 23 | 13 | 27 | 60 | 127 | 47 | 84 | 22 | 13 | 188 | 26 | 48 | 56 | 52 | 26 | 232 | 51 | 22 | 38 | 18 | 45 | 75 | 78 | 31 | 44 |
| 13 339 | -0,0687 | *TBL43* | 8 | 25 | 28 | 2625 | 56 | 10 | 54 | 4 | 2 | 15 | 36 | 2277 | 68 | 21 | 70 | 4 | 4 | 7 | 270 | 5 | 1 | 3 | 2 | 2 | 3 | 2 | 5 | 17 | 7 | 2 | 2 | 17 | 3 | 17 | 10 | 2 | 7 | 3 | 6 | 3 | 5 |
| 13 378 | -0,0691 | *TBL15* | 599 | 10218 | 162 | 204 | 14 | 8 | 391 | 95 | 1403 | 5370 | 7 | 598 | 12 | 20 | 238 | 2 | 33 | 8 | 82 | 13 | 5 | 6 | 3 | 9 | 4 | 2 | 4 | 1 | 7 | 1 | 8 | 2 | 1 | 2 | 4 | 5 | 5 | 9 | 2 | 4 | 13 |
| 14 329 | -0,0800 | *TBL42* | 480 | 7881 | 1079 | 856 | 2 | 32 | 252 | 55 | 850 | 3336 | 679 | 643 | 5 | 13 | 107 | 0 | 22 | 33 | 217 | 9 | 7 | 18 | 2 | 11 | 2 | 7 | 3 | 5 | 2 | 4 | 1 | 12 | 6 | 10 | 5 | 2 | 6 | 4 | 2 | 2 | 14 |
| 14 985 | -0,0876 | *TBR* | 81 | 10 | 45 | 11 | 162 | 160 | 205 | 80 | 23 | 27 | 38 | 17 | 159 | 155 | 105 | 105 | 146 | 77 | 33 | 126 | 192 | 129 | 133 | 238 | 72 | 25 | 131 | 98 | 105 | 114 | 172 | 32 | 71 | 147 | 105 | 162 | 479 | 294 | 232 | 86 | 111 |
| 15 156 | -0,0897 | *TBL39* | 105 | 20 | 14 | 23 | 81 | 75 | 26 | 258 | 16 | 21 | 16 | 24 | 42 | 47 | 45 | 123 | 51 | 42 | 29 | 87 | 18 | 36 | 90 | 34 | 11 | 2 | 64 | 6 | 62 | 114 | 14 | 41 | 50 | 17 | 93 | 51 | 14 | 16 | 19 | 22 | 2 |
| 15 766 | -0,0978 | *TBL41* | 2225 | 19267 | 1673 | 10135 | 183 | 164 | 1631 | 72 | 2266 | 7572 | 69 | 8757 | 423 | 16 | 476 | 9 | 30 | 5 | 2234 | 83 | 22 | 53 | 5 | 16 | 16 | 19 | 75 | 36 | 17 | 9 | 10 | 126 | 119 | 5 | 11 | 13 | 9 | 2 | 22 | 85 | 14 |
| 16 074 | -0,1021 | *TBL36* | 4 | 4 | 1 | 5 | 13 | 2 | 12 | 56 | 5 | 3 | 2 | 4 | 15 | 14 | 2 | 105 | 4 | 2 | 3 | 1 | 1 | 30 | 114 | 50 | 6 | 2 | 1 | 4 | 46 | 73 | 6 | 1 | 2 | 2 | 53 | 164 | 379 | 3 | 7 | 1 | 7 |
| 16 513 | -0,1081 | *TBL01* | 65 | 11 | 18 | 34 | 32 | 29 | 30 | 68 | 91 | 7 | 4 | 23 | 37 | 19 | 11 | 109 | 48 | 25 | 19 | 31 | 19 | 33 | 185 | 48 | 24 | 4 | 35 | 18 | 57 | 162 | 11 | 13 | 23 | 9 | 64 | 140 | 9 | 17 | 22 | 16 | 21 |
| 17 654 | -0,1236 | *TBL37* | 13 | 19 | 6 | 14 | 10 | 160 | 71 | 76 | 6 | 9 | 10 | 10 | 16 | 110 | 98 | 31 | 13 | 9 | 20 | 28 | 98 | 42 | 25 | 3 | 3 | 16 | 14 | 11 | 40 | 41 | 6 | 8 | 9 | 15 | 24 | 102 | 1 | 8 | 5 | 13 | 12 |
| 17 817 | -0,1265 | *TBL25/MOAT3* | 9 | 49 | 19 | 23 | 42 | 453 | 227 | 23 | 3 | 25 | 7 | 19 | 23 | 246 | 193 | 12 | 15 | 24 | 8 | 29 | 177 | 45 | 69 | 72 | 143 | 5 | 12 | 43 | 53 | 92 | 181 | 22 | 24 | 43 | 122 | 87 | 59 | 76 | 56 | 2 | 11 |
| 18 136 | -0,1321 | *TBL32/XOAT6* | 656 | 790 | 533 | 178 | 658 | 1020 | 675 | 927 | 979 | 1326 | 1002 | 538 | 1043 | 1719 | 344 | 786 | 1432 | 1023 | 164 | 599 | 667 | 749 | 732 | 649 | 1571 | 879 | 825 | 731 | 452 | 566 | 1059 | 732 | 590 | 1182 | 381 | 937 | 763 | 657 | 1284 | 433 | 1008 |
| 21 035 | -0,1912 | *TBL16* | 42 | 221 | 16 | 24 | 1 | 36 | 25 | 43 | 40 | 104 | 28 | 68 | 15 | 42 | 16 | 43 | 22 | 43 | 2 | 5 | 10 | 10 | 40 | 66 | 38 | 11 | 15 | 10 | 31 | 62 | 46 | 62 | 9 | 21 | 34 | 58 | 37 | 65 | 64 | 34 | 29 |

**Supplemental Tables 3-5: Excel files**

**Supplemental Table 6: Random systematic screening of molecular docking of PRX36 on 124 oligogalacturonates (OGAs) of DP6-D2 covering all theoretical demethylesterification patterns highlights the specific interaction of PRX36 with the two hexagalacturonates constituting the epitope of JIM7.** For each OGA, the molecular docking performed on the whole PRX36 surface using autodock vina enabled retrieving the nine best poses including their individual energy level (EL) and Root-mean-square deviation (RMSD) indicating the affinity of the interaction (the lower, the better) and the distance in Angstrom of the 9 poses for a given OGA (the lower, the shorter), respectively. (i) Data mining was performed by calculating the sum of the EL (ΣEL) reflecting the global affinity of each OGA for PRX36 and the sum of RMSD (ΣRMSD) reflecting the dispersion vs concentration of the nine poses on the protein surface.(ii) Then, the OGAs were sorted according to ΣEL/ΣRMSD used to integrate both parameters (the lowest, the highest affinity and shorter distance). Red-to-yellow and blue-to-yellow heatmaps were drawn for EL and RMSD, respectively. The EL or RMSD values appearing in bold underlined frames correspond to poses fitting to the PRX36 valley previously demonstrated (Francoz et al., 2019b) to accommodate the hexagalacturonates clausen 3, 4 and 5 (047-oo888o, 063-o8888o and 027-8o8o8o) that are recognized by the JIM7 monoclonal antibody (Clausen et al., 2003). o, demethylesterified galacturonic acid, 8, methylated galacturonic acid. Note that the docking specificity is neither related to the degree of polymerization (DP) nor to the degree of methylation (DM), but rather relies on a peculiar demethylesterification bar code.

| Rank | Clausen et al 2013 | Oligo-galacturonates (code and structure) | DP | DM | Σ EL / Σ RMSD | Σ EL | EL = Energy level (kcal/mol) 9 poses of docking vs all surface of PRX36 | | | | | | | | | Σ RMSD | RMSD (Root-mean-square deviation) 9 poses of docking vs all surface of PRX36 | | | | | | | | |
|---|---|---|---|---|---|---|---|---|---|---|---|---|---|---|---|---|---|---|---|---|---|---|---|---|---|
| | | | | | | | 1 | 2 | 3 | 4 | 5 | 6 | 7 | 8 | 9 | | 1 | 2 | 3 | 4 | 5 | 6 | 7 | 8 | 9 |
| 1 | Clausen 4 | 063-o8888o | 6 | 67% | -1.1306 | -51.3 | **-6.7** | **-6.2** | **-5.7** | **-5.6** | **-5.6** | **-5.4** | **-5.4** | -5.4 | **-5.3** | 45.374 | **0.00** | **2.13** | **2.62** | **2.85** | **2.25** | **4.26** | **1.98** | 27.36 | **1.94** |
| 2 | Clausen 3 | 047-oo888o | 6 | 50% | -1.0269 | -53.8 | **-6.7** | **-6.3** | **-6.2** | **-6.1** | **-5.9** | **-5.8** | **-5.7** | **-5.6** | -5.5 | 52.390 | **0.00** | **6.05** | **4.88** | **4.20** | **5.03** | **7.04** | **1.75** | **4.60** | 18.84 |
| 3 | | 038-ooo8o8 | 6 | 33% | -0.8588 | -54.4 | -6.2 | -6.1 | -6.1 | -6.1 | -6.0 | **-6.0** | -6.0 | -6.0 | **-5.9** | 63.341 | 0.00 | 3.74 | 5.06 | 3.26 | 1.76 | **20.64** | 4.23 | 4.99 | **19.66** |
| 4 | | 040-ooo888 | 6 | 50% | -0.8098 | -51.7 | -6.0 | -5.9 | -5.8 | **-5.7** | -5.7 | -5.7 | **-5.7** | -5.6 | -5.6 | 63.839 | 0.00 | 6.51 | 2.99 | **20.97** | 2.35 | 2.93 | **20.10** | 1.57 | 6.42 |
| 5 | | 041-oo8ooo | 6 | 17% | -0.8022 | -54.9 | -6.8 | **-6.4** | -6.2 | **-6.0** | -6.0 | -5.9 | -5.9 | -5.9 | -5.8 | 68.434 | 0.00 | **19.41** | 3.13 | **20.53** | 9.12 | 4.92 | 1.76 | 7.48 | 2.10 |
| 6 | | 024-8oo888 | 6 | 50% | -0.7954 | -50.6 | **-5.7** | -5.7 | **-5.7** | **-5.7** | -5.6 | -5.6 | -5.6 | -5.5 | -5.5 | 63.615 | **0.00** | 21.83 | **4.18** | **2.49** | 15.39 | **3.41** | **2.66** | **2.22** | 11.44 |
| 7 | | 106-ooo8 | 4 | **25%** | -0.7944 | -49.7 | **-5.9** | **-5.7** | **-5.7** | **-5.5** | **-5.5** | **-5.4** | -5.4 | -5.3 | **-5.3** | 62.56 | **0.00** | **4.09** | **6.78** | **2.83** | **1.99** | **2.66** | 16.15 | 23.98 | **4.09** |
| 8 | | 069-88ooo | 5 | 40% | -0.7804 | -62.4 | -7.2 | **-7.1** | -7.1 | **-7.1** | -7.0 | **-6.8** | -6.7 | -6.7 | -6.7 | 79.96 | 0.00 | **19.49** | 9.15 | **18.24** | 5.91 | **18.59** | 2.26 | 3.83 | 2.49 |
| 9 | | 008-888o88 | 6 | 83% | -0.7742 | -58.3 | -6.7 | -6.7 | -6.6 | -6.5 | **-6.4** | -6.4 | -6.4 | -6.3 | **-6.3** | 75.30 | 0.00 | 0.51 | 1.61 | 12.10 | **17.42** | 8.63 | 8.07 | 10.57 | **16.40** |
| 10 | | 065-88888 | 5 | 100% | -0.7498 | -51.5 | -5.9 | -5.8 | -5.8 | -5.7 | -5.7 | **-5.7** | **-5.7** | **-5.6** | **-5.6** | 68.68 | 0.00 | 2.13 | 5.57 | 2.41 | 1.94 | **15.26** | **11.17** | **16.66** | **13.56** |
| 11 | | 107-oo8o | 4 | **25%** | -0.7421 | -49.8 | **-5.7** | -5.6 | **-5.6** | **-5.6** | **-5.5** | **-5.5** | -5.5 | **-5.4** | **-5.4** | 67.11 | **0.00** | 17.69 | **7.42** | **7.22** | **2.83** | **3.67** | 17.19 | **7.77** | **3.33** |
| 12 | | 101-8ooo | 4 | 25% | -0.7365 | -51.2 | -6.4 | **-5.9** | **-5.8** | -5.7 | -5.5 | **-5.5** | -5.5 | -5.5 | **-5.4** | 69.52 | 0.00 | **15.52** | **15.52** | 3.18 | 2.36 | **15.06** | 2.17 | 1.73 | **13.98** |
| 13 | | 099-88oo | 4 | 50% | -0.6895 | -52.9 | -6.5 | -6.3 | -5.9 | **-5.8** | **-5.8** | -5.7 | **-5.7** | **-5.7** | -5.5 | 76.73 | 0.00 | 3.02 | 3.84 | **14.30** | **15.55** | 4.10 | **15.57** | **15.61** | 4.74 |
| 14 | | 096-o8888 | 5 | 80% | -0.6427 | -52.0 | -6.1 | -5.9 | -5.8 | **-5.7** | **-5.7** | -5.7 | **-5.7** | -5.7 | **-5.7** | 80.91 | 0.00 | 3.48 | 4.72 | **14.74** | **16.89** | 4.48 | **15.79** | 4.00 | **16.81** |
| 15 | | 023-8oo88o | 6 | 50% | -0.5895 | -54.6 | -7.0 | **-6.3** | **-6.1** | **-5.9** | -5.9 | -5.9 | -5.9 | **-5.8** | -5.8 | 92.63 | 0.00 | **8.55** | **9.69** | **8.92** | 2.04 | 21.14 | 18.82 | **7.62** | 15.86 |
| 16 | Clausen 5 | 027-8o8o8o | 6 | 50% | -0.5806 | -50.9 | **-6.3** | **-6.1** | **-5.8** | **-5.8** | **-5.7** | **-5.4** | -5.3 | -5.3 | -5.2 | 87.67 | **0.00** | **2.58** | **2.41** | **3.03** | **2.97** | **4.85** | 25.48 | 14.61 | 31.75 |
| 17 | | 028-8o8o88 | 6 | 67% | -0.5750 | -54.5 | -6.7 | -6.4 | -6.1 | -6.0 | -5.9 | -5.9 | **-5.9** | **-5.8** | **-5.8** | 94.78 | 0.00 | 1.72 | 3.46 | 1.86 | 11.11 | 12.55 | **22.19** | **20.89** | **20.99** |
| 18 | | 009-88oooo | 6 | 33% | -0.5662 | -54.2 | -6.4 | -6.4 | -6.2 | -6.0 | **-5.9** | -5.9 | -5.8 | -5.8 | -5.8 | 95.72 | 0.00 | 21.24 | 3.50 | 21.34 | **11.17** | 21.66 | 3.27 | 8.83 | 4.71 |

| | | | | | | | | | | | | | | | | | | | | | | | | | |
|---|---|---|---|---|---|---|---|---|---|---|---|---|---|---|---|---|---|---|---|---|---|---|---|---|---|
| 19 | | 012-88o088 | 6 | 33% | -0.5633 | -48.4 | -5.6 | -5.6 | -5.4 | -5.4 | -5.4 | -5.4 | -5.3 | -5.2 | -5.1 | 85.93 | 0.00 | 7.63 | 26.74 | 1.81 | 1.83 | 1.86 | 11.13 | 27.92 | 7.02 |
| 20 | | 059-o88o8o | 6 | 50% | -0.5596 | -54.4 | -6.5 | -6.2 | -6.2 | -6.1 | -6.1 | -5.9 | -5.8 | -5.8 | -5.8 | 97.21 | 0.00 | 3.11 | 5.21 | 20.24 | 20.24 | 21.47 | 22.00 | 1.77 | 3.18 |
| 21 | | 108-oo88 | 4 | 50% | -0.5584 | -51.8 | -6.7 | -5.8 | -5.7 | -5.7 | -5.7 | -5.6 | -5.6 | -5.6 | -5.4 | 92.76 | 0.00 | 4.90 | 5.54 | 17.32 | 5.07 | 17.01 | 12.68 | 17.56 | 12.67 |
| 22 | | 060-o88o88 | 6 | 67% | -0.5569 | -51.5 | -5.9 | -5.8 | -5.8 | -5.7 | -5.7 | -5.7 | -5.7 | -5.6 | -5.6 | 92.48 | 0.00 | 6.52 | 3.61 | 26.08 | 2.06 | 3.48 | 18.50 | 25.88 | 6.35 |
| 23 | | 112-o888 | 4 | 75% | -0.5485 | -51.5 | -6.5 | -5.8 | -5.8 | -5.8 | -5.7 | -5.5 | -5.5 | -5.5 | -5.4 | 93.90 | 0.00 | 16.72 | 3.27 | 14.42 | 1.58 | 3.23 | 24.93 | 14.47 | 15.28 |
| 24 | | 097-8888 | 4 | 100% | -0.5415 | -51.7 | -6.5 | -5.8 | -5.8 | -5.8 | -5.7 | -5.6 | -5.5 | -5.5 | -5.5 | 95.47 | 0.00 | 3.44 | 1.66 | 1.66 | 15.42 | 15.75 | 15.30 | 14.67 | 27.58 |
| 25 | | 064-o88888 | 6 | 83% | -0.5366 | -52.9 | -6.6 | -6.2 | -5.9 | -5.8 | -5.7 | -5.7 | -5.7 | -5.7 | -5.6 | 98.58 | 0.00 | 3.09 | 2.34 | 3.46 | 20.90 | 21.42 | 5.16 | 20.56 | 21.65 |
| 26 | | 031-8o888o | 6 | 67% | -0.5304 | -54.6 | -6.8 | -6.2 | -6.1 | -6.0 | -6.0 | -5.9 | -5.9 | -5.9 | -5.8 | 102.94 | 0.00 | 21.38 | 13.80 | 6.25 | 21.46 | 12.06 | 1.77 | 19.83 | 6.40 |
| 27 | | 015-88o88o | 6 | 67% | -0.5291 | -53.6 | -6.8 | -6.0 | -5.9 | -5.9 | -5.9 | -5.9 | -5.8 | -5.7 | -5.7 | 101.30 | 0.00 | 1.24 | 5.77 | 5.08 | 20.34 | 19.86 | 21.25 | 13.46 | 14.30 |
| 28 | | 089-o8ooo | 5 | 20% | -0.5258 | -48.7 | -5.9 | -5.6 | -5.5 | -5.4 | -5.3 | -5.3 | -5.3 | -5.2 | -5.2 | 92.63 | 0.00 | 25.06 | 4.10 | 11.43 | 1.37 | 10.61 | 25.04 | 9.99 | 5.02 |
| 29 | | 037-ooo8oo | 6 | 17% | -0.5218 | -55.6 | -7.0 | -6.5 | -6.2 | -6.2 | -6.1 | -6.0 | -5.9 | -5.9 | -5.8 | 106.55 | 0.00 | 1.61 | 2.90 | 21.13 | 13.80 | 4.44 | 21.03 | 20.45 | 21.20 |
| 30 | | 093-o88oo | 5 | 40% | -0.5175 | -50.5 | -5.8 | -5.7 | -5.7 | -5.7 | -5.6 | -5.5 | -5.5 | -5.5 | -5.5 | 97.59 | 0.00 | 11.20 | 6.85 | 10.54 | 1.38 | 6.32 | 11.08 | 25.14 | 25.08 |
| 31 | | 075-8oo8o | 5 | 40% | -0.5172 | -53.9 | -6.4 | -6.3 | -6.1 | -6.0 | -6.0 | -5.9 | -5.8 | -5.7 | -5.7 | 104.21 | 0.00 | 5.06 | 6.26 | 3.79 | 20.87 | 5.07 | 20.24 | 30.76 | 12.15 |
| 32 | | 029-8o88oo | 6 | 50% | -0.5124 | -54.8 | -7.0 | -6.2 | -6.1 | -6.0 | -6.0 | -5.9 | -5.9 | -5.9 | -5.8 | 106.96 | 0.00 | 1.59 | 2.07 | 11.61 | 21.19 | 25.43 | 2.35 | 21.00 | 21.72 |
| 33 | | 051-o8oo8o | 6 | 33% | -0.5093 | -56.4 | -6.7 | -6.5 | -6.5 | -6.3 | -6.2 | -6.2 | -6.0 | -6.0 | -6.0 | 110.73 | 0.00 | 8.64 | 8.21 | 6.03 | 1.66 | 31.52 | 21.20 | 12.54 | 20.93 |
| 34 | | 007-888o8o | 6 | 67% | -0.5085 | -53.8 | -6.6 | -6.3 | -6.0 | -5.9 | -5.9 | -5.8 | -5.8 | -5.8 | -5.7 | 105.80 | 0.00 | 1.80 | 1.74 | 5.49 | 20.29 | 11.85 | 21.14 | 21.33 | 22.17 |
| 35 | | 049-o8oooo | 6 | 17% | -0.5047 | -57.8 | -6.8 | -6.8 | -6.6 | -6.5 | -6.3 | -6.3 | -6.2 | -6.2 | -6.1 | 114.52 | 0.00 | 11.27 | 11.06 | 1.71 | 12.66 | 25.87 | 24.95 | 2.75 | 24.25 |
| 36 | | 061-o888oo | 6 | 50% | -0.4987 | -53.6 | -6.6 | -6.3 | -5.9 | -5.9 | -5.9 | -5.8 | -5.8 | -5.7 | -5.7 | 107.47 | 0.00 | 3.11 | 21.94 | 14.05 | 22.31 | 4.30 | 20.18 | 8.91 | 12.67 |
| 37 | | 044-oo8o88 | 6 | 50% | -0.4965 | -54.7 | -6.8 | -6.6 | -6.0 | -5.9 | -5.9 | -5.9 | -5.9 | -5.9 | -5.8 | 110.16 | 0.00 | 1.65 | 5.47 | 11.60 | 20.45 | 34.00 | 20.99 | 14.14 | 1.87 |
| 38 | | 050-o8ooo8 | 6 | 33% | -0.4929 | -52.8 | -6.1 | -6.0 | -6.0 | -5.9 | -5.8 | -5.8 | -5.8 | -5.7 | -5.7 | 107.13 | 0.00 | 25.36 | 24.45 | 14.90 | 2.05 | 14.84 | 1.64 | 6.71 | 17.18 |
| 39 | | 052-o8oo88 | 6 | 50% | -0.4861 | -52.5 | -6.1 | -5.9 | -5.9 | -5.8 | -5.8 | -5.8 | -5.8 | -5.7 | -5.7 | 108.01 | 0.00 | 21.22 | 5.15 | 4.51 | 5.62 | 30.04 | 29.19 | 6.67 | 5.60 |
| 40 | | 016-88o888 | 6 | 83% | -0.4778 | -51.5 | -6.7 | -5.8 | -5.7 | -5.6 | -5.6 | -5.6 | -5.6 | -5.5 | -5.4 | 107.78 | 0.00 | 6.19 | 3.55 | 21.40 | 22.74 | 11.81 | 4.08 | 29.24 | 8.77 |
| 41 | | 039-ooo88o | 6 | 33% | -0.4730 | -56.2 | -6.4 | -6.3 | -6.3 | -6.3 | -6.2 | -6.2 | -6.2 | -6.2 | -6.1 | 118.81 | 0.00 | 24.80 | 6.47 | 20.28 | 11.98 | 24.25 | 1.57 | 24.03 | 5.42 |
| 42 | | 062-o888o8 | 6 | 67% | -0.4719 | -54.1 | -6.5 | -6.1 | -6.1 | -6.0 | -5.9 | -5.9 | -5.9 | -5.9 | -5.8 | 114.64 | 0.00 | 6.58 | 5.79 | 2.73 | 21.78 | 5.24 | 21.36 | 29.20 | 21.96 |
| 43 | | 013-88o8oo | 6 | 50% | -0.4711 | -53.9 | -6.7 | -6.0 | -6.0 | -5.9 | -5.9 | -5.9 | -5.9 | -5.8 | -5.8 | 114.40 | 0.00 | 20.85 | 20.88 | 6.11 | 32.21 | 21.21 | 4.58 | 3.02 | 5.56 |
| 44 | | 105-oooo | 4 | 0% | -0.4702 | -53.8 | -6.6 | -6.1 | -6.0 | -6.0 | -5.9 | -5.9 | -5.8 | -5.8 | -5.7 | 114.43 | 0.00 | 14.38 | 15.32 | 4.12 | 15.29 | 15.40 | 20.67 | 14.92 | 14.34 |
| 45 | | 036-ooo088 | 6 | 33% | -0.4697 | -53.2 | -6.1 | -6.0 | -6.0 | -5.9 | -5.9 | -5.9 | -5.8 | -5.8 | -5.8 | 113.27 | 0.00 | 22.32 | 12.31 | 17.06 | 23.59 | 13.26 | 12.23 | 7.27 | 5.24 |
| 46 | | 003-8888oo | 6 | 67% | -0.4682 | -54.7 | -6.8 | -6.3 | -6.2 | -6.0 | -6.0 | -5.9 | -5.9 | -5.8 | -5.8 | 116.83 | 0.00 | 3.23 | 1.69 | 21.00 | 19.27 | 5.32 | 14.67 | 20.44 | 31.21 |
| 47 | | 048-oo8888 | 6 | 67% | -0.4663 | -56.3 | -6.8 | -6.7 | -6.2 | -6.2 | -6.2 | -6.1 | -6.1 | -6.0 | -6.0 | 120.74 | 0.00 | 1.69 | 20.64 | 21.06 | 1.20 | 20.97 | 21.75 | 20.26 | 13.17 |

| | | | | | | | | | | | | | | | | | | | | | | | | | |
|---|---|---|---|---|---|---|---|---|---|---|---|---|---|---|---|---|---|---|---|---|---|---|---|---|---|
| 48 | | 114-88o | 3 | 67% | -0.4578 | -48.3 | -5.9 | -5.7 | -5.4 | -5.3 | -5.3 | -5.3 | -5.2 | -5.1 | -5.1 | 105.50 | 0.00 | 26.87 | 10.33 | 3.09 | 1.40 | 32.61 | 3.48 | 25.98 | 1.74 |
| 49 | | 068-888o8 | 5 | 80% | -0.4552 | -49.4 | -5.6 | -5.6 | -5.5 | -5.5 | -5.5 | -5.5 | -5.4 | -5.4 | -5.4 | 108.52 | 0.00 | 11.32 | 11.34 | 1.95 | 25.09 | 25.10 | 30.14 | 1.41 | 2.18 |
| 50 | | 082-oooo8 | 5 | 20% | -0.4543 | -49.2 | -5.7 | -5.7 | -5.6 | -5.5 | -5.4 | -5.4 | -5.3 | -5.3 | -5.3 | 108.31 | 0.00 | 1.02 | 2.52 | 24.77 | 10.83 | 1.63 | 10.48 | 32.34 | 24.72 |
| 51 | | 033-oooooo | 6 | 0% | -0.4509 | -60.3 | -7.0 | -6.9 | -6.8 | -6.8 | -6.8 | -6.6 | -6.5 | -6.5 | -6.4 | 133.74 | 0.00 | 2.89 | 21.53 | 32.23 | 12.16 | 14.92 | 22.69 | 5.14 | 22.19 |
| 52 | | 006-888oo8 | 6 | 67% | -0.4508 | -51.7 | -6.1 | -5.9 | -5.8 | -5.7 | -5.7 | -5.7 | -5.6 | -5.6 | -5.6 | 114.67 | 0.00 | 11.77 | 23.48 | 2.99 | 26.64 | 28.46 | 1.71 | 2.03 | 17.59 |
| 53 | | 081-ooooo | 5 | 0% | -0.4435 | -55.1 | -6.3 | -6.2 | -6.2 | -6.2 | -6.2 | -6.0 | -6.0 | -6.0 | -6.0 | 124.25 | 0.00 | 13.79 | 1.34 | 5.11 | 1.78 | 25.34 | 25.63 | 25.91 | 25.35 |
| 54 | | 021-8oo8oo | 6 | 33% | -0.4388 | -55.3 | -6.9 | -6.3 | -6.2 | -6.1 | -6.0 | -6.0 | -6.0 | -5.9 | -5.9 | 126.02 | 0.00 | 6.23 | 20.93 | 3.69 | 6.08 | 31.83 | 20.92 | 5.66 | 30.70 |
| 55 | | 014-88o8o8 | 6 | 67% | -0.4382 | -54.1 | -6.8 | -6.3 | -6.1 | -6.0 | -6.0 | -5.8 | -5.8 | -5.7 | -5.6 | 123.46 | 0.00 | 7.60 | 21.70 | 4.04 | 21.49 | 11.62 | 12.12 | 22.06 | 22.83 |
| 56 | | 076-8oo88 | 5 | 60% | -0.4360 | -48.7 | -5.9 | -5.6 | -5.5 | -5.4 | -5.3 | -5.3 | -5.3 | -5.2 | -5.2 | 111.70 | 0.00 | 5.89 | 7.04 | 25.20 | 25.16 | 1.44 | 10.62 | 30.88 | 5.47 |
| 57 | | 070-88oo8 | 5 | 60% | -0.4342 | -49.5 | -5.7 | -5.6 | -5.5 | -5.5 | -5.5 | -5.5 | -5.4 | -5.4 | -5.4 | 113.99 | 0.00 | 10.88 | 3.95 | 30.97 | 2.76 | 24.26 | 10.90 | 5.79 | 24.50 |
| 58 | | 042-oo8oo8 | 6 | 33% | -0.4325 | -55.9 | -6.8 | -6.4 | -6.2 | -6.1 | -6.1 | -6.1 | -6.1 | -6.1 | -6.0 | 129.23 | 0.00 | 11.75 | 19.87 | 21.03 | 21.17 | 1.84 | 12.87 | 20.40 | 20.30 |
| 59 | | 055-o8o88o | 6 | 50% | -0.4253 | -54.9 | -6.6 | -6.2 | -6.1 | -6.0 | -6.0 | -6.0 | -6.0 | -6.0 | -6.0 | 129.07 | 0.00 | 5.90 | 11.80 | 3.64 | 20.30 | 22.22 | 21.43 | 22.04 | 21.74 |
| 60 | | 085-oo8oo | 5 | 20% | -0.4246 | -51.8 | -5.9 | -5.8 | -5.8 | -5.8 | -5.8 | -5.7 | -5.7 | -5.7 | -5.6 | 122.01 | 0.00 | 26.40 | 26.24 | 15.79 | 3.65 | 16.06 | 26.73 | 1.55 | 5.60 |
| 61 | | 054-o8o8o8 | 6 | 50% | -0.4214 | -54.3 | -6.5 | -6.1 | -6.0 | -6.0 | -6.0 | -6.0 | -5.9 | -5.9 | -5.9 | 128.85 | 0.00 | 2.97 | 8.87 | 21.78 | 22.44 | 23.17 | 24.11 | 21.32 | 4.21 |
| 62 | | 011-88oo8o | 6 | 50% | -0.4206 | -55.4 | -6.5 | -6.3 | -6.3 | -6.2 | -6.1 | -6.0 | -6.0 | -6.0 | -6.0 | 131.70 | 0.00 | 31.56 | 21.44 | 21.48 | 21.19 | 4.10 | 20.26 | 8.48 | 3.21 |
| 63 | | 002-88888o | 6 | 67% | -0.4200 | -52.7 | -6.7 | -6.1 | -5.9 | -5.9 | -5.8 | -5.7 | -5.6 | -5.5 | -5.5 | 125.49 | 0.00 | 3.04 | 19.07 | 3.00 | 8.36 | 20.12 | 31.68 | 19.94 | 20.28 |
| 64 | | 034-ooooo8 | 6 | 17% | -0.4197 | -51.3 | -5.8 | -5.8 | -5.8 | -5.7 | -5.7 | -5.7 | -5.6 | -5.6 | -5.6 | 122.24 | 0.00 | 27.17 | 16.78 | 28.87 | 13.51 | 2.34 | 2.72 | 28.67 | 2.19 |
| 65 | | 078-8o8o8 | 5 | 60% | -0.4165 | -51.9 | -5.9 | -5.9 | -5.9 | -5.8 | -5.8 | -5.7 | -5.7 | -5.6 | -5.6 | 124.61 | 0.00 | 15.10 | 9.12 | 22.44 | 1.61 | 28.94 | 1.73 | 23.24 | 22.42 |
| 66 | | 030-8o88o8 | 6 | 67% | -0.4139 | -50.9 | -5.9 | -5.9 | -5.7 | -5.6 | -5.6 | -5.6 | -5.6 | -5.5 | -5.5 | 122.97 | 0.00 | 11.16 | 26.63 | 10.87 | 1.77 | 12.01 | 28.25 | 4.07 | 28.21 |
| 67 | | 102-8oo8 | 4 | 50% | -0.4091 | -48.5 | -5.8 | -5.6 | -5.4 | -5.4 | -5.4 | -5.3 | -5.3 | -5.2 | -5.1 | 118.55 | 0.00 | 14.92 | 1.82 | 24.67 | 32.21 | 32.31 | 1.69 | 5.23 | 5.71 |
| 68 | | 109-o8oo | 4 | 25% | -0.4085 | -51.5 | -6.5 | -6.0 | -5.9 | -5.8 | -5.7 | -5.6 | -5.4 | -5.3 | -5.3 | 126.07 | 0.00 | 1.45 | 3.12 | 20.13 | 14.24 | 28.62 | 26.38 | 14.29 | 17.86 |
| 69 | | 045-oo88oo | 6 | 33% | -0.4052 | -55.0 | -6.9 | -6.2 | -6.1 | -6.0 | -6.0 | -6.0 | -6.0 | -5.9 | -5.9 | 135.72 | 0.00 | 1.57 | 22.21 | 11.35 | 14.20 | 20.52 | 23.00 | 21.02 | 21.85 |
| 70 | | 115-8oo | 3 | 33% | -0.4046 | -51.5 | -6.4 | -6.3 | -5.9 | -5.7 | -5.5 | -5.5 | -5.5 | -5.4 | -5.3 | 127.29 | 0.00 | 1.33 | 13.92 | 3.57 | 25.05 | 11.75 | 23.03 | 23.18 | 25.46 |
| 71 | | 019-8ooo8o | 6 | 33% | -0.4046 | -55.9 | -7.0 | -6.3 | -6.2 | -6.1 | -6.1 | -6.1 | -6.1 | -6.0 | -6.0 | 138.18 | 0.00 | 3.29 | 30.95 | 21.66 | 2.05 | 31.59 | 20.85 | 21.87 | 5.92 |
| 72 | | 090-o8oo8 | 5 | 40% | -0.4041 | -49.8 | -5.9 | -5.8 | -5.8 | -5.5 | -5.5 | -5.4 | -5.3 | -5.3 | -5.3 | 123.25 | 0.00 | 30.52 | 25.25 | 24.96 | 8.85 | 8.45 | 9.62 | 8.08 | 7.51 |
| 73 | | 025-8o8ooo | 6 | 33% | -0.4014 | -54.1 | -6.8 | -6.4 | -6.0 | -5.9 | -5.9 | -5.9 | -5.8 | -5.7 | -5.7 | 134.78 | 0.00 | 3.29 | 31.55 | 22.53 | 5.47 | 23.32 | 20.54 | 21.13 | 6.95 |
| 74 | | 092-o8o88 | 5 | 60% | -0.4012 | -49.7 | -5.7 | -5.6 | -5.6 | -5.6 | -5.5 | -5.5 | -5.4 | -5.4 | -5.4 | 123.87 | 0.00 | 3.99 | 14.85 | 14.56 | 5.42 | 24.33 | 31.36 | 5.32 | 24.06 |
| 75 | | 057-o88ooo | 6 | 33% | -0.4010 | -51.8 | -6.0 | -5.9 | -5.9 | -5.8 | -5.8 | -5.6 | -5.6 | -5.6 | -5.6 | 129.18 | 0.00 | 24.35 | 24.43 | 24.07 | 24.29 | 12.22 | 5.06 | 1.85 | 12.92 |
| 76 | | 005-888ooo | 6 | 50% | -0.3864 | -52.2 | -6.3 | -6.1 | -5.9 | -5.8 | -5.7 | -5.6 | -5.6 | -5.6 | -5.6 | 135.08 | 0.00 | 21.63 | 21.60 | 3.07 | 11.25 | 18.52 | 31.32 | 12.51 | 15.19 |

| | | | | | | | | | | | | | | | | | | | | | | | | | |
|---|---|---|---|---|---|---|---|---|---|---|---|---|---|---|---|---|---|---|---|---|---|---|---|---|---|
| 77 | | 058-o88oo8 | 6 | 50% | -0.3817 | -56.0 | -6.4 | -6.4 | -6.3 | -6.2 | -6.2 | -6.2 | -6.2 | -6.1 | -6.0 | 146.73 | 0.00 | 27.21 | 25.33 | 4.39 | 13.96 | 25.32 | 2.02 | 23.18 | 25.32 |
| 78 | | 095-o888o | 5 | 60% | -0.3807 | -50.2 | -5.8 | -5.8 | -5.6 | -5.6 | -5.6 | -5.5 | -5.5 | -5.4 | -5.4 | 131.85 | 0.00 | 7.69 | 7.38 | 11.98 | 20.97 | 23.48 | 21.65 | 35.96 | 2.74 |
| 79 | | 080 - 8o888 | 5 | 80% | -0.3778 | -52.2 | -6.1 | -6.0 | -5.9 | -5.8 | -5.8 | -5.7 | -5.7 | -5.6 | -5.6 | 138.15 | 0.00 | 25.61 | 24.34 | 5.49 | 1.77 | 24.05 | 15.16 | 25.50 | 16.23 |
| 80 | | 017-8ooooo | 6 | 17% | -0.3754 | -53.3 | -6.2 | -6.1 | -6.1 | -6.0 | -5.9 | -5.8 | -5.8 | -5.7 | -5.7 | 141.97 | 0.00 | 8.51 | 14.95 | 23.72 | 15.89 | 22.56 | 23.70 | 10.37 | 22.28 |
| 81 | | 022-8oo8o8 | 6 | 50% | -0.3721 | -52.5 | -6.8 | -6.0 | -5.8 | -5.7 | -5.7 | -5.7 | -5.6 | -5.6 | -5.6 | 141.09 | 0.00 | 5.24 | 20.48 | 18.84 | 19.52 | 20.56 | 3.41 | 21.11 | 31.95 |
| 82 | | 066-8888o | 5 | 80% | -0.3718 | -53.6 | -6.3 | -6.0 | -6.0 | -6.0 | -6.0 | -5.9 | -5.8 | -5.8 | -5.8 | 144.16 | 0.00 | 5.01 | 22.36 | 23.77 | 21.07 | 18.42 | 23.64 | 23.95 | 5.93 |
| 83 | | 111-o88o | 4 | 50% | -0.3692 | -48.9 | -5.8 | -5.6 | -5.5 | -5.5 | -5.4 | -5.3 | -5.3 | -5.3 | -5.2 | 132.45 | 0.00 | 14.72 | 14.75 | 34.39 | 9.55 | 15.27 | 22.02 | 2.76 | 18.99 |
| 84 | | 087-oo88o | 5 | 40% | -0.3655 | -50.0 | -5.7 | -5.7 | -5.6 | -5.5 | -5.5 | -5.5 | -5.5 | -5.5 | -5.5 | 136.81 | 0.00 | 7.39 | 15.02 | 25.59 | 5.45 | 15.36 | 27.59 | 12.90 | 27.53 |
| 85 | | 083-ooo8o | 5 | 20% | -0.3644 | -52.2 | -6.0 | -5.9 | -5.9 | -5.8 | -5.8 | -5.8 | -5.7 | -5.7 | -5.6 | 143.25 | 0.00 | 25.41 | 1.79 | 20.69 | 12.40 | 17.52 | 15.80 | 16.63 | 33.01 |
| 86 | | 074-8ooo8 | 5 | 40% | -0.3610 | -48.9 | -5.7 | -5.6 | -5.6 | -5.5 | -5.5 | -5.4 | -5.4 | -5.1 | -5.1 | 135.46 | 0.00 | 1.80 | 15.11 | 2.70 | 2.12 | 26.52 | 34.85 | 26.67 | 25.71 |
| 87 | | 067-888oo | 5 | 60% | -0.3607 | -54.7 | -6.4 | -6.3 | -6.2 | -6.1 | -6.0 | -6.0 | -5.9 | -5.9 | -5.9 | 151.66 | 0.00 | 22.89 | 30.59 | 24.53 | 15.13 | 14.97 | 31.48 | 1.25 | 10.83 |
| 88 | | 056-o8o888 | 6 | 67% | -0.3572 | -53.8 | -6.1 | -6.1 | -6.0 | -6.0 | -6.0 | -6.0 | -5.9 | -5.9 | -5.8 | 150.60 | 0.00 | 25.16 | 14.54 | 26.90 | 1.63 | 26.46 | 24.08 | 6.06 | 25.77 |
| 89 | | 046-oo88o8 | 6 | 50% | -0.3544 | -54.7 | -6.8 | -6.3 | -6.1 | -6.0 | -5.9 | -5.9 | -5.9 | -5.9 | -5.9 | 154.35 | 0.00 | 21.38 | 21.13 | 6.75 | 20.38 | 19.81 | 21.72 | 21.08 | 22.11 |
| 90 | | 020-8ooo88 | 6 | 50% | -0.3528 | -53.6 | -6.4 | -6.1 | -6.0 | -6.0 | -5.9 | -5.8 | -5.8 | -5.8 | -5.8 | 151.91 | 0.00 | 1.75 | 1.16 | 24.26 | 25.76 | 24.26 | 24.72 | 24.85 | 25.16 |
| 91 | Clausen 2 | 010-88ooo8 | 6 | 50% | -0.3528 | -49.4 | -6.1 | -5.8 | -5.7 | -5.6 | -5.3 | -5.3 | -5.3 | -5.2 | -5.1 | 140.03 | 0.00 | 32.68 | 23.11 | 2.37 | 32.30 | 14.93 | 10.36 | 5.33 | 18.96 |
| 92 | | 043-oo8o8o | 6 | 33% | -0.3515 | -54.8 | -6.2 | -6.2 | -6.2 | -6.1 | -6.1 | -6.1 | -6.0 | -6.0 | -5.9 | 155.91 | 0.00 | 23.81 | 27.20 | 27.36 | 23.24 | 25.20 | 1.94 | 3.69 | 23.50 |
| 93 | | 032-8o8888 | 6 | 83% | -0.3506 | -52.5 | -6.7 | -5.8 | -5.8 | -5.7 | -5.7 | -5.7 | -5.7 | -5.7 | -5.7 | 149.75 | 0.00 | 6.12 | 21.21 | 31.92 | 3.27 | 21.31 | 5.23 | 30.80 | 29.91 |
| 94 | | 072-88o88 | 5 | 80% | -0.3484 | -48.6 | -5.9 | -5.8 | -5.5 | -5.3 | -5.3 | -5.3 | -5.2 | -5.2 | -5.1 | 139.51 | 0.00 | 11.11 | 5.18 | 11.60 | 10.17 | 21.59 | 32.54 | 24.50 | 22.84 |
| 95 | | 122-8o | 2 | 50% | -0.3461 | -48.8 | -6.1 | -5.8 | -5.5 | -5.4 | -5.3 | -5.3 | -5.2 | -5.1 | -5.1 | 140.99 | 0.00 | 2.90 | 2.91 | 14.85 | 30.71 | 16.53 | 16.98 | 31.32 | 24.80 |
| 96 | | 103-8o8o | 4 | 50% | -0.3442 | -51.9 | -6.4 | -6.0 | -5.8 | -5.8 | -5.7 | -5.6 | -5.6 | -5.5 | -5.5 | 150.78 | 0.00 | 25.54 | 10.03 | 24.51 | 3.33 | 25.49 | 11.47 | 17.79 | 32.64 |
| 97 | | 124-o8 | 2 | 50% | -0.3438 | -50.4 | -6.2 | -6.1 | -5.8 | -5.6 | -5.4 | -5.4 | -5.3 | -5.3 | -5.3 | 146.60 | 0.00 | 47.24 | 12.82 | 12.54 | 24.50 | 1.33 | 13.16 | 23.84 | 11.17 |
| 98 | | 113-888 | 3 | 100% | -0.3401 | -45.4 | -5.6 | -5.2 | -5.1 | -5.0 | -5.0 | -5.0 | -5.0 | -4.8 | -4.7 | 133.47 | 0.00 | 3.17 | 3.08 | 3.41 | 29.64 | 30.83 | 29.94 | 3.38 | 30.03 |
| 99 | | 035-oooo8o | 6 | 17% | -0.3349 | -57.3 | -6.6 | -6.5 | -6.5 | -6.4 | -6.4 | -6.4 | -6.2 | -6.2 | -6.1 | 171.10 | 0.00 | 11.35 | 25.39 | 24.74 | 21.06 | 14.74 | 23.67 | 24.48 | 25.67 |
| 100 | | 110-o8o8 | 4 | 50% | -0.3316 | -48.9 | -5.7 | -5.5 | -5.5 | -5.5 | -5.4 | -5.4 | -5.3 | -5.3 | -5.3 | 147.45 | 0.00 | 14.78 | 32.44 | 28.20 | 15.67 | 5.33 | 22.74 | 23.30 | 4.99 |
| 101 | | 100-88o8 | 4 | 75% | -0.3298 | -48.3 | -5.7 | -5.5 | -5.4 | -5.4 | -5.4 | -5.3 | -5.2 | -5.2 | -5.2 | 146.44 | 0.00 | 3.01 | 15.71 | 15.61 | 28.20 | 29.83 | 11.01 | 15.25 | 27.81 |
| 102 | | 084-ooo88 | 5 | 40% | -0.3277 | -50.6 | -6.1 | -5.8 | -5.8 | -5.6 | -5.5 | -5.5 | -5.5 | -5.5 | -5.3 | 154.40 | 0.00 | 16.99 | 17.56 | 1.47 | 27.17 | 7.39 | 24.51 | 28.23 | 31.08 |
| 103 | | 026-8o8oo8 | 6 | 50% | -0.3168 | -49.2 | -5.8 | -5.7 | -5.7 | -5.5 | -5.4 | -5.4 | -5.3 | -5.2 | -5.2 | 155.28 | 0.00 | 5.68 | 31.34 | 20.20 | 29.67 | 22.10 | 4.93 | 25.01 | 16.36 |
| 104 | | 077-8o8oo | 5 | 40% | -0.3159 | -57.8 | -6.7 | -6.7 | -6.5 | -6.4 | -6.4 | -6.3 | -6.3 | -6.3 | -6.2 | 182.98 | 0.00 | 17.34 | 23.09 | 24.11 | 31.12 | 30.36 | 24.84 | 14.55 | 17.58 |
| 105 | | 088-oo888 | 5 | 60% | -0.3158 | -50.2 | -5.6 | -5.6 | -5.6 | -5.6 | -5.6 | -5.6 | -5.6 | -5.5 | -5.5 | 158.97 | 0.00 | 21.31 | 41.43 | 17.44 | 15.93 | 28.79 | 29.05 | 1.59 | 3.45 |

| | | | | | | | | | | | | | | | | | | | | | | | | | |
|---|---|---|---|---|---|---|---|---|---|---|---|---|---|---|---|---|---|---|---|---|---|---|---|---|---|
| 106 | | 094-o88o8 | 5 | 60% | -0.3140 | -51.0 | -6.1 | -5.9 | -5.7 | -5.6 | -5.6 | -5.6 | -5.5 | -5.5 | -5.5 | 162.41 | 0.00 | 20.65 | 16.33 | 21.63 | 13.52 | 20.61 | 34.56 | 21.13 | 13.99 |
| 107 | | 104-8o88 | 4 | 75% | -0.3125 | -48.0 | -5.8 | -5.6 | -5.4 | -5.4 | -5.3 | -5.2 | -5.2 | -5.1 | -5.0 | 153.61 | 0.00 | 3.03 | 15.07 | 20.49 | 20.30 | 15.90 | 33.89 | 15.79 | 29.15 |
| 108 | | 004-8888o8 | 6 | 83% | -0.3121 | -50.6 | -6.0 | -5.7 | -5.7 | -5.7 | -5.6 | -5.6 | -5.5 | -5.4 | -5.4 | 162.10 | 0.00 | 24.14 | 24.58 | 17.00 | 28.74 | 32.70 | 12.88 | 6.53 | 15.52 |
| 109 | | 079-8o88o | 5 | 60% | -0.3040 | -50.9 | -5.8 | -5.7 | -5.7 | -5.7 | -5.6 | -5.6 | -5.6 | -5.6 | -5.6 | 167.44 | 0.00 | 3.98 | 19.78 | 20.76 | 7.62 | 20.85 | 28.56 | 29.91 | 35.98 |
| 110 | | 091-o8o8o | 5 | 40% | -0.3014 | -48.9 | -5.8 | -5.6 | -5.5 | -5.4 | -5.4 | -5.3 | -5.3 | -5.3 | -5.3 | 162.24 | 0.00 | 1.47 | 25.46 | 22.66 | 24.37 | 34.68 | 25.35 | 3.75 | 24.50 |
| 111 | | 071-88o8o | 5 | 60% | -0.2991 | -49.1 | -5.8 | -5.7 | -5.5 | -5.5 | -5.4 | -5.4 | -5.3 | -5.3 | -5.2 | 164.14 | 0.00 | 10.40 | 10.79 | 4.86 | 22.54 | 25.43 | 22.88 | 42.11 | 25.13 |
| 112 | | 120-o88 | 3 | 67% | -0.2951 | -45.6 | -5.5 | -5.2 | -5.1 | -5.1 | -5.1 | -5.0 | -5.0 | -4.8 | -4.8 | 154.54 | 0.00 | 11.76 | 31.47 | 5.72 | 12.24 | 24.21 | 2.01 | 35.76 | 31.37 |
| 113 | | 053-o8o8oo | 6 | 33% | -0.2950 | -51.4 | -6.1 | -5.9 | -5.8 | -5.7 | -5.7 | -5.6 | -5.6 | -5.5 | -5.5 | 174.26 | 0.00 | 8.86 | 33.44 | 30.98 | 18.22 | 16.11 | 16.60 | 16.67 | 33.38 |
| 114 | | 119-o8o | 3 | 33% | -0.2894 | -46.3 | -5.6 | -5.4 | -5.4 | -5.2 | -5.0 | -5.0 | -5.0 | -4.9 | -4.8 | 159.99 | 0.00 | 3.15 | 13.65 | 12.91 | 24.49 | 30.57 | 16.59 | 41.64 | 16.99 |
| 115 | Clausen 1 | 018-8oooo8 | 6 | 33% | -0.2883 | -51.8 | -6.4 | -6.2 | -5.8 | -5.7 | -5.6 | -5.6 | -5.6 | -5.5 | -5.4 | 179.67 | 0.00 | 22.05 | 13.23 | 31.69 | 28.30 | 10.87 | 22.03 | 30.89 | 20.61 |
| 116 | | 073-8oooo | 5 | 20% | -0.2834 | -48.7 | -5.8 | -5.7 | -5.6 | -5.5 | -5.3 | -5.2 | -5.2 | -5.2 | -5.2 | 171.87 | 0.00 | 6.86 | 25.18 | 30.34 | 22.76 | 31.06 | 6.60 | 25.18 | 23.90 |
| 117 | | 086-oo8o8 | 5 | 40% | -0.2790 | -50.0 | -5.7 | -5.7 | -5.7 | -5.6 | -5.5 | -5.5 | -5.5 | -5.4 | -5.4 | 179.22 | 0.00 | 27.05 | 24.91 | 10.77 | 25.22 | 25.40 | 24.69 | 26.61 | 14.57 |
| 118 | | 123-oo | 2 | 0% | -0.2760 | -48.1 | -6.1 | -5.4 | -5.4 | -5.3 | -5.3 | -5.2 | -5.2 | -5.1 | -5.1 | 174.30 | 0.00 | 1.34 | 47.48 | 24.55 | 22.74 | 23.72 | 28.23 | 1.64 | 24.60 |
| 119 | | 117-ooo | 3 | 0% | -0.2724 | -47.0 | -5.5 | -5.4 | -5.3 | -5.2 | -5.2 | -5.2 | -5.1 | -5.1 | -5.0 | 172.54 | 0.00 | 24.15 | 26.32 | 25.95 | 26.37 | 3.84 | 30.81 | 3.69 | 31.42 |
| 120 | | 116-8o8 | 3 | 67% | -0.2702 | -46.6 | -5.6 | -5.2 | -5.2 | -5.2 | -5.1 | -5.1 | -5.1 | -5.1 | -5.0 | 172.45 | 0.00 | 10.12 | 23.02 | 35.26 | 12.68 | 26.05 | 26.12 | 13.35 | 25.85 |
| 121 | | 098-888o | 4 | 75% | -0.2494 | -47.1 | -5.4 | -5.3 | -5.3 | -5.2 | -5.2 | -5.2 | -5.2 | -5.2 | -5.1 | 188.86 | 0.00 | 31.47 | 15.30 | 17.07 | 18.76 | 14.94 | 43.72 | 15.75 | 31.86 |
| 122 | | 118-oo8 | 3 | 33% | -0.2397 | -46.7 | -5.7 | -5.5 | -5.2 | -5.2 | -5.1 | -5.1 | -5.0 | -5.0 | -4.9 | 194.81 | 0.00 | 3.79 | 30.26 | 24.28 | 30.37 | 13.22 | 30.42 | 30.54 | 31.93 |
| 123 | | 121-88 | 2 | 100% | -0.2392 | -48.8 | -6.0 | -5.7 | -5.6 | -5.5 | -5.4 | -5.3 | -5.2 | -5.1 | -5.0 | 204.03 | 0.00 | 47.13 | 12.46 | 22.31 | 47.45 | 12.69 | 24.48 | 12.27 | 25.23 |
| 124 | | 001-888888 | 6 | 100% | -0.2093 | -41.3 | -5.6 | -4.8 | -4.6 | -4.5 | -4.5 | -4.4 | -4.4 | -4.3 | -4.2 | 197.36 | 0.00 | 1.68 | 29.33 | 32.80 | 34.84 | 29.77 | 29.46 | 19.68 | 19.80 |

**Supplemental Table 7: PCR oligonucleotide primers.** For the Level 0 cloning, the extremities that were added were located at the 5’ end of the primer sequence. The light blue bases are identical to the template DNA. The purple bases were added either to separate the ORF from the TagRFP sequence or to prevent an early STOP codon. The red bases are the linkers connecting the level 0 blocks, one black base was added in order to match the pattern of the BsaI cutting site indicted by the underlined bold black bases.

| Primer code | Primer name | Primer sequence (5'-3') | Usage/Utility | Target template | Tm |
|---|---|---|---|---|---|
| BF2 | AT1G29050_For2 | CTTGGCATTGGTGGACTCACAAAG | Genotyping | *tbl38* mutant | 67 |
| BR2 | AT1G29050_Rev2 | CATCGAACTCAACACTTTCGACACTACTCC | Genotyping | *tbl38* mutant | 68,6 |
| T2 | LB1 (SAIL T-DNA) | GCCTTTTCAGAAATGGATAAATAGCCTTGCTTCC | Genotyping | SAIL mutants | 72,2 |
| F10 | pPRX36_GGAG_F | **GGTCTC**CGGAGGGCCCATATAAGTT | Golden Gate lvl / Genotyping 0 | Primers not used for cloning: previously cloned (Francoz et al., 2019b) | 68,8 |
| R10 | pPRX36_CATT_R | **GGTCTC**GCATTTTGGACTCTCAGC | Golden Gate lvl0 | | 67,5 |
| F11 | PRX36CDS_AATG_F | **GGTCTC**CAATGAATACAAAAACGGT | Golden Gate lvl0 | *PRX36* cDNA (RIKEN pda20378) | 63,4 |
| R11 | PRX36CDS_TAGC_R | **GGTCTC**GTAGCAACATCATGGTTAA | Golden Gate lvl0 | | 63,4 |
| F12 | TagRFP_GCTA_F | **GGTCTC**CGCTACCGGTATGGTGAG | Golden Gate lvl0 | TagRFP-AS-N entry clone (Evrogen FP149) | 68,8 |
| R12 | TagRFP_AAGC_R | **GGTCTC**GAAGCAAGAAAGCTGGGT | Golden Gate lvl0 | | 68 |
| F15 | TBL38CDS_AATG_F | **GGTCTC**CAATGATGGGTTTCAAACT | Golden Gate lvl0 | *TBL38* cDNA (RIKEN pda09004) | 66,3 |
| R15 | TBL38CDS_TAGC_R | **GGTCTC**CTAGCCATCGTAAGAGCTG | Golden Gate lvl0 | | 65,6 |
| F25 | M13_FOR | GTAAAACGACGGCCAGT | Lv0 sequencing | pGEM-T easy / pCR ZeroBlunt vectors | 58,8 |
| R25 | M13_REV | CAGGAAACAGCTATGACCA | Lv0 sequencing | | 58,7 |
| F31 | p35S CaMV_GGAG_F | **GGTCTC**CGGAGCCAGTGAATTGT | Golden Gate lvl0 / Genotyping | pEAQ-HT-DEST1<br>Primers not used for cloning: Genscript synthesis pUC57-*p35S* (Genscript) | 71,3 |
| R31 | 5'UTR CPMV_CATT_R | **GGTCTC**GCATTATCGAATTTGGG | Golden Gate lvl0 / Genotyping | | 69 |
| F33* | pPRX36_3'_F | CATCCAACAAATTAAAAGCC | Lv1 sequencing | pL1V-R2 recombinant vector | 58,6 |
| F34* | pTBL38_3'_F | TTAATTTATGTGTGTGTGTGTCAG | Lv1 sequencing | pL1V-R2 recombinant vector | 59 |
| F35* | 5'UTR CPMV_3'_F | TACTTCTGCTTGACGAGGTATTGTT | Lv1 sequencing | pL1V-R2 recombinant vector | 63,8 |
| R36* | TagRFP_N_R | GTACACTTAAAGTGGTGATTATTGACAG | Lv1 sequencing | pL1V-R2 recombinant vector | 61,7 |
| F37* | TagRFP_C_F | AAGCAGACAAAGAAACTTACGTGGA | Lv1 sequencing | pL1V-R2 recombinant vector | 65,8 |
| R40* | T-Nos_5'_R | ATTATATGATAATCATCGCAAGAC | Lv1 sequencing | pL1V-R2 recombinant vector | 57,7 |

| | | | | | |
|---|---|---|---|---|---|
| F50 | pL1V-R2_RBshort_F | GCCAATATATCCTGTCAAACACTG | Genotyping / Sequencing | pL1V-R2 / pL2V-HYG/KAN recombinant vectors | 63,2 |
| R51 | pL1V-R2_LB_R | TAGACAACTTAATAACACATTGCGGAC | Golden Gate lvl1 | pL1V-R2 recombinant vector | 64,7 |
| F52 | pL2V-3'p35S_F | TTGGAGTAGACCAGAGTGTCGTG | Golden Gate lvl2 | pL2V-HYG/KAN recombinant vector | 65,4 |
| F61 | TBL38_RT_OUT_F | AAAACCCAAAACCTTTGATATTTGTCTAA | RT-PCR | *TBL38* CDNA | 66 |
| R61 | TBL38_RT_OUT_R | CACTAGAAGATTTACGAATGTCATTTACAGA | RT-PCR | | 65,2 |
| F65 | ACT-2_RT_OUT_F | GGTAACATTGTGCTCAGTGGTGG | RT-PCR | *ACT2* CDNA | 67,8 |
| R65 | ACT-2_RT_OUT_R | CTCGGCCTTGGAGATCCAC | RT-PCR | | 67,5 |

**Supplemental Table 8: Histochemical dyes, primary and secondary antibodies.** A488, Alexa fluor 488; AP, alkaline phosphatase; IF, Immunofluorescence.

| Identifier | Type | Target/epitope | Use | Source |
|---|---|---|---|---|
| Ruthenium Red | Dye | Negatively charged polysacharides | Mucilage release assays | Schuchardt Müchen |
| JIM7 | Rat IgA primary antibody | Highly ME HG | IF / ImmunoDotblot | Kerafast ELD005 |
| LM20 | Rat IgM primary antibody | Highly ME HG | IF / ImmunoDotblot | Kerafast ELD003 |
| αPRX36 | Rabbit polyclonal primary antibody | CSLIGSMENIPSPES PRX36 peptide-KLH conjugate | IF | Genscript private order (Francoz et al., 2019b) |
| αTagRFP | Rabbit polyclonal primary antibody | TagRFP | Western-blot | Invitrogen R10367 |
| Goat αRat IgG-A488 | secondary antibody | Rat primary antibody | IF | Invitrogen A-11006 |
| Goat αRat IgG-AP | secondary antibody | Rat primary antibody | ImmunoDotblot | SIGMA A8438 |
| Goat αRabbit IgG-A488 | secondary antibody | Rabbit primary antibody | IF | Invitrogen A11034 |
| Goat αRabbit IgG-AP | secondary antibody | Rabbit primary antibody | Western-blot | SIGMA A3687 |
| αDig Fab-AP | Anti-Digoxigenin secondary antibody | Dig labeled riboprobes | *In situ* Hybridization | Roche 11 093 274 90 |

**Supplemental Table 9: cDNA and custom-ordered plasmids used as template DNA for further cloning.**

| Sequence identifier | Sequence type | Source | Supplier |
|---|---|---|---|
| *PRX36 ORF* | ORF of *AT3G50990* | RAFL22-03-B11 (pda20378) | RIKEN BioResource Center (https://www.brc.riken.jp) (Seki et al., 1998; Seki et al., 2002) |
| *TBL38 ORF* | ORF of *AT1G29050* | RAFL09-90-M11 (pda09004) | |
| *TagRFP* | ORF of *TagRFP* (codon use optimized for Arabidopsis and Saccharomyces) | Gateway TagRFP-AS-N entry clone (FP149) | Evrogen (https://evrogen.com/) |
| *pPRX36* | Promoter of *AT3G50990* | Sequence synthesis | GenScript (https://www.genscript.com/) |
| *p35S_short* | Promoter of CaMV and 5'UTR of CPMV | pEAQ-HT-DEST1 GQ497235 (5906-6769)<br>Sequence synthesis[a] | (Sainsbury et al., 2009)<br>GenScript (https://www.genscript.com/) |
| T-Nos | 3'UTR, polyadenylation signal/NOS terminator (*A. tumefaciens*) | pICH41421 | (Engler et al., 2014)<br>Gift from P.M. Delaux (LRSV, Auzeville, France) |

[a] The sequence was adapted for easier Golden Gate cloning by adding GGTCTCCGGAG upstream of promoter and AATGCGAGACC downstream of 5'UTR as 5' and 3' security margin, respectively, and by removal of two BbsI site before synthesis (6530: A to C; 6549: T to C). The sequence was provided in pUC57 by https://www.genscript.com/.

**Supplemental Table 10: Level 0 Golden Gate generated constructs.** each block was individually made or ordered and further used to build the Level 1 block assembly (**Supplemental Table 11**). The template used are detailed in **Supplemental Table 9**.

| Lvl 0 blocks | Identifiers | Forward primer (**Supplemental Table 7**) | Reverse primer (**Supplemental Table 7**) |
|---|---|---|---|
| *pPRX36* | 1 | **F10 (primer not used, previously cloned (Francoz et al., 2019b))** | **R10 (primer not used, previously cloned (Francoz et al., 2019b))** |
| *p35S* | 3 | **F31 (primer not used, genescript synthesis)** | **R31 (primer not used, genescript synthesis)** |
| *PRX36* CDS | 4 | **F11** | **R11** |
| *TBL38* CDS | 5 | **F15** | **R15** |
| *Tag-RFP* | 20 | **F12** | **R12** |
| *T-Nos* | 21 | Gift from P.M. Delaux (LRSV, Auzeville, France) | |
| *TBL38* CDS stop | 27 | **F15** | **R75** |

**Supplemental Table 11: Level 1 in pL1V-R2 vector and finalized Level 2 Golden Gate generated constructs.** Selected level 0 blocks (**Supplemental Table 10**) were mixed together to produce different constructs used in stable or transient transformations. Level 1 plasmid backbone was pICH47811 “pL1V-R2”pL1V-R2 vector (Weber et al., 2011). Level 2 plasmid backbone was EC15027 “pL2V-HYG” with pICH471744 “pL1M-ELE-2” as a linker-containing plasmid (gift from P.M. Delaux, LRSV, Auzeville-Tolosane, France).

| Finalized constructs | Lvl 0 combination | Lvl 1& 2 Identifiers | Lvl 2 Resistance |
|---|---|---|---|
| *p35S ::TBL38-TagRFP T-Nos* | 3-5-20-21 | 100 | Hygromycin |
| *pPRX36 ::TBL38-TagRFP T-Nos* | 1-5-20-21 | 107 | Hygromycin |
| *pPRX36 ::PRX36-TagRFP T-Nos* | 1-4-20-21 | 108 | Hygromycin |

### Supplemental methods

#### Ruthenium red mucilage release test, image analysis and statistical analysis

The high-throughput adherent mucilage release semi-quantitative phenotyping used the ruthenium red (**Supplemental Table 8**) staining method previously described allowing to calculate adherent mucilage area and circularity (Francoz et al., 2019b). Briefly, following staining, seeds were transferred in 12-well microplates and scanned at 6400 dpi with an Epson Perfection V700 photo scanner. Images were analyzed using ImageJ 1.8 ([https://imagej.nih.gov/ij/](https://imagej.nih.gov/ij/)) without edition of native images with an updated ImageJ script described below that facilitates cleaning of the data and allows automatic storage of the data. Seed area (without mucilage) and whole seed area (seed + mucilage) were manually edged in a semi-automatic fashion. Non-individualized seeds were automatically removed from the analysis and selected parameters were measured. A manual examination allowed to double check the absence of artifactual data. The cleaned data were extracted and pooled into a Microsoft Excel sheet for further analyses. At least two technical repeats, each with about 50-100 seeds were performed for seed batches coming from 6 individual Col-0 and *tbl38* plants simultaneously grown in the same conditions. Results are presented as mean ±SD with n > 1,000 seeds. Statistical data were obtained using ANOVA tests with n ≥ 3.

#### ImageJ script for Ruthenium red staining semi quantitative analysis

```
run("Clear Results");

roiManager("Reset");

roiManager("UseNames", "true");

run("Set Measurements...", "area mean perimeter shape display redirect=None decimal=3");


setForegroundColor(255, 255, 0);

run("Line Width...", "line=1");

roiManager("UseNames", "true");
```

```
title=getTitle();
nom=File.nameWithoutExtension();
dir=File.directory;

run("Duplicate...", " ");
title2=getTitle();
selectWindow(title);

run("Properties...", "channels=1 slices=1 frames=1 unit=um pixel_width=4.1344 pixel_height=4.1344 voxel_depth=4.1344 global");
//run("Subtract Background...", "rolling=100 light separate sliding");
run("Split Channels");
selectWindow(title+" (blue)");
close();

//threshold graines ALL (graine + mucillage indiv

selectWindow(title+" (green)");
run("Threshold...");
waitForUser("graines entiere","mettre en rouge les graines entieres");
setOption("BlackBackground", false);
run("Convert to Mask");
run("Watershed");
run("Set Measurements...", "area shape display redirect=None decimal=3");
```

```
run("Analyze Particles...", "size=60000-Infinity circularity=0.7-1.00 show=Masks exclude add");
//run("Analyze Particles...", "size=200000-Infinity show=Masks exclude add");


c=roiManager("Count");



for(i=0;i<c;i++)
{
roiManager("Select", i);
roiManager("Rename","graine ALL"+i+1);
}



//threshold graines seules et add manager =graines seules global


selectWindow(title+" (red)");


run("Threshold...");
waitForUser("centre des graines","mettre en rouge le centre des graines");
setOption("BlackBackground", false);
run("Create Selection");
roiManager("Add");
roiManager("Select", c);
roiManager("Rename", "graines noires global");
```

```
for(i=0;i<c;i++)

{

roiManager("Select", newArray(i,c));

roiManager("AND");

roiManager("Add");

roiManager("Select", c+i+1);

roiManager("Rename", "graine"+i+1);

}


roiManager("Show All");

roiManager("Measure");

selectWindow(title2);

roiManager("Show All");


waitForUser("trier les ROI","trier le ROI");

run("Clear Results");

roiManager("Show All");

roiManager("Measure");

saveAs("Results", dir+nom);

roiManager("draw");

saveAs("jpeg", dir+nom+"_DRAW");

roiManager("save", dir+nom+".zip");
```

### *A. thaliana* silique fixation, paraffin tissue array embedding and microtomy

The protocol was as previously described (Francoz et al., 2019b), with minor improvement using biopsy foam enabling handling more samples. Briefly, for each genotype, flowers were marked on the developing floral stem at the petal emergence stage used as a proxy to define the pollination time. At the end of floral stem development, sequential kinetics of developing siliques of each genotype were systematically harvested, placed in order (pedicel on top) and sandwiched in Deltalab histoset 2 embedding cassettes (Dutscher 039751) between two 30,2 × 25,4 × 2 mm SimPort™ biopsy foams (Dutscher 040666). Selected marked siliques that were initially labeled to account for a given day after pollination (DAP) were placed with pedicels facing down for better identification. Shortly after, cassettes were submerged in FAA fixative (3.7 % (v/v) formaldehyde from 37 % formaldehyde solution (Sigma 1.04002); 5 % (v/v) acetic acid; 50 % (v/v) ethanol; 35 % (v/v) Milli-Q water or RNase-free DEPC-treated water, depending on the necessity), vacuum infiltrated for 8 x 1 min and fixed at 4°C for 6-8 h. Cassettes were then washed 4 times in 50 % ethanol and placed at 4°C for 72-84 h. Sample dehydration and paraffin embedding were done as previously described (Francoz et al., 2019a; Francoz et al., 2019b; Francoz et al., 2016). The ordered siliques containing seed development kinetics were used to assemble organized tissue arrays each encompassing up to 1000 developing seeds (Francoz et al., 2019a; Francoz et al., 2019b) To assemble paraffin blocs, cassettes were delicately opened on a slide warmer set to 60°C (LabScientific XH-2001) and siliques were deposited on metallic mold with respect to the initial order. The molds were covered with a fitting plastic ring (Simport M460), the paraffin tissue-array blocks were allowed to solidify at least overnight at 4°C and could be stored for months/years. Ten-to 12 µm-thick serial sections were made with a rotary microtome and spread on silane coated slides.

### *In situ* RNA hybridization

*In situ* RNA hybridization experiments were performed as described (Francoz et al., 2019a; Francoz et al., 2016). In short, a plasmid (RAFL09-90-M11, pda09004) containing *TBL38* full length cDNA including 5'-UTR and 3'-UTR was ordered at RIKEN BioResource Center (https://www.brc.riken.jp) (Seki et al., 1998; Seki et al., 2002) to be used as a template for riboprobe synthesis. The plasmid was first linearized with EcoRI at the 5' end of the *TBL38* ORF and with BamHI at the 3' end to further generate anti-sense and sense probe used as a negative control, respectively. The synthesis of digoxygenin-labeled riboprobes was performed by *in vitro* transcription using DigRNA labeling mix (Roche 11277073910) and T3 (Promega P2083) or T7 RNA polymerase (Promega P2075), respectively. Serial sections of paraffin-embedded wild-type seed development kinetics encompassing all developmental stages were first deparaffinized with xylene and rehydrated with an inverted ethanol series up to water. Each probe was hybridized overnight on paraffin serial sections of seeds. Following stringent washing steps, detection of hybridized riboprobes was performed with the anti-digoxigenin-AP conjugates (**Supplemental Table 8**) revealed with its chromogenic substrate (BCIP/NBT) overnight. Finally, slides were mounted in EUKITT® (Dutscher 045799) and scanned with the nanozoomer HT (Hamamatsu, https://www.hamamatsu.com/). Image analysis was performed using NDPview (Hamamatsu, https://www.hamamatsu.com/) and figures were assembled using Corel Photopaint.

**Immunofluorescence on tissue-array sections**

We used our previously described protocol (Francoz et al., 2019b). Briefly, serial sections of paraffin-embedded Col-0 or *t bl38*mutant seed development kinetics were first deparaffinized with xylene and rehydrated with an inverted ethanol series up to water. The slides were placed in a 20-slide plastic rack and collectively blocked in 200 mL 5 % TTBS-milk (5 % non-fat dry milk (Regilait), 0.01 M Tris-HCl pH 7.5, 0.5 M NaCl, 0.3 % Tween 20) for 30 min at RT. Slides containing Col-0 and *tbl38* samples were separately incubated with 150 µL of anti-PRX36 primary antibodies (**Supplemental Table 8**) diluted 1:10 dilution in TTBS-milk, for a 3-4 h under a coverslip in a home-made humid chamber. The coverslip was carefully removed by dipping the slide in TTBS-milk and the slides were collectively washed 6 x 5 min in 200 mL TTBS. 150 µL of goat-anti rabbit-A488 secondary antibodies (**Supplemental Table 8**) of diluted 1:100 in TTBS-milk was incubated for 1-2 h at RT. The coverslips were carefully removed and the slides were collectively washed 3 x 5 min in 200 mL TTBS and 3 x 1 min in Milli-Q water. Slides were mounted in Prolong Gold antifade (Molecular probes P36934) or Fluoromount-G™ (Invitrogen 15586276). Serial section of Col-0, *tbl38*, PTR2.4 and PTR3.5 developmental kinetics were similarly labeled with JIM7 or LM20 primary antibodies followed by Goat anti-rat A488 secondary antibodies (**Supplemental Table 8**). Slides were scanned using a Nanozoomer 2.0RS scanner (Hamamatsu) at 20 × or 40 ×. The FITC (excitation: 482/18 nm; dichroic mirror 488 nm; emission: 525/30 nm) filter sets and the bright field (BF) mode were sequentially used to visualize Alexa488 fluorescence and the morphology, respectively. Scans were analyzed using NDP view (Hamamatsu, Hamamatsu City, Japan).

**Confocal spinning disk microscopy of seed development kinetics of *A. thaliana* stable transformants (high throughput strategy)**

Developing siliques taken from plants expressing the different TagRFP constructs in various genetic backgrounds were dissected and the replums containing the seeds were mounted under a coverslip in distilled water. Image were taken with a PLAN APO 20x/0.75 dry objective using the confocal spinning disk microscope from Perkin Elmer driven by the Volocity 6.3.0 software and equipped with a YokogamaCSU-X1 scan head, two EmCCD Hamamatsu C9100-13 cameras (Hamamatsu) and a 580 nm beam splitter to separate dual staining on the two cameras as described (Francoz et al., 2019b). Images were acquired for TagRFP fluorescence with a 561 nm laser (Laser power intensity: 7% or 15%, exposure time: 200 msec; gain: 5; sensitivity: 148) and the fluorescence was selected between 580 and 612 nm. Image J was used to analyze the Z distribution of the fluorescence intensity profile along regions of interest (ROIs). The set display range was set to 16-bit (0-65635) for the 16-bit spinning disk images that were further calibrated using the set scale option (1 pixel = 0.66 µm). Three modes were used: (i) the maximum intensity Z projection mode was used to build a stack image of a given number of 1 µm-slices taking the maximum fluorescence value among the slices for each pixel; (ii) the sum intensity Z projection mode was used for fair comparison of relative fluorescence intensities among the analyzed lines since all the intensity values from all selected slices were summed for each pixel; (iii) the orthogonal view along an axis positioned in the XY plan was used to better visualize the Z distribution of the fluorescence along the chosen XY axis.

**SDS-PAGE and anti-TagRFP western-blots from PTR complemented lines**

The TagRFP fluorescence patterns of PTR lines was controlled using spinning disk confocal microscopy along developmental kinetics of seeds from staged floral stem. The seeds that displayed the early (6-8 DAP), medium (8-12 DAP) and late (> 12 DAP) fluorescence patterns were carefully extracted from the dissected siliques (3-4 siliques per fluorescence pattern) and each pooled in 2 mL tubes with a metallic grinball before freezing in liquid nitrogen. Total proteins were extracted and analyzed by western blot as previously described (Francoz et al., 2019b) using anti-TagRFP primary and anti-rabbit-AP secondary antibodies (**Supplemental Table 8**) with minor modifications:

For SDS-PAGE analysis, total proteins were extracted as follows: frozen weighed samples were grinded using a Mixer Mill MM 400 (RETSCH). with a metallic grindball twice 30 s at 30 Hz. Eight µL of 5 mM sodium acetate pH 4.6, 0.2 M $CaCl_2$ extemporaneously complemented with 1 $\mu L.mL^{-1}$ of β-mercaptoethanol and 1.5 $\mu L.mL^{-1}$ of plant protease inhibitor cocktail (Sigma P9599) were added per mg of seed and the tubes were plunged in liquid nitrogen. The samples were further grinded in the frozen buffer 3 times 30 s at 30 Hz. The samples were spinned and thoroughly vortexed. Then, they were vertically agitated at 400 rpm on an orbital shaker at 4°C for 45 min. After 10 min of centrifugation at 14,000 rpm, the supernatant was discarded (controls showed that no protein of interest was present in this fraction). The remaining pellet was mixed with 8 µL of denaturation solution containing 10 % (v/v) β-mercaptoethanol, 6 M urea, 10 % (v/v) glycerol, 5 % (v/v) SDS (Kato et al., 2002) complemented with 0.01 % bromophenol blue and additional 0.2 M $CaCl_2$ per mg of initial seeds. The samples were run for a second vertical extraction again at 400 rpm on an orbital shaker at 4°C for 45 min. After 10 min of centrifugation at 14,000 rpm, the supernatant was carefully transferred in a clean 1.5 mL tube. Twenty-five µL of each sample were analyzed by SDS-PAGE in a 13 % or 8% resolving and 4 % polyacrylamide denaturing stacking gels under electric current set at 120 mA and 500 V.

Protein were transferred onto nitrocellulose membranes for 45 min to 1 h under electric current set at 20 V and 800 mA. Membranes were transiently stained with Ponceau Red solution and imaged. Nitrocellulose membranes were blocked overnight in 0.02 M Tris-HCl pH 7.5, 0.5 M NaCl, 0.5 $g.L^{-1}$ Tween 20 and 5 % Régilait non-fat milk (TTBS-milk) under constant agitation at 4°C and 60 rpm on an orbital shaker. Membranes were incubated at RT for 1 h 30 min and then overnight at 4°C in 10 mL of anti-TagRFP primary antibody (**Supplemental Table 8**) diluted 1:3,000 in TTBS-milk, then rinsed 6 x 5 min with TTBS. Secondary antibody incubation was performed for 2 h in 10 mL of goat anti-rat AP secondary antibody (**Supplemental Table 8**) diluted 1:5000 in TTBS-milk at room temperature. Membranes were rinsed 3 x 5 min with TTBS and once with deionized water before revelation in 50 mL of alkaline phosphatase BCIP/NBT chromogenic substrate for 30 min. Membranes were imaged with a Canon EOS 550D digital camera.

**Dry seed MSC surface wall abrasive fraction**

Abrasion columns were a home-made design allowing for homogenous dry seed MSC surface wall abrasive enrichment using plastic column and collector tube from the GeneJET Plasmid Miniprep Kit (Thermo Scientific K0502). The bottom of the column was cut with a razor blade, the silica layer of the column was removed and replaced with a 50 µm Seffar Nylon mesh (Dutscher 074010) wrapped around the bottom of the open column. Then, a 2.1 x 1.6 cm piece of P500 sand-paper (Dexter, Castorama) was gently rolled in the column and placed with the abrasive side inward. The P500 sand paper theoretically corresponded to 30.2 µm average abrasive particle size (https://www.fine-tools.com/G10019.html), *i.e.* about 1:10 of *A. thaliana* seed length) Finally, the so-called abrasive column was put back in the collector tube. The collector tube was weighted at the time of use and 50 mg dry seeds (about 2,500 seeds) were deposited inside the paper sand roll before locking the cap. Abrasion was done with a FastPrep (MP Biomedicals™ 116004500) for 5 to 6 cycles of 1 min at 6.5 $m.s^{-1}$. The abrasion was not linear and the powder usually appeared in the collection tube after the $4^{th}$ or $5^{th}$ cycle probably corresponding to a breaking limit. While most of the extracted surface wall powder passes through the nylon mesh during the last cycles of abrasion, a final centrifugation step for 5 min at 8,500 rpm allowed total recovery of the powder. The powder was carefully weighted in the collector tube (in the mg range per 50 mg dry seeds) and additionally grinded with a metallic grindball for 3 min 30 s at 30 Hz (Mixer Mill MM 400, RETSCH). To evaluate the abrasion efficiency, Col-0 dry seeds were analyzed through four different methods for comparison before and after abrasion. (i) Ruthenium red mucilage release assay using the protocol described above: seeds are shaken at 250 rpm in Tris Buffer (0.01 M Tris-HCl pH 7.5) during 1 h at RT, rinsed with Tris Buffer, and shaked at 250 rpm in a 0.02 % Ruthenium Red solution in Tris Buffer during 1 h. Following 2 Tris washing steps, seeds are transferred in a 24-well microscopy plate and imaged. (ii) UV autofluorescence of dry seed surface: Seeds were imaged by epifluorescence using an UV filter set (excitation: 387/11 nm; dichroic mirror: 405 nm; emission: 440/40 nm) using a Leica DM IRB/E inverted microscope equipped with a Leica MC190HD digital camera. (iii) Fixation and embedding in LRW acrylic resin (Francoz et al., 2019b), microtomy and immunofluorescence on 1 µm semi-thin sections using a previously described protocol (Oudin et al., 2007) with JIM7 and goat anti-rat A488 antibodies (**Supplemental Table 8**). (iv) Finally, abraded and non-abraded seeds, 100 µm nylon mesh and the abrasion powder were imaged by scanning electron microscopy. Samples were metalized using a MED 020 modular high vacuum coating and directly observed with no further treatment using a Quanta 250 FEG FEI electron microscopy. Images were taken with a 2.0 spot size and 10.00 kV acceleration. Following these setup controls, powder from various genotypes was analyzed by immuno dotblots, enzymatic assays and mass spectrometry. > 10 seeds were observed with similar results.

**Immuno dotblot and semi-quantitative analysis from dry seed MSC abrasive fractions**

Powder of MSC surface CW was chemically extracted in the collector tube with 80 µL of extraction buffer [80 µL of 5 mM sodium acetate pH 4.6 containing 1 $\mu L.mL^{-1}$ of plant protease inhibitor cocktail (Sigma P9599)] per mg of powder. The collector tube with cap was shaken at 250 rpm during 1 h at 4°C to ensure proper extraction. After a 5 min centrifugation at 14,400 rpm, the supernatant was recovered, diluted (1 µL in 50 µL milliQ-$H_2O$) and deposited into sample wells onto nitrocellulose membrane disposed into a 96-well Bio-Dot microfiltration apparatus (BioRad) previously soaked with in TBS (0.02 M Tris-HCl pH 7.5, 0.15 M NaCl) for 10 min. A 1 min 30 s of vacuum insured homogeneous transfer among the wells. The membrane was briefly air-dried to ensure proper adsorption and blocked in 5 % milk (Regilait® non fat dry milk) in TTBS (0.05 % Tween 20, 0.02 M Tris-HCl pH 7.5, 0.5 M NaCl) overnight at 4°C, under a 50 rpm gentle shaking. The membranes were rolled in a hemolysis tube containing 5 mL of JIM7 or LM20 primary antibodies (**Supplemental Table 8**) diluted 1:100 in TTBS-milk for 3-4 h at RT. Membranes were washed 6 x 5 min each in about 50 mL TTBS before incubation for 1 h in a hemolysis tube filled with 5 mL of 1:5000 alkaline phosphatase (AP) conjugated secondary antibody adapted to the species used to raise the primary antibody (**Supplemental Table 8**). After 3 x 5 min wash in about 50 mL TTBS and 3 x 1 min in deionized water, membranes were immersed in the alkaline phosphatase (AP) chromogenic substrate 0.3 $mg.mL^{-1}$ NBT 0.15 $mg.mL^{-1}$ BCIP in 100 mM Tris-HCl pH 9.5, 100 mM NaCl, 10 mM $MgCl_2$ for 5 to 15 min. The membranes were air dried and imaged with a Canon EOS 550D digital camera. Dot blots were analyzed using ImageJ 1.8 without edition of native images. Multiple ROIs of a similar selected size were used to measure the mean intensity of the signal. Two negative controls were used (i) the extraction buffer that was blotted on the membrane and (ii) for each lane, another ROI placed outside of the dots to account for possible vertically different background noises. Measurements were extracted on an Excel file ready for R statistical analyses. Corresponding Col-0 values were assigned as “100 % of signal” and percentage of mutant signal values were calculated accordingly ± standard deviation (SD). Statistical data was obtained using ANOVA / Tukey HSD tests with $n \geq 3$.

**Acetyl group deesterification and acetic acid semi-quantitative analysis from dry seed MSC abrasive fractions**

The acetylation of cell wall polymer in MSC surface wall abrasive samples was analyzed directly in the collection tube following adaptation of previously described method (Stranne et al., 2018). Acetyl groups were released by ester alkaline hydrolysis with 1 M NaOH (80 µL of NaOH per mg powder). The samples were vigorously shaken horizontally for 1 h at 250 rpm at 4°C. The reaction was stopped with an equal volume of 1 M HCl which stabilizes pH at a neutral value. Acetic acid content was then determined using the K-ACET enzymatic Acetic Acid Assay Kit (Megazyme) by measuring the stoichiometric conversion of acetic acid to NADH following three enzymatic reactions. The same volume of samples (100 µL) was used following the suggested protocol with the following adaptation (1.5 mL of final reaction volume was used instead of 2.84 mL). NADH was measured at 340 nm with a ©Agilent Cary 60 UV-Vis spectrophotometer using the Eppendorf Uvette (Dutscher 033189). The linearity of the assay was checked with acetic acid standard provided with the kit and with an acetic acid standard curve (0-5µg). The final results were expressed as mean µg of acetic acid released per mg of original powder ± standard deviation (SD) with $n \geq 3$. Statistical data was obtained using ANOVA / Tukey HSD tests.

**Production of recombinant TBL38 in *Pichia pastoris* and enzymatic activity assays**

The coding sequence of TBL38 (Q8VY22) was codon optimized for *Pichia pastoris* and synthesized without signal peptide in frame with His-tag in pPCIZ-αB by ProteoGenix (Schiltigheim, France). rTBL38 was produced in *P. pastoris* following the previously described protocol (Lemaire et al., 2020). rTBL38 purification was carried out using 1 mL HisTrap excel column (GE Healthcare, Chicago, Illinois, United State). 100 mL of the culture supernatant was loaded onto the column at 1 mL/min flow rate. Column was washed with 10 column volumes of wash buffer (50 mM NaP pH 7.2, 250 mM NaCl, 5 mM imidazole). rTBL38 was purified using 10 column volumes of elution buffer (50 mM NaP pH 7.2, 250 mM NaCl, 100 mM imidazole). The elution fraction of rTBL38 was concentrated with Amicon Ultra Centrifugal filter with a 10 kDa cut-off (Merck Millipore, Burlington, Massachusetts, United States) up to a volume between ~150 µl. The buffer was exchanged to the activity buffer (Mcilvaine's 50 mM pH 6.5, 100 mM NaCl) using PD SpinTrap G-25 (Cytiva, Björkgatan, Uppsala, Sweden). Enzyme purity and molecular weight were estimated by 12 % SDS-PAGE using mini-PROTEAN 3 system (BioRad, Hercules, California, United States). Gels were stained using

PageBlue Protein Staining Solution (Thermo Fisher Scientific) according to the manufacturer's protocol. Activity assays of rTBL38 was performed with the acetic acid assay kit (K-ACETRM, Megazyme) using the activity buffer at 40°C and with three different acetylated substrates (Triacetine 100 mM, Xylan 24 % acetylation, 10 $mg.ml^{-1}$ sugar beet pectins 31 % acetylation 10 $mg.ml^{-1}$). Activity of rTBL38 was expressed as nmole acetic acid.µg $protein^{-1}.min^{-1}$ and compared to that of boiled sample.

### *In silico* model and docking simulations

Homology models for PRX36 and TBL38 (UniProt accession number Q9SD46 and Q8VY22, respectively) were built with the Phyre2 server (Kelley et al., 2015). For PRX36, the template was the crystallographic structure (X-Ray diffraction, 1.45 Å) of *A. thaliana* PRX53 (At5g06720; Protein Data Bank no.1PA2) (Ostergaard et al., 2000). For TBL38, the template was the crystallographic structure (X-Ray diffraction, 1.85 Å) of *A. thaliana* TBL29/ESK1/XOAT1 (Lunin et al., 2020). For comparison with the TBL38 model, the TBL29/ESK1/XOAT1 structure was drawn as well using the 6cci pdb file (Lunin et al., 2020) visualized and analyzed with Swiss-PdbViewer (http://www.expasy.org/spdbv/) (Guex and Peitsch, 1997).

For PRX36 docking experiments, α-D-(1-4) polygalacturonic acid structural model (Braccini et al., 1999) was retrieved from the Glyco3D portal (Perez et al., 2015) http://glyco3d.cermav.cnrs.fr/mol.php?type=polysaccharide&molecule=2504) and was modified to initially build the five hexagalacturonates models used to establish JIM7 specificity (Clausen et al., 2003). This selection was extended to the 124 oligogalacturonates (OGAs) from DP2 to DP6 covering all the theoretical combination of methylation (64 OGAs of DP6 + 32 OGAs of DP5 + 16 OGAs of DP4 + 8 OGAs of DP3 +4 OGAs of DP2). AutoDock Tools (Morris et al., 2009) and AutoDock Vina (Trott and Olson, 2010) were used for simulating the binding of the 124 OGAs to PRX36 within a search box encompassing the whole target protein enabling to recover the 9 best poses for each OGA. Energy level and Root-mean-square deviation (RMSD) upper bound values were obtained with AutoDock tools, representing the affinity of each pose and distances from pose 1 for a given OGA, respectively. RMSD upper bound matches each atom of a given OGA in each conformations (poses) with itself in the conformation of reference (pose 1). As an imperfect mean to sort and rank the 124 OGAs using Microsoft Excel, we used the following proxy: We summed the energy levels (negative values) of the nine poses (the lower, the higher affinity) and summed the nine RMSD for each OGA (the lower, the less dispersed). The ratio of both was used as a proxy integrating both parameters (the lower ratio, the higher affinity and higher gathering of the poses on the protein). Structural models were visualized and analyzed with Swiss-PdbViewer (http://www.expasy.org/spdbv/) (Guex and Peitsch, 1997). The visualization of the models and color edition were performed using Swiss-PdbViewer. Finally, an additional docking simulation of PRX36 was made with the two best hits (OGA of DP 6, so-called Clausen 4 and Clausen 3) to which an acetyl was first added *in silico* at *O*-2 or *O*-3 position. The same calculation as above were made and the results for the four new acetylated OGAs (Clausen-3 *O*-2 ac, Clausen 3 *O*-3 ac, Clausen 4 *O*-2 ac and Clausen 4 *O*-3 ac) were integrated in the previous ranking.
